# Engineering Binding Efficiency and Interaction Stability of a Thermostable Cohesin–Dockerin Pair on the Bacterial Cell Surface

**DOI:** 10.64898/2026.08.09.743725

**Authors:** Barbora Jankovičová, Aleksandra Bigos, Bartłomiej Surpeta, Miguel Silva, Jan Brezovský, Pavel Dvořák

## Abstract

Efficient conversion of polymeric feedstocks for sustainable bioprocessing requires robust strategies for enzyme assembly and cell-surface attachment. In nature, cellulosomes achieve highly efficient lignocellulosic polysaccharide deconstruction through scaffoldin-mediated organization of carbohydrate-active enzymes via specific cohesin–dockerin interactions. These modular binding pairs are therefore attractive tools for synthetic biology and engineered whole-cell biocatalysis, yet their performance has been studied mainly *in vitro* or in yeast or Gram-positive bacteria. The factors governing their function on the microbial surfaces - particularly those of Gram-negative bacteria - remain incompletely understood. Here, we investigated the binding efficiency and interaction stability of two thermophilic cohesin–dockerin pairs from *Acetivibrio thermocellus* and *Acetivibrio clariflavus* displayed on the surface of the genome-streamlined strain *Pseudomonas putida* EM371 using an Ag43-based display system from *Escherichia coli* and a dockerin-tagged fluorescent reporter. We show that binding efficiency is strongly affected by the temperature at which the cohesin–dockerin complex is formed. We further demonstrate that the interaction stability of the *A. clariflavus* pair can be substantially improved by targeted amino acid substitutions in the dockerin domain guided by molecular dynamics simulations and free-energy calculations. These results identify key parameters controlling the performance of thermophilic cohesin–dockerin modules on living bacterial cell surfaces and establish a computation-guided strategy for engineering more stable cellulosome-derived assembly interfaces, advancing the development of modular whole-cell platforms for sustainable biotechnology applications.

**TOC graphics:** 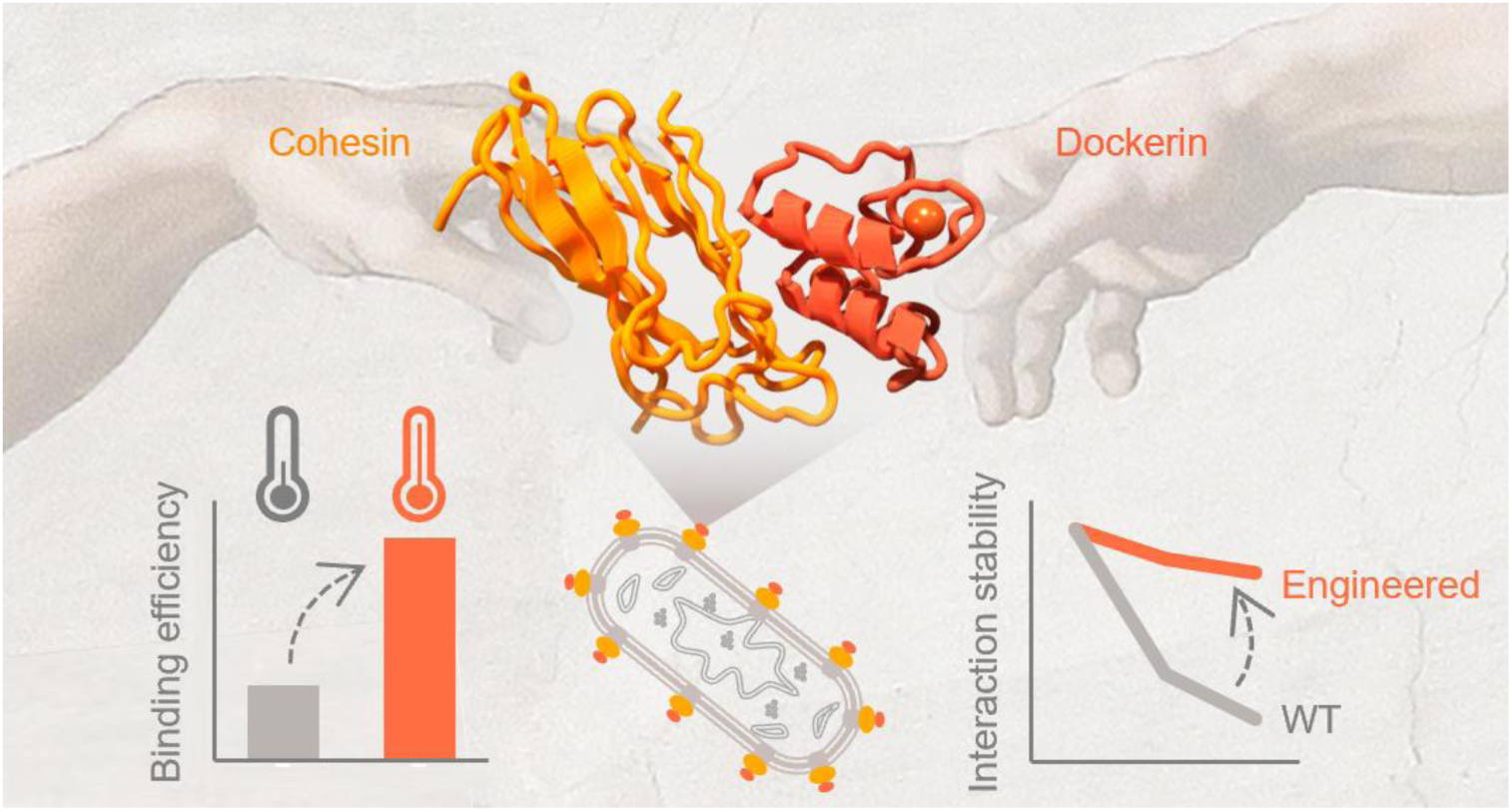

Cohesin–dockerin pairs provide strong and modular non-covalent interactions for synthetic biology and biotechnology applications. We establish an experimental and computational pipeline to improve their two key properties - binding efficiency and interaction stability - on the surface of *Pseudomonas putida*, enabling more robust cell-surface assembly systems.

## Introduction

Lignocellulosic biomass is the most abundant renewable organic material on Earth, and its efficient depolymerization into monomers and oligomers that can be assimilated by microorganisms is central both to global carbon cycling and to the development of sustainable biotechnological processes within a circular bioeconomy. Among the most effective natural lignocellulose degraders are bacteria such as *Acetivibrio thermocellus* and *Acetivibrio cellulolyticus*, which display highly specialized biocatalytic assemblies, termed cellulosomes, on their surface.^1–3^ Cellulosomes are multienzyme complexes organized by scaffoldin proteins that recruit and spatially coordinate multiple carbohydrate-active enzymes (**Fig. 1**). Enzyme attachment to scaffoldins is mediated by specific interactions between type I cohesin and dockerin domains.^4^ Because of their efficiency, modularity, and hierarchical organization, cellulosomes have become an important model for studying the evolution of complex biocatalytic systems.^4–6^ They also represent a valuable source of biological parts for synthetic biology and biotechnology, particularly for the development of engineered whole-cell catalysts and consolidated bioprocessing strategies for lignocellulosic feedstocks.^7–9^

**Figure 1.**
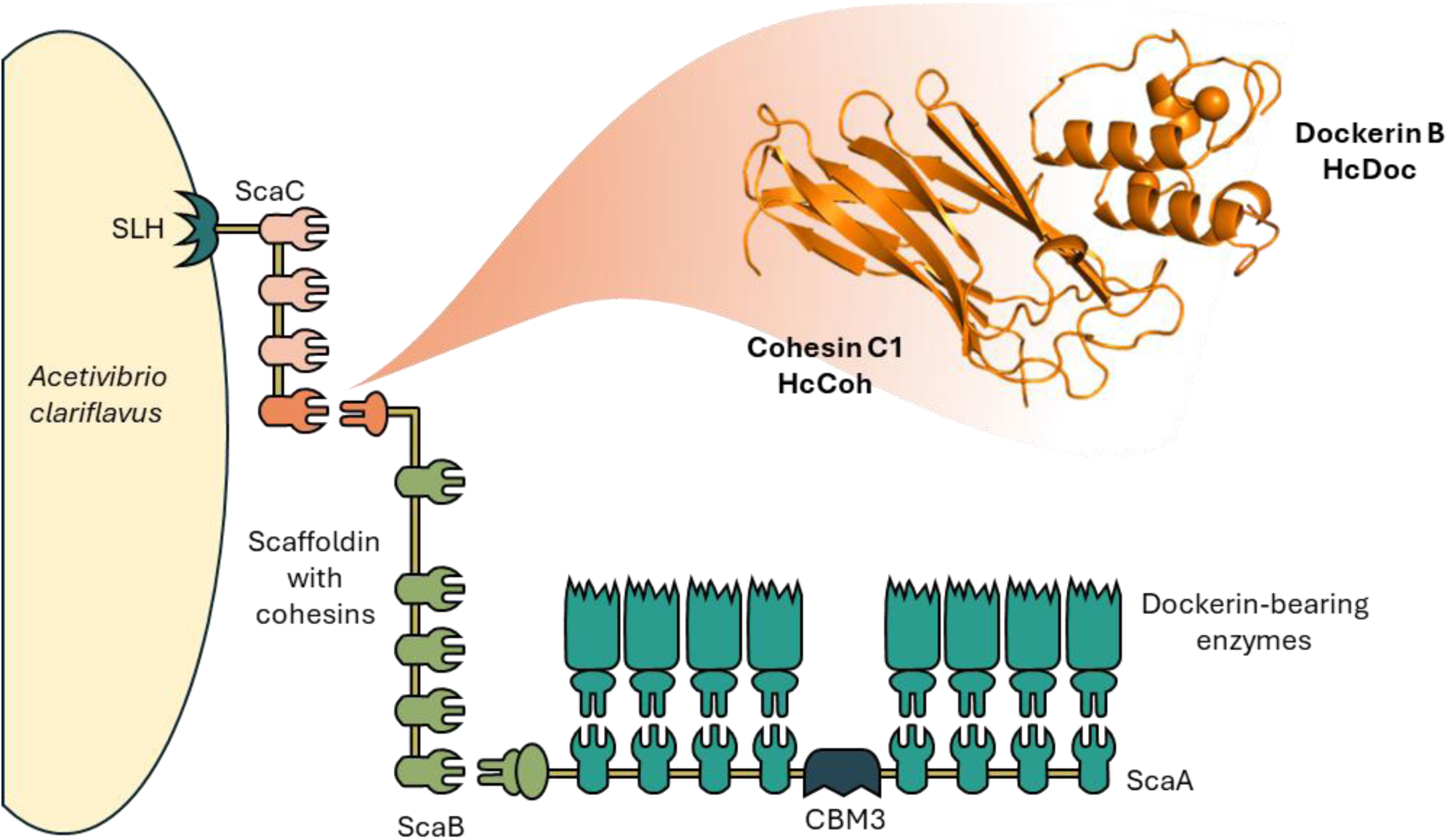
Schematic representation of the cellulosome architecture in *Acetivibrio clariflavus*. (formerly *Hungateiclostridium* or *Clostridium clariflavum*; for the sake of clarity, only a portion of the cellulosome is shown). The diagram illustrates the modular organization of the cellulosome, including scaffoldin proteins containing cohesin modules and enzymatic subunits bearing dockerin domains. The interaction between cohesin and dockerin mediates the assembly of the multi-enzyme complex on the bacterial cell surface. The cohesin–dockerin pair highlighted in orange (CohC1, abbreviated here as HcCoh; DocB, abbreviated here as HcDoc) is in the focus of this study and is shown in detail as a structural model to emphasize its non-covalent molecular interaction. Abbreviations: SLH, surface-level-homology module; CBM3, carbohydrate-binding module; ScaC, anchoring scaffoldin; ScaB, adaptor scaffoldin; ScaA, primary scaffoldin. Adapted and modified from Artzi *et al*. (2014).^34^

Designer cellulosomes, sometimes referred to as minicellulosomes because of their simplified architecture, have been assembled on the surfaces of only a limited number of model hosts, including the yeasts *Saccharomyces cerevisiae*, *Pichia pastoris*, and *Kluyveromyces marxianus*, and the Gram-positive bacteria *Bacillus subtilis*, *Lactococcus lactis*, and *Lactobacillus plantarum*.^10–16^ In contrast, their implementation in Gram-negative hosts remains much less explored. This is a notable limitation, as Gram-negative bacteria offer several advantages for sustainable bioprocessing, including metabolic versatility, environmental robustness, and compatibility with the conversion of heterogeneous feedstocks. We previously investigated the display of designer miniscaffoldins and minicellulosome on the surface of *Pseudomonas putida* KT2440.^17^ This metabolically versatile Gram-negative bacterium is an attractive host for biotechnology and synthetic biology, including lignocellulose valorization, because of its robustness and ability to metabolize a broad spectrum of substrates, including sugars, organic acids, amino acids, or aromatic compounds.^18–23^ We further showed that EM371, a genome-streamlined derivative of wild-type *P. putida* KT2440 lacking many nonessential outer-membrane structures, is a highly suitable host for the surface display of recombinant proteins, including miniscaffoldins, via the Ag43 autotransporter adapted from *Escherichia coli*.^17^

The strong and specific noncovalent interactions between cohesin and dockerin domains are central to cellulosome assembly. With dissociation constants typically in the nano- to subnanomolar range, these interactions appear to have been evolutionarily optimized to ensure robust complex formation under demanding environmental conditions.^24,25^ Their strength, specificity, and modularity have also made cohesin–dockerin pairs attractive tools for biotechnology and synthetic biology. However, most studies of these interactions have been performed *in vitro* using purified proteins and methods such as ELISA, isothermal titration calorimetry, or single-molecule force spectroscopy.^26–29^ Several studies have additionally reported cohesin or dockerin variants with improved affinity or stability, again evaluated primarily *in vitro*.^9,28^ By contrast, systematic *in vivo* analyses remain scarce and are lacking in Gram-negative bacterial hosts.^30,31^ In particular, the factors governing binding efficiency and interaction stability after heterologous expression, surface display, and protein attachment on living cells remain poorly defined.

In our previous work, we focused on identifying an optimal *P. putida* host background and display system to maximize the abundance of surface-exposed miniscaffoldins and the number of cohesin–dockerin complexes formed on the cell surface. These experiments, using pairs derived from *A. thermocellus* and *A. cellulolyticus*, suggested in agreement with earlier *in vitro*^32^ and *in silico*^33^ studies that thermophilic cohesin–dockerin pairs exhibit higher binding efficiency than their mesophilic counterparts. Yet two questions remained unresolved: which conditions maximize binding on the bacterial surface, and how stable are the resulting interactions under relevant conditions? Addressing these questions is important both for understanding the functional constraints of natural cellulosome modules and for adapting them as robust molecular connectors in engineered microbial platforms.

Here, we investigated the binding efficiency and interaction stability of two thermophilic cohesin–dockerin pairs from *A. thermocellus* and *Acetivibrio clariflavus* (formerly *Hungateiclostridium clariflavum*) on the surface of *P. putida* EM371 using a dockerin-tagged fluorescent reporter. We specifically sought to determine how interaction temperature affects complex formation and whether limited stability could be improved by targeted engineering of the dockerin and cohesin domains. We show that binding efficiency is strongly influenced by the temperature at which the interaction is formed and that interaction stability can be substantially enhanced by targeted amino acid substitutions guided by molecular dynamics simulations and free-energy calculations. These results define key parameters governing the performance of thermophilic cohesin–dockerin pairs on a bacterial surface and establish a computation-guided strategy for engineering more stable cellulosome-derived assembly modules for biotechnology, synthetic biology, and the sustainable valorization of polymeric lignocellulosic feedstocks.

## Experimental Section

### Bacterial strains, media, and growth conditions

Bacterial strains used in this study are summarized in **Table S1**. *Escherichia coli* CC118 was used for cloning. Reporter proteins for purification were produced in *Escherichia coli* BL21-Gold (DE3). Synthetic scaffoldins were produced and displayed on the surface of *Pseudomonas putida* EM371.^35^ All strains were grown in lysogeny broth (LB; 10 g L^−1^ tryptone, 5 g L^−1^ yeast extract, 5 g L^−1^ NaCl, pH 7.0; for solid medium, 16 g L^-1^ of bacteriological agar was added). When appropriate, antibiotics were added at the following final concentrations: ampicillin (Amp) 150 µg mL^−1^ and kanamycin (Km) 50 µg mL^−1^. For all experiments, overnight (16 h) precultures were prepared in 2.5 mL of LB medium and grown at 30°C (*P. putida*) or 37°C (*E. coli*) in a shaking incubator at 300 rpm (Heidolph Unimax 1010 and Heidolph Incubator 1000; Heidolph Instruments). The cultivation parameters of the main cultures are described in the following sections.

### Construction of plasmids and strains

All plasmids and plasmid constructs used in this study are summarized in **Table S2**. The sequences encoding HcCoh (GenBank^36^ accession number WP_052306610.1; 1^st^ Coh in the protein) and HcDoc (WP_014256341.1) were codon-optimized for expression in *P. putida* using GeneArt software (Thermo Fisher Scientific) and commercially synthesized (Eurofins Scientific).

Commercial reagents were used according to the manufacturers’ instructions unless stated otherwise. Plasmids were isolated from 5 mL LB overnight cultures using the EZNA Plasmid DNA Mini Kit (Omega BIO-TEK). Q5 High-Fidelity DNA Polymerase with its buffer, Q5 High GC Enhancer (both from New England Biolabs), and PCR Nucleotide Mix (10 mM each dNTP; Roche) were used to amplify genes for cloning. VeriFi PCR Mix Red (PCRbio) was used for difficult amplifications. Oligonucleotide primers (**Table S3**) for all reactions were purchased from Merck. Two-step overlap extension PCR (SOEing) was used to construct chimeric genes.^37^ The amplicons prepared by SOEing were subcloned into target plasmids using restriction cloning with restriction enzymes (for details, see **Table S2**), CutSmart buffer, and T4 DNA ligase from New England Biolabs. The resulting ligation mixtures were used to transform chemocompetent *E. coli* CC118 cells by heat shock (90 s at 42°C). The transformants were selected on LB plates with the corresponding antibiotics and verified by colony PCR (DreamTaq Green Master Mix (2x); Thermo Fisher Scientific), restriction analysis, and Sanger sequencing (SEQme s.r.o., Czech Republic). Verified clones were stored in LB with 20% (v/v) glycerol at -70°C.

DNA concentrations were measured throughout the cloning procedure using a NanoDrop 2000/2000c spectrophotometer (Thermo Fisher Scientific). DNA fragments and amplicons were visualized by gel electrophoresis (agarose concentration varied from 0.8% to 2% according to the expected length of DNA) using SYBR Safe DNA gel stain (Invitrogen).

Plasmids isolated from the cloning strain (*E. coli* CC118) were transferred to the production strains by heat shock-induced transformation or electroporation in the case of *E. coli* BL21-Gold(DE3) or *P. putida* EM371, respectively. Electroporation (2.5 kV, 4−5 ms pulse) was performed using the GenePulser Xcell electroporator equipped with 2 mm-gap cuvettes (Bio-Rad). Electrocompetent cells of *P. putida* were prepared as described by Aparicio and colleagues.^38^ All transformants of production strains were verified by colony PCR and restriction analysis before use in further work.

### Detection of scaffoldin expression in *P. putida* EM371

Precultures of *P. putida* EM371 strains carrying pSEVA238b*_ag43AT*, pSEVA238b*_ag43_hcCoh-ctCoh-his*, pSEVA238b*_ag43_hcCoh^m^*^1^*-ctCoh-his*, or pSEVA238b*_ag43_hcCoh^m^*^2^*-ctCoh-his* were prepared as described above. Main cultures of 20 mL LB with the appropriate antibiotics were inoculated to an initial OD_600_ of 0.05. The cultures were grown at 30°C and 300 rpm (Unimax 1010) for 2.5 h. I0 samples were collected for SDS-PAGE and western blotting (2 mL of culture was centrifuged at 4,000 rpm; 10 min; 4°C). Scaffoldin production was induced by 3-methylbenzoate (3MB; 0.5 mM final concentration), and the cultivation continued for 4.5 h. Samples I were collected (2 mL of culture was centrifuged at 4,000 rpm; 10 min; 4°C) at the end of cultivation. Sample preparation, SDS-PAGE, and western blotting were performed as described in **Supplementary methods**.

### Designer scaffoldin display assay: Testing the binding efficiency of the Coh-Doc pairs

Precultures of *P. putida* strains with plasmids encoding the synthetic scaffoldins were prepared as described in **Bacterial strains, media, and growth conditions**, with the addition of 10 mM CaCl_2_. Main cultures of 10 mL LB with the appropriate antibiotics and 10 mM CaCl_2_ were inoculated to an initial OD_600_ of 0.05 and grown at 30°C, 300 rpm (Unimax 1010). After 2.5 h, scaffoldin production was induced by 3MB (final concentration 0.5 mM). The cells were cultured for 4.5 h and then harvested by centrifugation (4,000 rpm; 10 min; 4°C). The resulting pellets were washed with 10 mL of TBS-Ca-T buffer and centrifuged again (4,000 rpm; 10 min; 4°C). Cells were resuspended in TBS-Ca-T buffer to OD_600_ = 10.0 and divided into 0.5 ml aliquots (∼2.275 × 10^9^ cells per sample).^17^ The cells were mixed with 25 µL of purified reporter protein (∼4 × 10^14^ molecules; the molecular weight was calculated using the ExPASy^39^ Compute pI/Mw tool; https://web.expasy.org; **Supplementary methods**), resulting in a ratio of ∼1.8 × 10^5^ reporter protein molecules per bacterial cell.

Cells displaying scaffoldins were incubated with the fluorescent dockerin-tagged reporter proteins by slow mixing in various conditions: i) at 4°C for 13 h; ii) at 30°C for 1 h; iii) at 50°C for 40 min; or iv) at 50°C for 1 h. The samples were then divided into two samples of 240 µL and washed twice with 1 mL of ice-cold TBS-Ca-T buffer (centrifugation: 5,000 g; 3 min; 4°C). After the second wash, the cells were resuspended in 1.25 mL TBS-Ca-T. Samples of 150 µL were transferred to a black 96-well microtiter plate (Thermo Fisher Scientific) and measured using the Infinite 200Pro microplate reader (Tecan). OD_600_ was measured for each sample. Cell-associated reporter proteins were quantified by top-read fluorescence measurements. The excitation and emission wavelengths were set at 560 and 600 nm, with the bandwidths of 9 and 20 nm, respectively. The optimal gain (186) was determined in the first experiment and used for all subsequent measurements. Fluorescence values were normalized to cell density (OD_600_).

### Designer scaffoldin display assay: Testing the interaction stability of the Coh-Doc pairs

After binding the dockerin-tagged mScarlet reporter proteins to their respective cohesins displayed on the cell surface of *P. putida* at i) 30°C for 1 h, or ii) 50°C for 40 min (see **Testing the binding efficiency of the Coh-Doc pairs**), the samples were washed with 1 mL of ice-cold TBS-Ca-T buffer (centrifugation: 5,000 g; 3 min; 4°C) and resuspended in 0.5 mL of ice-cold TBS-Ca-T. Samples were subsequently incubated at 30°C with slow mixing. Aliquots (50 µL) were collected after 0, 1, and 3 h. All samples were then washed twice with 200 µl of TBS-Ca-T buffer (centrifugation: 5,000 g; 3 min; 4°C). After the final centrifugation, the samples were resuspended in 250 µl of TBS-Ca-T. Samples of 150 µl were measured as described in **Testing the binding efficiency of the Coh-Doc pairs**.

### Structural model of HcCoh-HcDoc

The 3D structure of the wild-type HcCoh-HcDoc complex (including two Ca2+ ions) was predicted by AlphaFold3 webserver on the 10th of June 2024 with model seed 154702551.^40^ The sequences of HcCoh and HcDoc were obtained from GenBank^36^ (accession numbers WP_052306610.1 (1^st^ Coh) and WP_014256341.1, respectively). The model ranked 0 was selected for further steps. The structure was protonated at pH 7.2 (matching TBS-Ca-T buffer) using H++ webserver^41^ with the following settings: salinity 0.15 M (matching TBS-Ca-T buffer composition); internal and external dielectric constants 10 and 80, respectively. The structured inner hydration shell was predicted with the 3D-RISM program^42^ from the AmberTools22 suite^43^ using the Kovalenko–Hirata closure, following the standard Placevent workflow^44^ at 323 K and 0.15 M NaCl and removing water molecules within 2.0 Å of the solute. The solute, together with this inner shell, was placed in a truncated octahedron of OPC waters^45^ with a 12 Å solute-to-edge distance using tleap module of AmberTools22^43^. Na⁺ and Cl⁻ ions were added proportionally to the water count to reach 0.15 M and to enforce electroneutrality. To enable 4 fs timestep, the hydrogen mass repartitioning^46^ was performed by the parmed^47^ module of AmberTools22^43^.

### Molecular dynamics (MD) simulation

All calculations were performed with Amber22^48^ (pmemd.MPI for minimization, pmemd.cuda for dynamics). The protein complex was described with ff19SB force field,^49^ water molecules with the OPC model,^45^ monovalent ions (Na^+^, Cl^-^) were described using Joung-Cheatham^50^ parameters and divalent ions (Ca^2+^) using the Li/Merz^51^ parameter set. A 8.0 Å cutoff was used for non-bonded interactions and long-range electrostatics were treated with the Particle Mesh Ewald method^52^ in all calculations, whereas SHAKE method^53^ was applied to constrain all bonds involving hydrogen during all MD simulations. Energy minimization was carried out in five sequential 500-step rounds (100 steps steepest descent followed by 400 steps conjugate gradient) with a stepwise release of harmonic positional restraints. The first round restrained all heavy atoms of the complex, including the two Ca²⁺ ions at 500 kcal·mol⁻¹·Å⁻²; the next four rounds restrained the protein backbone heavy atoms and the two Ca²⁺ ions at 500, 125, 25, and 0.0001 kcal·mol⁻¹·Å⁻², respectively. Equilibration was performed in four stages, all using a Langevin thermostat^54^ with a collision frequency of 2.0 ps⁻¹. First, the system was heated to 200 K over 20 ps under NVT with all heavy atoms of the complex, including Ca²⁺ ions, restrained at 5 kcal·mol⁻¹·Å⁻². A 1 ns NVT simulation then increased the temperature to 323 K over the first 100 ps and then held it constant with the same restraints. This was followed by 1 ns NPT simulation at 323 K and 1 bar with the protein backbone and Ca²⁺ ions restrained at 5 kcal·mol⁻¹·Å⁻², and a final 1 ns of unrestrained NPT. Pressure during NPT equilibration was controlled with a Berendsen barostat^55^ with a 1 ps relaxation time. Finally, five independent 500 ns unrestrained NPT production simulations (2.5 μs aggregate) were run at 323 K and 1 bar using the Langevin thermostat^54^ and a Monte Carlo barostat.^56^ Coordinates were saved every 200 ps, resulting in 2 500 frames per replica.

### Trajectory analysis and postprocessing

The cpptraj^57^ module of AmberTools23^43^ was used for trajectory analysis. First, solvent molecules were stripped, and periodic boundary conditions were autoimaged. Convergence was assessed by monitoring backbone and Ca^2+^ ions root mean square deviation (RMSD) relative to the minimized structure, temperature, and total energy over time, together with per-residue root mean square fluctuation (RMSF) of protein backbone (terminal residues were excluded from analysis). Then for further analysis, five replicas were merged, keeping every 5th frame, resulting in a combined trajectory of 2500 frames, covering the aggregated 2.5 μs simulation time.

### Identification of interface residues and their saturation mutagenesis

The trajectory was converted to ensemble of 2,500 pdb files including only protein complex with Ca^2+^ ions using cpptraj module.^57^ These snapshots were processed with FoldX 5.1.^58,59^ First, the *RepairPDB* module was used to identify and fix residues with steric clashes. Then the repaired structures were analyzed with *AnalyseComplex* module to identify residues forming the interface between HcCoh and HcDoc. Residues present in more than 50% of the analyzed structures were selected as candidates for mutagenesis. Next, the *PositionScan* module with the default settings was used for the saturation mutagenesis, and the resulting change in energy was averaged across all snapshots. From the resulting mutant library, stabilizing substitutions with ΔΔG < -1 kcal·mol^-1^ were selected.

### Identification of key residues stabilizing HcCoh-HcDoc wild-type complex

The single-trajectory per-residue binding free energy decomposition was performed with MMPBSA.py^60,61^ module of AmberTools25^43^ to identify key residues contributing to either stabilization or destabilization of the complex (|ΔG| > 1 kcal·mol^-1^). The Generalized Born model igb = 8 with salt concentration of 0.15 M and default values for the remaining parameters were used for this calculation on the 2,500 frames, considering HcDoc + two Ca^2+^ ions as receptor and HcCoh as ligand; entropic contributions were not included. List of stabilizing residues (ΔG < -1 kcal·mol^-1^) were used to filter out mutant library.

### Construction and ranking of combinatorial mutant library

The promising selected mutations were combined to form library of 1-, 2- and 3-point mutants. The *BuildModel* module from FoldX 5.1^58,59^ was then employed to predict structure and stability change of these mutant complexes. Finally, the structures of mutant complexes were then processed through *RepairPDB* (removing potential clashes after introduced mutation) and *AnalyzeComplex* modules analogously to wild-type complexes to evaluate their interaction energies.

### Dissecting stabilizing mechanisms of introduced mutations

Four mutants selected for experimental evaluation, the mutated residues from either HcCoh or HcDoc were introduced to the already protonated structure of wild-type complex. These mutant complexes then underwent MD simulations following the same protocol used for wild-type complex. Similarly, the analogous analyses and trajectory postprocessing were performed. Finally, the single-trajectory pairwise binding free energy decomposition was performed MMPBSA.py^60,61^ module of AmberTools25^43^ to explore changes in residue-residue interactions between the mutated residues and the rest of the complex. These calculations were performed on 2,500 frames per mutant, employing MMPBSA nonpolar optimization method inp = 2 (default), internal dielectric constants of 5 and 10 (calculation conducted with both parameters), external dielectric constant of 80, ionic strength of 0.15 M, and default values for the remaining parameters. For double mutants in HcCoh, the individual contributions of each mutated position were summed to obtain the total predicted ΔΔG (e.g., S37/N39), whereas single mutants in HcDoc were evaluated based on the contribution of interaction with one position alone.

### Data and statistical analyses

The number of independent experiments or biological replicates is indicated in the figure and table legends. Reported results include mean values and their standard deviations, calculated in Microsoft Office Excel 2013. Where appropriate, statistical significance was assessed using two-tailed Student’s t-tests performed in Excel (Microsoft O365). Confidence intervals were calculated for selected parameters as indicated in the corresponding figure and table legends.

## Results and Discussion

### Cohesin and dockerins from thermophilic *Acetivibrio clariflavus* can be functionally produced in and displayed on the surface of mesophilic bacterial hosts

We first constructed a binary HcCoh–CtCoh scaffoldin (**Fig. 2A**) containing cohesins from *Acetivibrio clariflavus* (HcCoh) and *Acetivibrio thermocellus* (CtCoh) connected by a flexible linker derived from the *A. cellulolyticus* scaffoldin ScaC^62^ (**Supplementary sequence 1**). This linker was previously shown to support efficient assembly of a functional designer cellulosome.^63^ The dual-cohesin design enabled direct comparison of Hc and Ct cohesin–dockerin interactions within the same scaffoldin architecture while minimizing potential effects of differences in expression or surface display. The HcCoh–CtCoh scaffoldin was displayed on the surface of *Pseudomonas putida* EM371 using the Ag43 autotransporter, previously identified as an efficient surface-display system in this host.^17^ In our former study, the dual-cohesin scaffoldin was shown to anchor significantly more dockerin-tagged reporter proteins than single cohesins displayed on the surface of EM371 strain.^17^

**Figure 2.**
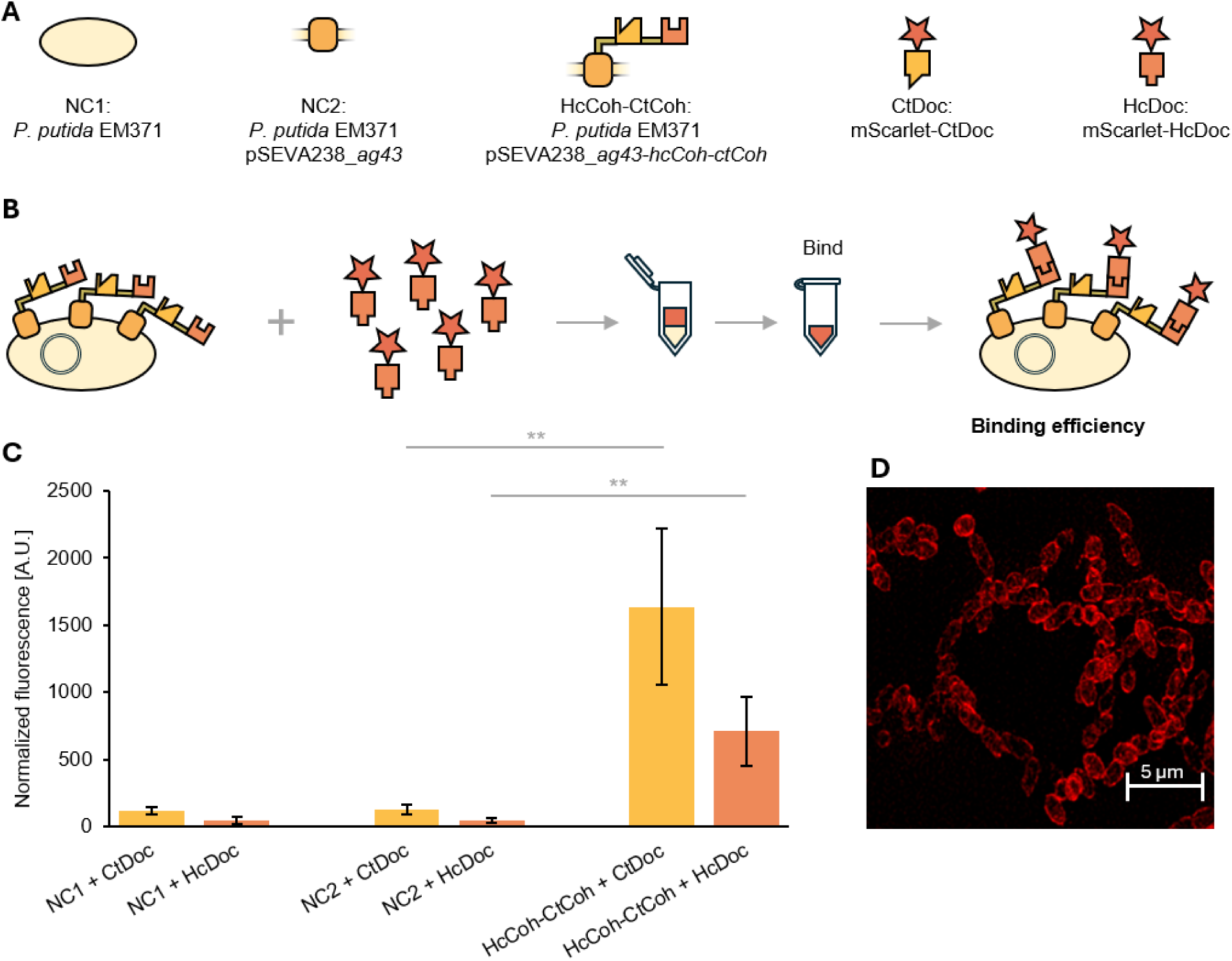
Overview of the strains, constructs, and the experimental approach used to test the properties of *Acetivibrio thermocellus* (Ct) and *Acetivibrio clariflavus* (Hc) cohesin-dockerin pairs when displayed on the surface of *Pseudomonas putida* EM371. (A) Pictograms and abbreviations for strains and reporter proteins used in this study. (B) The experimental workflow used to estimate binding efficiency of cohesin-dockerin pairs from normalized fluorescence measurements after initiating the interaction at 4°C overnight (C), as previously described.^17^ Yellow and orange colors denote the data measured for the Ct and Hc Coh-Doc pairs, respectively. Data are shown as mean ± SD from four biological replicates collected in two independent experiments. Asterisk denotes significance in the difference between two means at p < 0.01 (**). (D) Microphotography of *P. putida* EM371 cells with the mScarlet-I fluorescent protein attached to their surface through the interaction of protein-born dockerin and cell-displayed cohesin.

Previously, fluorescence of *P. putida* EM371 cells bearing surface-attached dockerin-tagged fluorophores was established as a robust proxy for monitoring the number of formed cohesin–dockerin pairs, with the results further validated by fluorescence microscopy, enzyme activity assays, and whole-cell growth experiments.^17^ Therefore, to monitor cohesin–dockerin interactions in the current work, we generated mScarlet-I reporters fused to either HcDoc or CtDoc (**Fig. 2A**; **Supplementary sequences 2** and **3**).

Codon-optimized HcCoh and HcDoc genes were assembled into the scaffoldin-display plasmid pSEVA238_*ag43AT-hcCoh-ctCoh* and the reporter plasmid pET21b+_*mScarlet-I-hcDoc-his*, respectively. These plasmids and pET21b+_*mScarlet-I-ctDoc-his* were introduced into *P. putida* EM371 (pSEVA construct) or *E. coli* BL21-Gold (DE3) (pET constructs) . Dockerin-tagged reporter proteins were successfully produced in *E. coli* and purified by affinity chromatography (**Fig. S1**). Expression of the HcCoh–CtCoh scaffoldin in *P. putida* EM371 was confirmed by SDS-PAGE and Western blotting (**Fig. S2**).

Following verification of scaffoldin and reporter production, we evaluated cohesin–dockerin interactions using a fluorescence-based binding assay and protocol established in our previous study (**Fig. 2B**).^17^ Scaffoldin-displaying cells were incubated with purified dockerin-tagged reporters at 4°C overnight, then washed to remove unbound protein, and analyzed by measuring their fluorescence. Binding efficiency (**Fig. 2C**) was inferred from normalized fluorescence (fluorescence divided by the optical density of respective cells) measured after the initial reporter attachment.

*P. putida* EM371 cells with Ct or Hc cohesin–dockerin pairs formed on their surface produced significantly stronger fluorescence signals than their respective controls (for NC2, p = 1.5 × 10^-4^ and 1.6 × 10^-4^ in the case of the Ct and Hc pair, respectively; **Fig. 2C**), confirming that the reporter fusions retained functionality and that HcCoh can be functionally displayed on the surface of *P. putida*. This was further supported by subsequent observation of EM371 cells with surface-attached mScarlet reporter molecules in a confocal microscope (**Fig. 2D**). Together, these results established a robust platform for comparative analysis of thermophilic cohesin–dockerin interactions on the surface of an industrially relevant Gram-negative mesophilic bacterial host.

### The binding efficiency of thermophilic Coh-Doc pairs increases if the interaction is initiated at higher temperatures

*Acetivibrio thermocellus* and *A. clariflavus* are among the few thermophilic bacteria known to produce cellulosomes, with optimal growth temperatures of approximately 60–64 °C and 55–60 °C, respectively.^64^ Their cohesin–dockerin interactions are attractive targets for protein engineering because thermophilic proteins typically exhibit high stability and are therefore well suited for industrial biocatalytic applications.^65^ However, most described cellulosome-producing organisms and almost all established microbial hosts for designer cellulosome testing are mesophiles.^66^ Mesophiles have been preferred due to the availability of advanced engineering tools and better genetic tractability. As a consequence, engineered cellulosome systems often combine thermophilic and mesophilic modules, and selecting an appropriate temperature for initiating cohesin–dockerin interactions is not straightforward. Previous studies have initiated these interactions over a wide temperature range, from low temperatures^17,67–71^ (≤10 °C) through moderate temperatures^24,27,28,32,63,67,69,72–77^ (20–37 °C) to high temperatures^77–79^ reaching those encountered by thermophilic organisms (≥50 °C). In some *in vitro* studies, cohesin–dockerin complexes were first assembled at moderate temperatures (typically 37 °C), whereas the stability or enzymatic activity of the resulting protein complexes, including attached cellulolytic enzymes, was subsequently evaluated at elevated temperatures (50–80 °C).^32,73–75^ Here, we investigated how the temperature used to initiate complex formation influences the binding efficiency of thermophilic Ct and Hc Coh–Doc pairs displayed on the surface of *P. putida* EM371.

To evaluate the effect of interaction temperature on thermophilic Coh–Doc complex formation, we selected three temperatures representing distinct biologically relevant conditions. The 4 °C condition reflects temperatures commonly used to initiate Coh–Doc interactions^17,67–71^ and minimize protein degradation during binding assays. The 30 °C condition corresponds to the optimal growth temperature of the mesophilic host *P. putida* EM371. Finally, 50 °C was chosen as a compromise between maintaining the mesophilic host integrity and approaching the optimal growth temperatures of the thermophilic bacteria *A. thermocellus* and *A. clariflavus* from which the tested Coh–Doc pairs originate. This temperature should be compatible with the use of *P. putida* as a surface-display platform, as Tozakidis *et al*. previously showed that approximately 70% of KT2440 cells remained structurally intact after incubation at 55 °C for 24 h.^80^ Although the cells exhibited signs of arrested metabolism, including reduced glucose uptake, they retained their ability to serve as passive carriers of surface-displayed proteins.

The herein measured binding efficiency depended strongly on the temperature at which the cohesin–dockerin complex was formed (**Fig. 3A–C**). Across all tested conditions, the Ct pair consistently outperformed the Hc pair (p = 0.002, 0.040, and 0.002 for 4 °C, 30 °C, and 50 °C, respectively). For the *P. putida* attaching the reporter through the Ct pair, normalized fluorescence remained similar after incubation at 4 °C (1,637 ± 580 arbitrary units, or A.U.) and 30 °C (1,821 ± 603 A.U.; p = 0.543), but increased markedly after incubation at 50 °C, reaching 5,066 ± 746 A.U. (p < 10^−6^ versus both lower temperatures). In contrast, the Hc pair-containing samples showed a progressive increase in binding efficiency with temperature, with the normalized fluorescence rising from 710 ± 259 A.U. at 4 °C to 1,227 ± 416 A.U. at 30 °C (p = 0.012) and 3,264 ± 139 A.U. at 50 °C. Together, these results demonstrate that elevated temperatures promote formation of thermophilic Coh–Doc complexes on the bacterial surface, although the magnitude of the effect differs between the Ct and Hc pairs. The improvement observed after high-temperature initiation may reflect more favorable complex assembly under conditions closer to the physiological environment of the source organism. This effect may be explained by the increased conformational sampling of the interacting proteins at elevated temperatures, allowing them to overcome local energetic barriers and adopt binding-competent conformations that favor formation of the Coh–Doc complex.^81,82^

**Figure 3.**
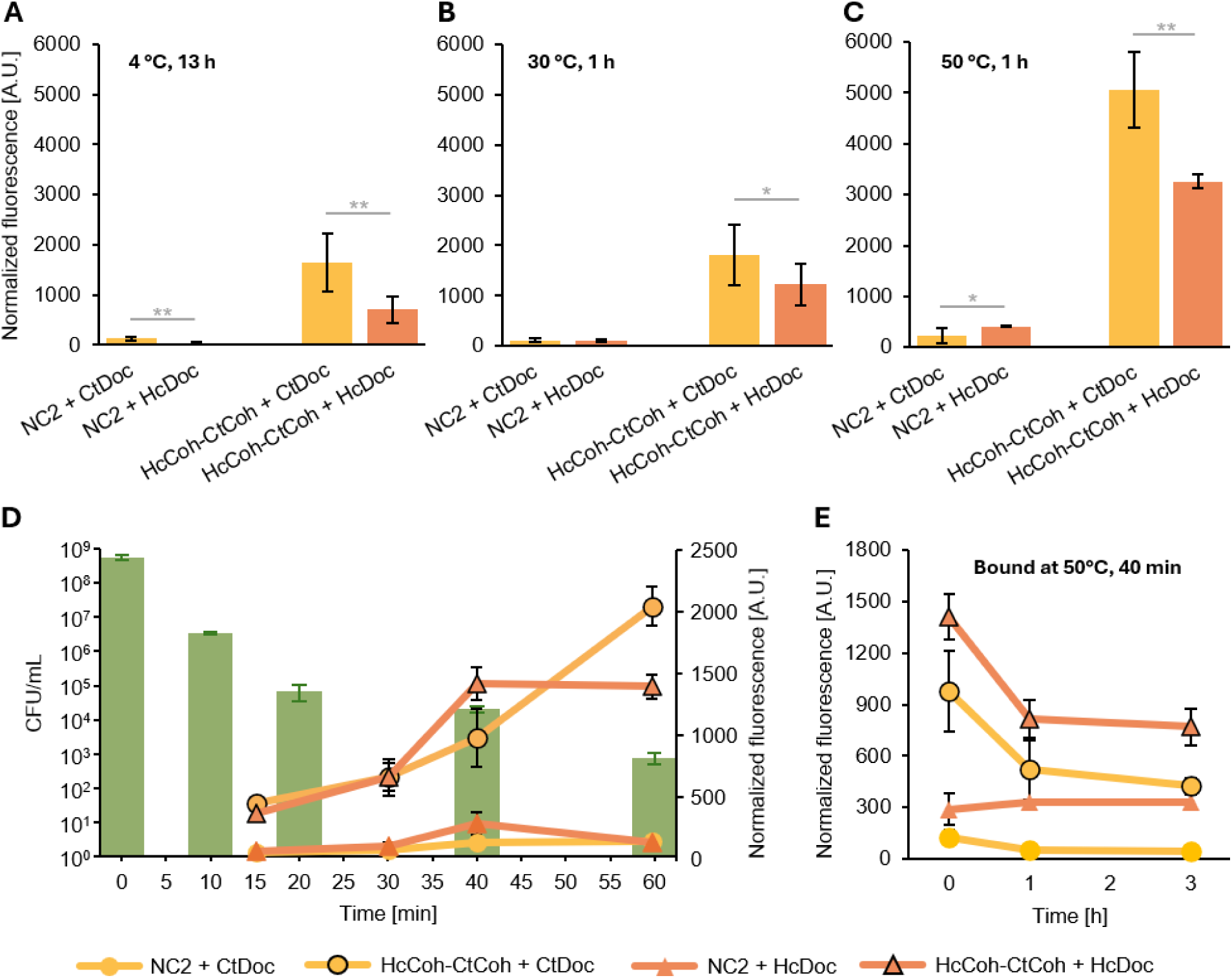
Testing the binding efficiency and interaction stability of thermophilic Coh-Doc pairs across different conditions. (A-C) Binding efficiency (represented by normalized fluorescence of mScarlet attached to the cell surface of *P. putida* EM371 via the Ct or Hc Coh-Doc interaction) after initiating the interaction at 4 °C for 13 h (A), at 30 °C for 1 h (B), and at 50 °C for 1 h (C). (D, main axis) Survival of *P. putida* EM371 at 50 °C determined as the number of colony-forming units (CFU) per 1 mL of cell suspension. (D, secondary axis) Screening of the effect of the length of incubation at 50 °C on the binding efficiency of the Hc and Ct Coh-Doc pairs. (E) Interaction stability of the Ct and Hc Coh-Doc pairs at 30 °C after initiating the interaction at 50 °C for 40 min. Data are shown as mean ± SD from: at least three biological replicates (two technical replicates each; A, B, C); one biological replicate with two or three technical replicates (D); or four biological replicates measured in two independent experiments (E). Asterisk denotes significance in the difference between two means at p < 0.05 (*) or at p < 0.01 (**).

Although incubation at 50 °C for 1 h (**Fig. 3C**) produced the strongest normalized fluorescence signals, it also resulted in cell aggregation that complicated sample handling (**Fig. S3**). To identify a more practical condition, we monitored cell viability and binding performance during incubation at 50 °C (**Fig. 3D**). CFU (colony-forming unit) counts decreased from 5.60 × 10^8^ CFU mL^−1^ before incubation to 8 × 10^2^ CFU mL^−1^ after 60 min, indicating substantial loss of viability. At the same time, binding signals continued to increase with incubation time. We eventually selected 50 °C applied for 40 min as a compromise between binding performance and cell integrity. Under these conditions, cells remained structurally intact (**Fig. S4**) and did not exhibit the extensive aggregation observed after 60 min incubation. This observation contrasts indirectly with the results of Tozakidis *et al.* who recovered 70% of *P. putida* cells after 24 h at 55 °C.^80^ The difference can be attributed to the use of different strains of *P. putida*, with the genome-derived EM371 strain used here being better suited for recombinant protein display but less robust and viable at the elevated temperature than the wild-type strain KT2440 used by Tozakidis and colleagues.^17,35^

### The suboptimal interaction stability of thermophilic Coh-Doc pairs on the *P. putida* surface

Despite its importance for the performance of cell-surface assemblies, interaction stability is rarely evaluated in studies of cohesin–dockerin systems.^16^ In our previous study, we did not assess the stability of cohesin–dockerin interactions formed on the surface of *P. putida*.^17^ Here, we focused on this key property of the system. Binding efficiency was estimated from the normalized fluorescence measured immediately after the initial attachment of the reporter proteins at 50 °C for 40 min (**Fig. 3E**, 0 h). Interaction stability was then evaluated from the normalized fluorescence retained after subsequent incubation of reporter-bound cells at 30 °C (host’s optimal temperature securing its cellular integrity), followed by washing after 1 and 3 h (**Fig. 3E**). Although retained normalized fluorescence may also be influenced by factors unrelated to Coh–Doc dissociation, the use of identical scaffoldin architectures, reporter constructs, and incubation conditions across all tested variants allows relative differences to be interpreted primarily as differences in Coh–Doc interaction stability

The two thermophilic Coh–Doc pairs responded similarly in the interaction stability assay (**Fig. 3E**). For cells displaying either Coh–Doc pair, most signal loss occurred during the first one hour of incubation, after which fluorescence remained relatively stable. The cells with the Hc and Ct Coh-Doc pairs retained 58 ± 11 % and 54 ± 20 % of their fluorescence after 1 h of incubation, respectively.

The interactions observed on the surface of *P. putida* remained less stable than those typically reported for purified Coh–Doc complexes.^32,73–75^ Diverse factors could contribute to this effect. Our first consideration was the intrinsic stability of the Coh–Doc pairs used in this study. Although Coh–Doc interactions are often described as being among the strongest known non-covalent protein–protein interactions, high binding affinity does not necessarily make assembled complexes irreversible. Borne and colleagues demonstrated that dockerin-bearing enzymes could displace catalytic subunits from assembled miniscaffoldin complexes and native cellulosomes and that the stability of these complexes depended on the dockerin sequence.^83^ These findings indicate that high binding affinity alone does not guarantee long-term complex stability and that the kinetic robustness of Coh–Doc interactions varies among systems. Such variability may enable the remodelling of cellulosome composition in response to changes in substrate availability or environmental conditions.^66,84^

Host- and condition-specific factors may also contribute. The unique surface properties of the genome-streamlined strain *P. putida* EM371 may influence the behavior of the displayed complexes through electrostatic interactions with the scaffoldin itself.^35^ It is also conceivable that bacterial cells sequester Ca²⁺ ions required for stable Coh–Doc interactions, for example through the local microenvironmental competition due to surface charge effects or calcium binding to the extracellular matrix and large adhesins such as LapF.^85^ However, this possibility is unlikely, considering the relatively high CaCl₂ concentration (10 mM) in the incubation buffer and the absence of numerous native outer membrane structures, including lipopolysaccharides or LapF, in the “naked” *P. putida* EM371 strain.^35^Another possible explanation is the relatively low temperature used during the stability assay. Although the interactions were initiated at 50 °C, subsequent incubation was performed at 30 °C, well below the optimal growth temperatures of both *A. thermocellus* and *A. clariflavus*.^64^ It is therefore conceivable that interaction stability would be further enhanced if the complexes were maintained at elevated temperatures throughout the experiment. This hypothesis could not be tested in the present system because prolonged incubation at 50 °C compromised the integrity of *P. putida* cells. However, evidence from other surface-display platforms suggests that temperature alone is unlikely to fully explain the observed instability. Stern and colleagues reported that designer cellulosomes comprising both mesophilic and thermophilic Coh–Doc pairs retained 94 ± 6% to 100 ± 0% activity after 48 h at 37 °C when displayed on the surface of *Lactobacillus plantarum*.^16^ Similarly, Wang and colleagues observed approximately 85%, 70%, and 30% activity retention after 60 min at 60, 65, and 70 °C, respectively, for β-galactosidase attached to *Bacillus subtilis* spores through mesophilic Coh–Doc interactions.^86^ Overall, the observed instability most likely results from a combination of factors, including the specific display architecture, host surface properties, the non-native assay conditions, and the intrinsic stability of the studied Coh-Doc pairs. Because the contribution of each factor cannot be separated within the present experimental framework, we focused on the parameter most amenable to rational improvement, namely the molecular stability of the interaction, through protein engineering.

Protein engineering has recently been used to enhance the stability of cohesin–dockerin complexes by covalently locking the interacting partners, either through engineered intermolecular disulfide bonds or by incorporating the SpyTag–SpyCatcher system.^30,87^ We sought to enhance the stability of complexes assembled on the surface of *P. putida* through computationally guided protein engineering while preserving the native non-covalent nature of the cohesin–dockerin interaction. The Hc pair was selected as the engineering target because it exhibited high binding efficiency under the selected interaction conditions (**Fig. 3D,E**) while still providing substantial room for improvement in interaction stability. Furthermore, the Hc pair is understudied compared to the widely used Ct system, and it originates from *A. clariflavus* with a temperature optimum (55–60 °C) closer to the 50 °C used here for binding.

### Computation-guided mutagenesis identifies substitutions with the potential to improve the interaction stability of the Hc Coh-Doc pair

The rational design and computational screening workflow employed in this study is outlined in **Fig. 4**. Briefly, ensemble-based free energy calculations were used to identify and prioritize mutations in both HcCoh and HcDoc with the potential to enhance complex stability.

**Figure 4.**
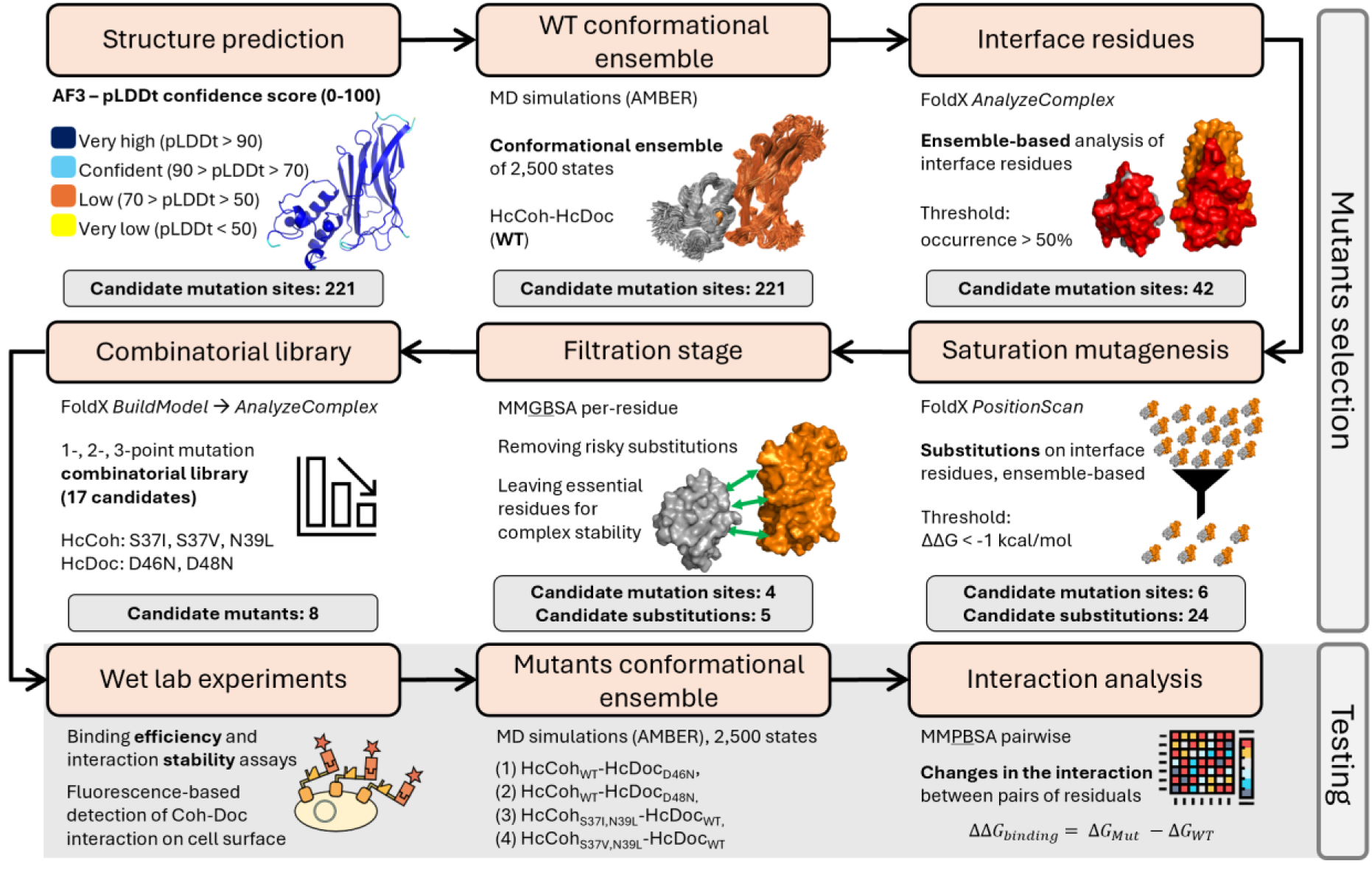
Computational workflow for rational design of stabilizing HcCoh-HcDoc interface mutants. The workflow consists of two stages: mutant selection (top) and testing (bottom).

To date, the 3D structure of the HcCoh-HcDoc complex has not been experimentally determined. Hence, we used AlphaFold3^40^ (AF3) to predict the structure of the HcCoh-HcDoc complex. The AF3 generated structure exhibits overall high confidence, with a mean of pLDDT (predicted Local Distance Difference Test) score per residue of 93.2/100, indicating high local structural accuracy. The model’s overall structural accuracy evaluated by pTM (predicted Template Modelling) was 0.9 (on a scale from 0-1, with values >0.5 indicating confident global fold prediction), while mean PAE (predicted Aligned Error) was less than 2 Å assessed accuracy of relative arrangements of HcCoh and HcDoc (in the complex values <5 Å indicating high confidence in relative positioning).

Five independent MD simulations of wild-type HcCoh-HcDoc complex were carried out to accumulate 2.5 μs overall sampling. The stability and convergence of these simulations were confirmed by RMSD and RMSF analyses (**Fig. S5**). Since the aim was to improve the stability of the complex, the next step was to determine which residues form the interface between HcCoh and HcDoc. We analyzed the interface on the 2,500 conformations assembled from the five replicated simulations, yielding 42 residues that form the interface in more than 50% of conformations (**Table S4**). Selected interface residues were then subjected to saturation mutagenesis using FoldX.^58^ At this step, each of the 42 interface residues was mutated to all 20 amino acids, and the most favorable substitutions (ΔΔG < -1 kcal/mol) were selected, leaving 24 candidate substitutions across 6 positions (**Table S5**).

Next, the MMGBSA (Molecular Mechanics / Generalized Born Surface Area) was used to calculate the free energy change between a bound and free state of a HcCoh and HcDoc across the ensemble of conformations. By decomposing the contribution of each residue to the binding free energy, the most crucial residues for the stability of the HcCoh-HcDoc complex were identified (**Table S6**). The 27 residues stabilizing the complex were evenly distributed to both binding partners, whereas the three residues exhibiting considerable unfavorable contribution to the complex stability were D61, D55, and D46 from HcDoc. To avoid potential disruption of the complex formation, all 27 stabilizing residues were excluded from the list of substitution candidates.

The list of 24 mutation variants was narrowed down by removing risky substitutions, which were identified based on the following criteria (**Table S5**): (i) disrupting stability hotspots (ΔΔG < -1 kcal/mol) identified earlier as mutation would disrupt already-stabilizing contacts; (ii) bringing a positive charge to the proximity of the coordinated Ca^2+^ in the HcDoc; (iii) disrupting the Ca^2+^-coordinating loop in HcDoc by substitutions other than asparagine at positions D46 and D48;^88^ (iv) introducing methionine at the interface as vulnerable to oxidation residue could lead to protein destabilization.^89^ The choice of asparagine as the only permitted substitution is supported by its conservation at calcium-coordinating positions of the *C. thermocellum* dockerin F-hand motif.^25^ After removing the risky substitutions, 5 variants remained: HcDoc – D46N, D48N; HcCoh – S37V, S37I, N39L. A combinatorial library of 1-, 2-, and 3-point mutants was prepared, yielding 17 combinations (**Table S7**), with the constraint that D46 and D48 were not mutated simultaneously. This limitation is supported by systematic mutagenesis showing that substituting one or even two positions within a single Ca^2+^ coordinating loop does not disrupt cohesin binding, but impairs calcium coordination, which is essential for complex formation.^25^ To increase the likelihood of successfully enhancing stability substitution, we decide to mutate only one position at once. Then these mutations were introduced into 2,500 conformations of wild-type HcCoh-HcDoc complex, and their impact on the stability of both target proteins as well as their mutual interactions (**Tables S7-S8**) was evaluated. Mutations had stabilizing (D46N, D48N, N39L) or neutral (S37V, S37I) effects on individual proteins, with the combined mutants exhibiting a cumulative decrease in energy, suggesting that the effects of individual substitutions are approximately additive (**Tables S7-S8**). Analysis of the energy contributions suggests that HcDoc mutations mainly improved electrostatic interactions; on the other hand, HcCoh mutations resulted in enhanced nonpolar solvation and van der Waals interactions, however at the expense of often introducing also considerable structural clashes (**Tables S7-S8**).

Since all evaluated combinations of mutations did not exhibit any considerable antagonistic behavior, the following 4 mutation variants were ultimately selected for further computational and laboratory experiments: HcDoc^m1^ bearing substitution D46N, HcDoc^m2^ with substitution D48N, HcCoh^m1^ with substitutions S37V and N39L, and HcCoh^m2^ bearing substitutions S37I and N39L.

### Mutant variants of HcDoc and HcCoh retain their ability to form functional complexes

To experimentally validate the computation-guided designs, we constructed two HcDoc variants, HcDoc^m1^ and HcDoc^m2^. We also constructed two HcCoh variants displayed within the HcCoh–CtCoh scaffoldin, HcCoh^m1^–CtCoh and HcCoh^m2^–CtCoh. The dockerin variants were produced as mScarlet-I fusions in *E. coli* and purified by affinity chromatography, whereas the cohesin variants were displayed on the surface of *P. putida* EM371 using the Ag43-based system. SDS-PAGE and Western blot analyses confirmed successful production of all reporter and scaffoldin variants (**Fig. S6**, **Fig. S7**), indicating that the introduced substitutions did not compromise protein production, purification, or surface display in mesophilic bacterial hosts.

Relative scaffoldin abundance in cell lysates was comparable among the wild-type HcCoh–CtCoh, HcCoh^m1^–CtCoh, and HcCoh^m2^–CtCoh variants, which accounted for 2.33 ± 0.15%, 2.87 ± 0.31%, and 2.40 ± 0.17% of total cellular protein, respectively. Although HcCoh^m1^–CtCoh showed a slightly higher apparent abundance than the wild-type scaffoldin, this difference was not statistically significant (p = 0.075; **Fig. S7C**). Notably, the abundance values measured for all three scaffoldins were comparable to those reported previously for related Ag43-displayed scaffoldins in *P. putida* EM371,^17^ indicating that the introduced mutations did not substantially affect scaffoldin production in the host cells.

We next tested whether the mutant Hc Coh–Doc variants retained their ability to form functional complexes under the interaction conditions used for the wild-type pairs (**Fig. 5A, Fig. S8A–D; Supplementary results**). After interaction initiation at 50 °C for 40 min, *P. putida* cells with most mutant combinations reached normalized fluorescence levels comparable to the cells with the wild-type pair. The only exception was cells with HcCoh–CtCoh + HcDoc^m2^, which showed significantly lower fluorescence than the cells with the wild-type pair (p = 0.045; **Fig. 5A**). Because normalized fluorescence serves as a proxy for reporter attachment to the cell surface, these results indicate that mutant variants retained the ability to form functional Coh–Doc complexes under the tested conditions.

**Figure 5.**
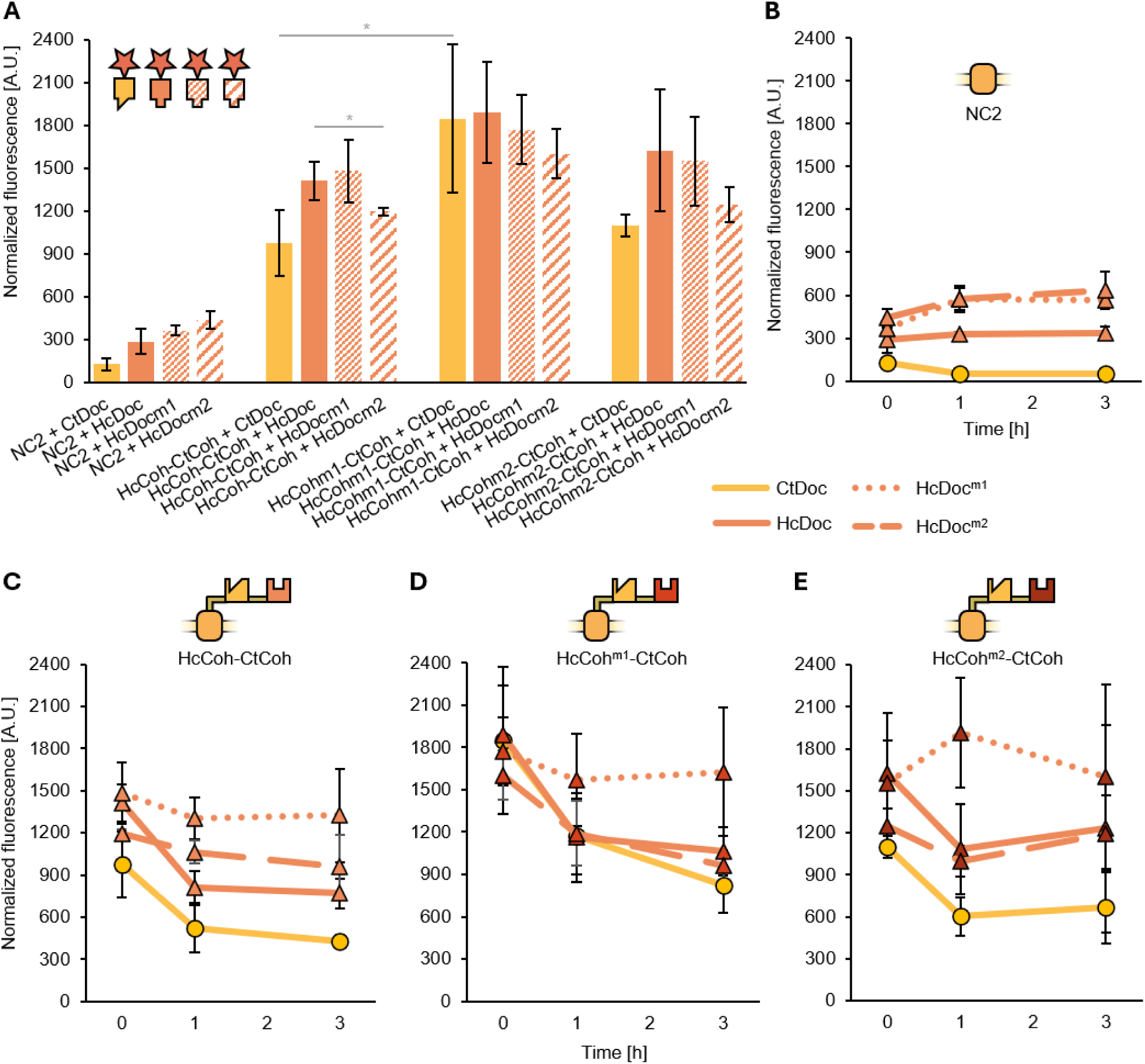
Amino acid substitutions in HcDoc retain the binding efficiency and improve the interaction stability of the Hc Coh-Doc pair. (A) Binding efficiency (represented by normalized fluorescence of mScarlet attached to the cell surface of *P. putida* EM371 via the Ct or Hc Coh-Doc interaction) after initiating the interaction at 50 °C for 40 min. (B-E) Interaction stability of Coh–Doc pairs represented by retention of normalized fluorescence of mScarlet attached to the cell surface of *P. putida* EM371 via the Ct or Hc Coh-Doc interaction (formed at 50 °C for 40 min) after 0, 1, and 3 hours of incubation at 30 °C. CtDoc, HcDoc, HcDoc^m1^, and HcDoc^m2^ were attached to the cell surface through non-specific interactions (negative control, B), or via the interaction with their respective cohesins in (C) wild-type HcCoh-CtCoh, (D) HcCoh^m1^-CtCoh, or (E) HcCoh^m2^-CtCoh. Data are shown as mean ± SD from four biological replicates measured in two independent experiments. Asterisk denotes significance in the difference between two means at p < 0.05 (*) or at p < 0.01 (**).

### Substitutions in HcDoc improve the interaction stability of the Hc Coh-Doc pair

We next investigated whether the computationally designed mutations improved the stability of Hc Coh–Doc interactions on *P. putida* EM371 cells under the optimized conditions established above. Coh–Doc complexes were formed at 50 °C for 40 min and subsequently incubated at 30 °C for 3 h (**Fig. 5B-E**, **Fig. 6**). Interaction stability was inferred from the percentage of normalized fluorescence retained relative to the signal measured with cells immediately after complex formation.

**Figure 6.**
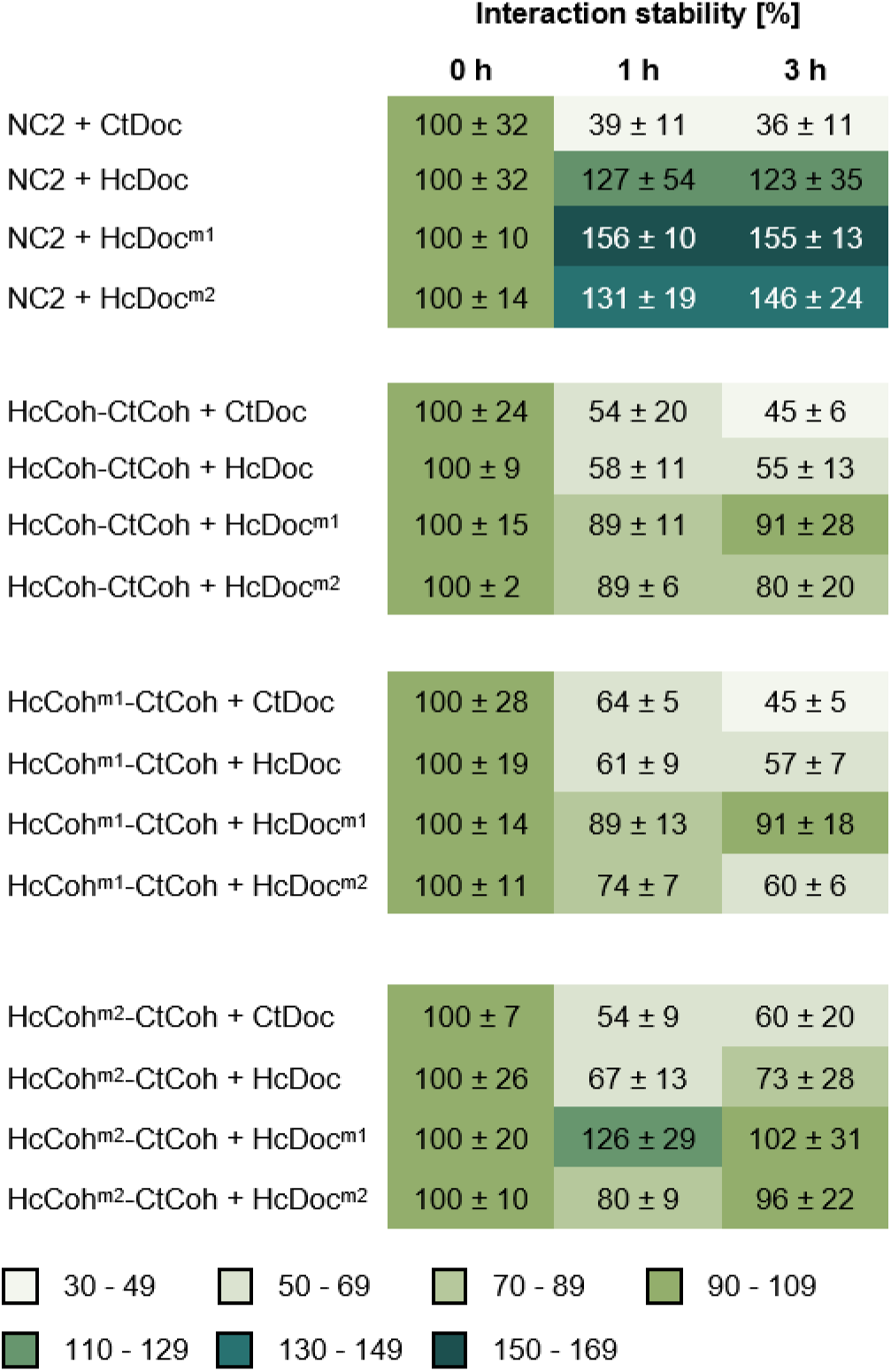
Interaction stability of Coh-Doc pairs represented by retained normalized fluorescence of mScarlet attached to the cell surface through Coh-Doc interactions. Dockerin-tagged reporter proteins were incubated with *P. putida* EM371 cells displaying synthetic scaffoldins at 50 °C for 40 min. After washing away the unbound proteins, the normalized fluorescence was measured (hour 0). The samples were then incubated at 30 °C. After 1 and 3 hours of incubation, samples of cells were withdrawn, washed, their normalized fluorescence was measured, and the retained normalized fluorescence was calculated as a percentage of the value measured at hour 0. Data are shown as mean ± SD from four biological replicates measured in two independent experiments.

Negative-control samples exhibited measurable nonspecific attachment of dockerin-tagged reporters to the cell surface (**Fig. 5B**, **Fig.6**). This trend was more pronounced for the mutant dockerins than for the wild-type reporter, suggesting that the introduced substitutions modestly increased nonspecific surface adherence. Although the underlying mechanism was not investigated, this behavior may reflect subtle changes in protein surface properties, including charge distribution or exposure of interaction-prone regions, resulting from the engineered amino acid substitutions.^90,91^

The *P. putida* cells with the wild-type Hc Coh–Doc pair retained 58 ± 11% and 55 ± 13% of their initial normalized fluorescence after 1 and 3 h of incubation at 30 °C, respectively (**Fig. 5C**, **Fig. 6**). Introducing mutations into the cohesin alone did not significantly improve interaction stability, as complexes containing HcCoh^m1^ (**Fig. 5D**, **Fig. 6**) or HcCoh^m2^ (**Fig. 5E**, **Fig. 6**) together with wild-type HcDoc exhibited retention values comparable to the wild type after 1 h (61 ± 9% and 67 ± 13%, respectively; p = 0.693 and 0.336).

In contrast, both dockerin variants improved the interaction stability. When HcDoc^m1^ or HcDoc^m2^ interacted with wild-type HcCoh (**Fig. 5C**, **Fig. 6**), 89 ± 11% and 89 ± 6% of the initial signal remained after 1 h, significantly exceeding the value observed for the wild-type pair (p = 7.29 × 10^−3^ and 4.64 × 10^−3^, respectively). The same trend remained apparent also after 3 h of incubation.

Several combinations of mutant cohesins and dockerins exhibited even greater improvements in interaction stability. In particular, *P. putida* cells attaching the fluorescent reporter through the HcCoh^m1^–CtCoh + HcDoc^m1^ and HcCoh^m2^–CtCoh + HcDoc^m2^ interactions retained 89 ± 13% and 80 ± 9% of the initial normalized fluorescence after 1 h, respectively, compared with 58 ± 11% for the wild-type pair (p = 0.012 and 0.026). These combinations of cohesins and dockerins remained significantly more stable than the wild-type pair also after 3 h of incubation (91 ± 18% and 96 ± 22% versus 55 ± 13%; p = 0.021 and 0.016, respectively). The *P. putida* cells with the HcCoh^m2^–CtCoh + HcDoc^m1^ combination exhibited the highest mean retained normalized fluorescence among all tested variants (126 ± 29% and 102 ± 31% after 1 and 3 h, respectively). Owing to substantial variability between biological replicates, these values could not be directly compared statistically with the wild-type pair. Nevertheless, the consistently elevated retained normalized fluorescence suggests that this variant also improves Coh–Doc interaction stability.

The fact that both dockerin variants improved stability in combination with multiple cohesin backgrounds indicates that the primary stabilizing effect originated from the amino acid substitutions in dockerins. Nevertheless, the superior performance of selected mutant Coh–Doc combinations suggests that cohesin substitutions can further enhance stability when combined with appropriately engineered dockerins, highlighting the importance of optimizing both interaction partners. These results collectively underscore the positive effect of the introduced mutations on Coh–Doc interaction stability. They further suggest that structure–function effects make a major contribution to the observed interaction instability, although adverse factors discussed above and others such as proteolytic degradation of surface-exposed complexes, structural constraints, or fluorophore quenching cannot be fully excluded.

Among all tested variants, the HcCoh^m1^–CtCoh + HcDoc^m1^ combination provided the most balanced improvement of the system. In addition to enhanced interaction stability, this variant produced higher absolute fluorescence signals on the surface of *P. putida* than the wild-type pair after both 1 and 3 h of incubation (1,573 ± 328 and 1,626 ± 457 A.U. versus 1,301 ± 149 and 1,325 ± 330 A.U., respectively; p = 0.014 and 0.030). This improvement likely reflects the combined effects of enhanced interaction stability and improved overall performance of the surface-displayed scaffoldin HcCoh^m1^-CtCoh (**Fig. 5A, Fig. S8E**, and **Supplementary results**). Consequently, the engineered system not only retained reporter proteins for longer periods but also maintained a greater number of surface-associated molecules over time.

To explore the impact of mutations in greater depth, MD simulations were conducted for single protein mutants: HcDoc-D46N (HcDoc^m1^), HcDoc-D48N (HcDoc^m2^), HcCoh-S37V/N39L (HcCoh^m1^), and HcCoh-S37I/N39L (HcCoh^m2^) to prepare an ensemble of 2,500 conformations, following the same protocol as the WT simulations. All simulations exhibited stable behavior, with RMSD and RMSF values comparable to those of the WT complex (**Figs. S9-S12**). The conformational ensemble was subjected to Molecular Mechanics / Poisson Boltzmann Surface Area (MMPBSA) calculations of binding free energy in pairwise decomposition mode. The differential reliability of FoldX for hydrophobic versus polar mutations documented by Slutzki et al. through direct comparison of predicted and experimental ΔΔG values at the cohesin–dockerin interface,^88^ and further quantified in a large-scale deep mutational scanning benchmark where FoldX achieved reasonable overall discrimination but showed its greatest disagreements with experiment at charged and polar interface positions informed our decision to use MMPBSA as a complementary and more rigorous evaluation for the electrostatic redesign on the HcDoc side.^92^ This approach allows changes in pairwise residue interactions to be observed and compared with the WT. The internal dielectric constant of 5 and 10 were tested, as higher values have been recommended in the literature for MMPBSA calculations.^93^ In pairwise decomposition calculations, the solvation free energies are not strictly pairwise decomposable since the dielectric boundary is a collective parameter of the whole system, not exactly additive.^60^ Therefore, the analysis is based on relative comparison between mutants and WT with focus on highest changes in interaction. Regardless of the internal dielectric constant, the overall calculated ΔG_binding_ (**Table S9**), was within 1.5 kcal/mol or less of WT for both HcCoh mutants (S37I+N39L, S37V+N39L), implying complex stability equivalent to that of WT. Conversely, both HcDoc (D46N, D48N) mutants produced complexes with markedly elevated stability, with ΔG_binding_ improvements of 6-17 kcal/mol depending on the dielectric constant used (**Table S9**).

Since these calculations were in line with the experimental observations, we have further used the energy decomposition to identify the key changes in pairwise interactions caused by the mutations, thereby hinting at their mechanism of action. Both HcDoc mutants, D46N and D48N, altered primarily the electrostatic interactions with other charged residues (**Fig. 7** and **Tables S10-S11**). The WT HcCoh-HcDoc complex features numerous charged residues on its surface (**Fig. 8A**): HcCoh carries 9 positively and 18 negatively charged residues, and two charged termini, while HcDoc carries 7 positively and 15 negatively charged residues, and two charged termini. Replacing the aspartic acids with uncharged asparagines eliminated the repulsive interactions with 18 negatively charged residues + C-terminus (N146 of HcCoh), each interaction enhancing the binding energy by more than 2 kcal/mol (**Fig. 7** and **Tables S10-S11**).

**Figure 7.**
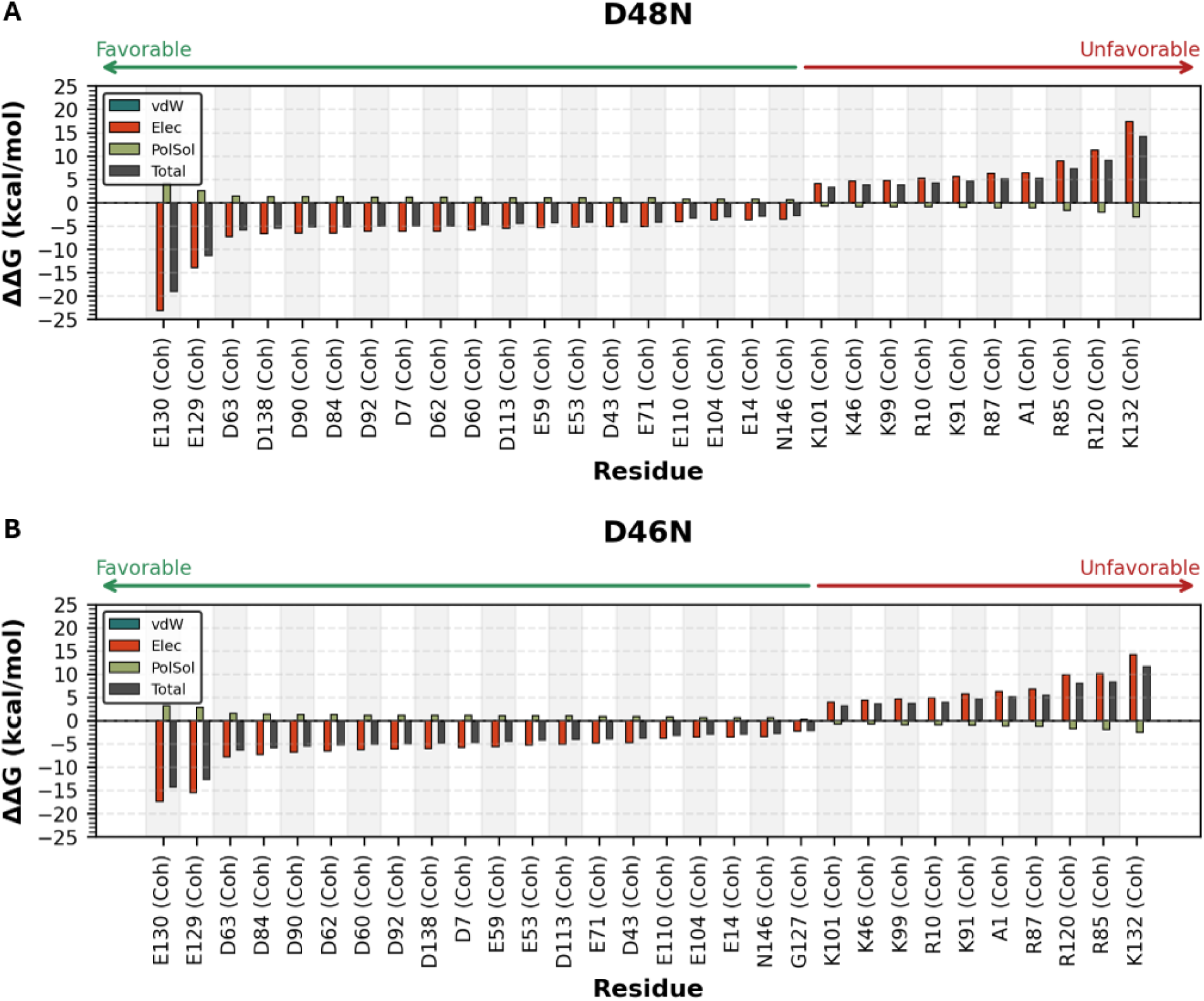
Pairwise interaction energy changes of key residues for HcDoc mutants D46N (HcDoc^m1^) and D48N (HcDoc^m2^). Changes in pairwise interaction energy between mutated HcDoc residues and HcCoh residues for (A) D46N and (B) D48N mutants. ΔΔG = ΔG_mut_ – ΔG_WT_, represents the change in pairwise interaction energy upon mutation, decomposed into van der Waals (vdW), electrostatic (Elec), and polar solvation (PolSol) terms and total (Total) contributions also shown. Only HcCoh residues with |ΔΔG| > 2 kcal/mol are displayed. Negative values indicate improved interactions relative to the wild type (more favorable); positive values indicate worsened interactions (unfavorable).

**Figure 8.**
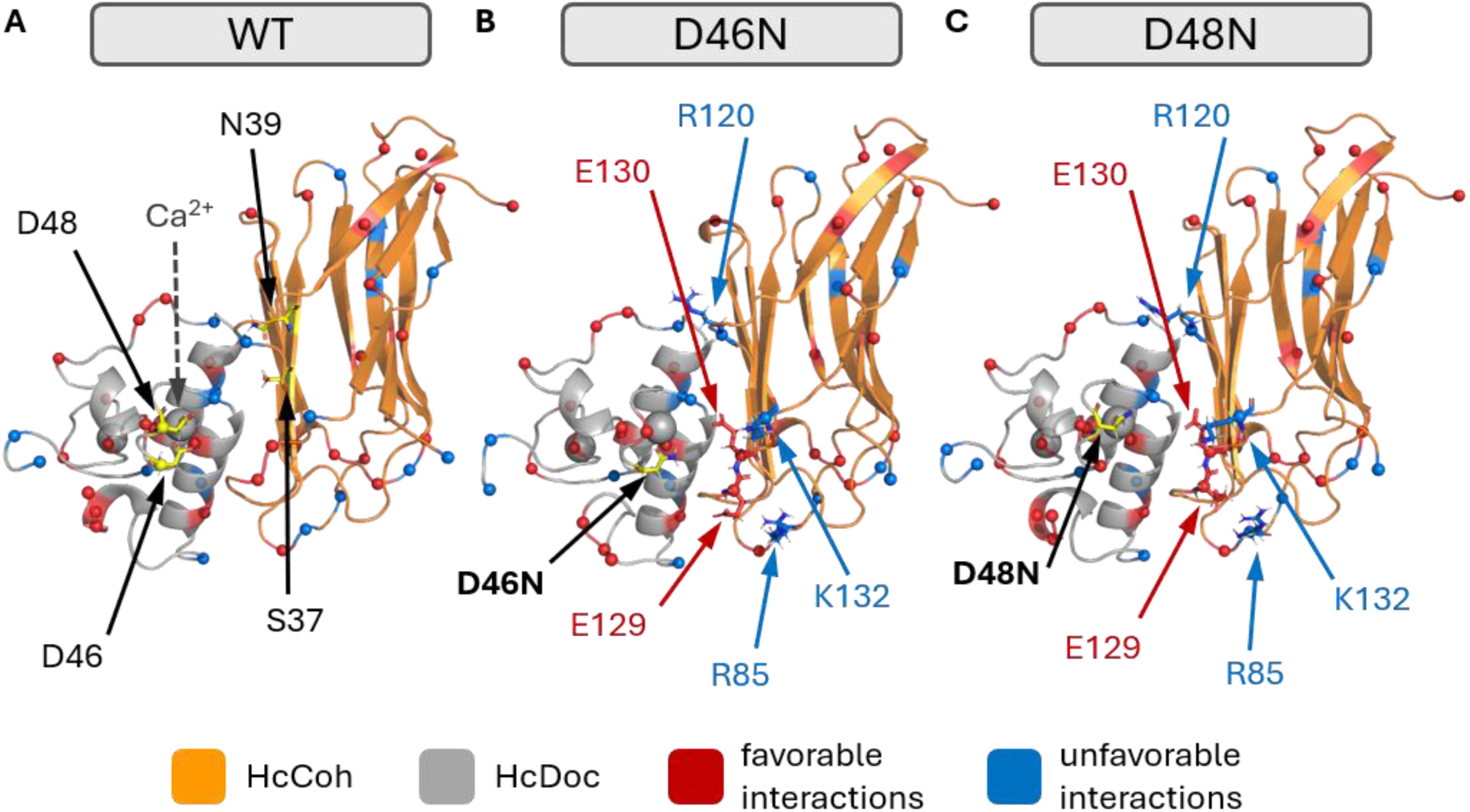
Structural mapping of key residues for HcDoc mutants D46N (HcDoc^m1^) and D48N (HcDoc^m1^). Structural overview of mutation positions and the residues exhibiting the largest changes. A) Distribution of charged residues in the HcCoh-HcDoc WT structure. HcCoh is shown in orange, HcDoc in gray. Mutated residues (D46, D48) are shown as yellow sticks with labeled arrows. The Ca^2+^ ion, coordinated by D46 and D48, is indicated by a gray dashed line. Charged residues are represented as spheres at their Cα atoms (positively charged residues in blue: Lys, Arg; negatively charged in red: Asp, Glu). (B, C) Key pairwise interactions changes for HcDoc B) D46N and C) D48N mutants. Residues with improved interactions (ΔΔG < 0) are shown in red and those with unfavorable interactions (ΔΔG > 0) in blue. In panels (B) and (C), residues with the strongest changes in pairwise interactions are shown as sticks with labeled arrows. Notably, the most favorable interactions (shown in red) involve negatively charged residues (Asp/Glu), while unfavorable interactions (shown in blue) involve positively charged residues (Lys/Arg), consistent with the charge distribution shown in Fig. 7.

The strongest effects (gains of more than 10 kcal/mol) were observed for interactions with E130 and E129 of HcCoh, which are localized in the vicinity of mutated sites (**Fig. 8B-C**). At the same time, attractive interactions of D46N and D48N with 9 positively charged residues and N-terminus (Ala1 of HcCoh) were lost after these substitutions were introduced (**Fig. 7** and **Tables S10-S11**). The strongest loss of attractive interactions was traced to K132 of HcCoh which is in the vicinity of both mutation sites (**Fig. 8B-C**).

Additionally, R85 and R120 swap their relative contributions between the D46N and D48N mutants due to their proximity to the respective mutated residues (**Fig. 8B-C**). In case of D46N, both arginines exhibit a similar change in interaction energy. In contrast, for D48N, the interaction energy change is stronger for R120, which points directly toward the mutated residue D48N. For D46N, besides negatively charged residues, G127 also exceeded the threshold, reflecting repulsion with the backbone carbonyl oxygen of the G127. The change in electrostatic interactions, compensated a bit by opposing solvation energies, was notable for all charged residues and termini. Given that the HcCoh protein contains 19 negatively charged and 10 positively charged sites, which are responsible for favorable and unfavorable changes in the binding energy, respectively, these opposing changes result in a favorable net effect, as reflected in the overall improvement in ΔG_binding_ and the complex stability observed experimentally.

HcCoh mutations are responsible for altering the shape of the interface, improving complementarity between HcCoh and HcDoc. Major favorable changes included: (i) markedly reduced desolvation penalty for burying polar serine and asparagine sidechains upon the complex formation by changing them to hydrophobic residues; (ii) decreased electrostatic repulsion of N39L with K30 and/or R24 and (iii) improved van der Waals packing of S37I or S37V with L28 (**Figs. S13-S14; Tables S12-S13**). This is consistent with observations in related cohesin–dockerin complexes, where increasing the hydrophobicity of the binding interface (T→L substitutions) enhanced affinity, while the reverse substitution of a hydrophobic residue with a polar one (L→S) reduced it, in both cases attributed to changes in the desolvation penalty upon complex formation.^28,88^ On the other hand, major unfavorable changes include decreased electrostatic attraction between N39L and D25 as well as S37I or S37V and R52 (**Figs. S13-S14; Tables S12-S13**). Hydrophobic interactions and electrostatic complementarity drive binding affinity in complex formation. During protein–protein association, solvent molecules must be displaced from the interface. Desolvation of hydrophobic surface patches is thermodynamically favorable due to the hydrophobic effect, whereas desolvation of polar residues incurs an energetic penalty. Therefore, interfaces enriched in hydrophobic residues experience stronger binding affinity, as water displacement is more favorable and requires fewer compensatory interactions.^94–96^

Taken together, our computational analysis indicates that the two HcDoc mutations (D46N and D48N) substantially enhance complex stability by remodeling electrostatic interactions, converting repulsive interactions with negatively charged HcCoh residues into neutral contacts while partially sacrificing attractive interactions with some positively charged ones, with a strongly favorable outcome. In contrast, the HcCoh mutations (S37V/N39L and S37I/N39L) act through a different mechanism, mainly improving hydrophobic packing and reducing solvation penalties at the interface without dramatically altering the overall binding affinity. Previous engineering efforts have shown that dockerins are often more amenable to affinity enhancement than cohesins. For example, mutations within calcium-binding regions or hydrophobic interaction patches have improved the binding affinity of several purified Coh–Doc systems.^28,97^ In contrast, large-scale mutational studies of cohesins have generally identified only modest improvements in binding energetics.^92^ Our results are consistent with these observations: dockerin substitutions produced the strongest stabilizing effects, whereas cohesin substitutions alone had little impact on interaction stability. Importantly, selected combinations of mutant cohesins and dockerins outperformed variants containing engineered dockerins alone, demonstrating that simultaneous optimization of both interaction partners can yield additive benefits. Unlike previous studies, which improved the binding efficiency of Coh–Doc pairs *in vitro* or in bacterial cytoplasm,^28,97^ or increased interaction stability by introducing covalent bonds,^30,87^ our computationally guided approach enhanced the stability of a non-covalent Coh–Doc pair displayed on the surface of a bacterial host. Moreover, surface-display platform presented here also revealed mutation-dependent effects on scaffoldin performance beyond the Coh–Doc interaction itself, including properties that are probably related to better protein production and folding.

## Conclusions

Cohesin–dockerin interactions represent one of the most powerful natural strategies for organizing multienzyme assemblies and have therefore attracted considerable interest as modular connectors for synthetic biology, designer cellulosomes, and engineered whole-cell biocatalysts. However, despite extensive characterization of purified cohesin–dockerin systems, comparatively little is known about their behavior after heterologous expression and surface display in microbial hosts, especially in Gram-negative bacteria. In this study, we established a quantitative *in vivo* platform for investigating thermophilic cohesin–dockerin interactions on the surface of *Pseudomonas putida* EM371 and used it to identify factors governing their performance under application-relevant conditions.

We successfully produced and displayed cohesin and dockerin modules from the thermophilic cellulosome-producing bacterium *Acetivibrio clariflavus* in a mesophilic Gram-negative host and directly compared their behavior with the well-characterized *A. thermocellus* system. Our results demonstrate that two distinct parameters must be considered when evaluating cohesin–dockerin systems for biotechnology: **binding efficiency**, reflecting the extent of complex formation during the assembly phase, and **interaction stability**, reflecting the ability of the assembled complex to remain intact over time. While binding efficiency is commonly evaluated in studies of designer cellulosomes and surface-display systems, interaction stability is often overlooked despite its direct relevance to long-term catalyst performance and enzyme retention.

Using these two metrics, we showed that the temperature used to initiate cohesin–dockerin interactions strongly influences the behavior of thermophilic systems. For both tested pairs, elevated temperatures substantially increased normalized fluorescence and therefore binding efficiency. The effect was particularly pronounced for the *P. putida* cells bearing the Coh-Doc pair from *A. clariflavus*, whose normalized fluorescence increased from 710 ± 259 A.U. after initiating the interaction at 4 °C to 3,264 ± 139 A.U. after incubation at 50 °C. These findings demonstrate that interaction conditions can profoundly affect the assembly of thermophilic Coh–Doc systems and should be optimized before implementing such modules in designer cellulosomes or other modular protein assemblies.

Despite their high binding efficiency, both thermophilic Coh–Doc pairs exhibited lower interaction stability on the bacterial surface than has been reported previously for purified protein systems.^32,73–75^ For the *P. putida* cells bearing the wild-type *A. clariflavus* pair, only 58 ± 11% and 55 ± 13% of the initial normalized fluorescence remained after 1 and 3 h of incubation, respectively. Such loss of surface-bound proteins would likely limit the long-term performance and reusability of Coh–Doc-based whole-cell biocatalysts or other systems. To address this limitation, we combined molecular dynamics simulations and free-energy calculations with experimental validation *in vivo*. Two dockerin variants carrying the D46N (HcDoc^m1^) and D48N (HcDoc^m2^) substitutions and two cohesin variants carrying the S37V/N39L (HcCoh^m1^) and S37I/N39L (HcCoh^m2^) substitutions were constructed and tested.

The computationally designed dockerin variants substantially improved retained normalized fluorescence on the surface of *P. putida*, increasing it from 58 ± 11% reached with the wild-type Hc pair to approximately 89% after 1 h of incubation. Selected mutant cohesin–dockerin combinations further enhanced reporter retention, experimentally validating the computational predictions and demonstrating that rational protein engineering can markedly improve the persistence of thermophilic Coh–Doc assemblies on bacterial surfaces.

Beyond improving interaction stability, the study revealed that mutations can influence the overall performance of the surface-display system beyond the Coh–Doc interface itself. In particular, mutations introduced into the HcCoh module unexpectedly enhanced attachment of the unmodified CtDoc reporter, indicating that protein engineering may affect scaffoldin production, folding, transport, or surface display in addition to Coh–Doc interactions. These findings highlight the importance of evaluating engineered variants within their intended biological context rather than relying solely on purified-protein assays.

Although *P. putida* EM371 proved to be a valuable experimental platform for studying Coh–Doc interactions, our results also revealed a major limitation of this host. Elevated temperatures favored assembly of the thermophilic complexes, yet prolonged incubation at 50 °C caused a rapid loss of cell viability and eventually led to aggregation and deterioration of the system. Consequently, the full potential of thermophilic cohesin–dockerin pairs may not be accessible in mesophilic hosts. Future studies should therefore explore the implementation of similar surface-display platforms in genetically tractable thermophilic bacteria. Particularly promising candidates with available engineering tool repertoire include *Caldimonas thermodepolymerans*,^98^ *Parageobacillus thermoglucosidasius*,^99^ *Bacillus smithii*, *Acetivibrio thermocellus*,^100^ or *Thermoanaerobacter ethanolicus*.^101^ Such hosts could allow both interaction initiation and long-term cultivation at temperatures closer to the physiological optima of thermophilic cellulosome systems, potentially further improving complex formation and stability.

Beyond their relevance for designer cellulosomes, the principles established in this work are applicable to a broad range of surface-display technologies. Stable and tunable Coh–Doc interactions could facilitate the construction of whole-cell biocatalysts with improved enzyme retention and reusability, but also enable alternative applications such as high-throughput screening of antimicrobial-peptide libraries,^102^ development of whole-cell electrochemical biosensors,^103^ and display of enzymes, antibodies, or metal-binding proteins for environmental pollutant removal.^104^ By identifying binding efficiency and interaction stability as two distinct but equally important determinants of Coh–Doc system performance, and by demonstrating that both can be improved through a combination of optimized assembly conditions and computation-guided protein engineering, this study provides a framework for the rational development of next-generation modular cell-surface assemblies for sustainable biotechnology.

## Associated content

### Data Availability Statement

The data that support the findings of this study are available in the Supporting Information of this article. Raw data for all graphs and tables displayed in the manuscript are deposited as Source Data file in the Zenodo repository under DOI identifier 10.5281/zenodo.21851233. The simulation parameters, input files, restarts, dry-simulations, outputs from computational mutagenesis and free energy calculations and other results from the computational analyses are available at: https://doi.org/10.5281/zenodo.21396016.

### Supporting information

**Supplementary methods**: Purification of fluorescent reporter proteins by affinity chromatography, SDS-PAGE and western blot analyses, Fluorescence microscopy, Testing the viability of *P. putida* EM371 at 50 °C, Testing the structural integrity of *P. putida* EM371 after 40 min at 50 °C using light microscopy; **Supplementary results**: Binding efficiency of mutant variants of the *A. clariflavus* (Hc) Coh-Doc pair under different experimental conditions, The expression, display, folding, or functional display of the HcCoh^m1^–CtCoh scaffoldin might be increased; **Supplementary tables**: Bacterial strains used in this study (**Table S1**), List of plasmids used in this study (**Table S2**), List of oligonucleotide primers used in this study (**Table S3**), List of residues at the HcCoh–HcDoc interface identified by FoldX *AnalyzeComplex* (**Table S4**), Results of FoldX PositionScan saturation mutagenesis of HcCoh-HcDoc interface residues (**Table S5**), Per-residue MMGBSA energy decomposition for residues with |ΔG| > 1 kcal/mol (**Table S6**), Results of FoldX *BuildModel* calculations on the 2500-frame conformational ensemble from the wild-type MD simulation (**Table S7**), Results of FoldX AnalyzeComplex calculations on the 2500-frame ensemble of mutant structures generated by FoldX *BuildModel* (**Table S8**), Overall MMPBSA binding free energy (ΔGbinding) calculated using internal dielectric constants of 5 and 10 (**Table S9**), Pairwise MMPBSA energy decomposition calculated using an internal dielectric constant of 5, for residue pairs with |ΔΔG_total| > 2 kcal/mol in the D46N mutant (**Table S10**), Pairwise MMPBSA energy decomposition calculated using an internal dielectric constant of 5, for residue pairs with |ΔΔG_total| > 2 kcal/mol in the D48N mutant (**Table S11**), Pairwise MMPBSA energy decomposition calculated using an internal dielectric constant of 5, for residue pairs with |ΔΔG_total| > 0.25 kcal/mol in the S37I/N39L mutant (**Table S12**), Pairwise MMPBSA energy decomposition calculated using an internal dielectric constant of 5, for residue pairs with |ΔΔG_total| > 0.25 kcal/mol in the S37V/N39L mutant (**Table S13**); **Supplementary figures**: Verification of expression and purification of dockerin-tagged reporter proteins by sodium dodecyl sulfate polyacrylamide gel electrophoresis (**Fig. S1**), Verification of expression of the HcCoh-CtCoh scaffoldin by sodium dodecyl sulfate polyacrylamide gel electrophoresis and western blotting (**Fig. S2**), Photography of cell clumps formed after incubating *P. putida* EM371 at 50°C (**Fig. S3**), Visual verification of the structural integrity of *P. putida* EM371 after 40 min at 50 °C using light microscopy (**Fig. S4**), MD simulations of the wild-type HcCoh-HcDoc complex demonstrate stable and convergent trajectories (**Fig. S5**), Verification of expression and purification of mutant variants of mScarlet-I-HcDoc by sodium dodecyl sulfate polyacrylamide gel electrophoresis (**Fig. S6**), Verification and quantification of expression of the HcCoh^m1^-CtCoh and HcCoh^m2^-CtCoh scaffoldins by sodium dodecyl sulfate polyacrylamide gel electrophoresis and western blotting (**Fig. S7**), Binding efficiency of mutant variants of the *A. clariflavus* (Hc) Coh-Doc pair under different experimental conditions (**Fig. S8**), MD simulations of the D46N HcCoh-HcDoc complex demonstrate stable and convergent trajectories (**Fig. S9**), MD simulations of the D48N HcCoh-HcDoc complex demonstrate stable and convergent trajectories (**Fig. S10**), MD simulations of the S37I/N39L HcCoh-HcDoc complex demonstrate stable and convergent trajectories (**Fig. S11**), MD simulations of the S37V/N39L HcCoh-HcDoc complex demonstrate stable and convergent trajectories (**Fig. S12**), Changes in pairwise interaction energy between mutated HcCoh residues for (a) S37I/N39L and (b) S37V/N39L mutants (**Fig. S13**), Overview on mutation position and strongest changes in pair interactions with mutated residues (**Fig. S14**); **Supplementary sequences**: DNA sequence encoding the Ag43-HcCoh-CtCoh chimeric protein (**Sequence 1**), DNA sequence encoding the mScarlet-HcDoc chimeric protein (**Sequence 2**), DNA sequence encoding the mScarlet-CtDoc chimeric protein (**Sequence 3**) (PDF).

### Author Contributions

**Barbora Jankovičová**: Conceptualization; investigation; methodology; writing – original draft; writing – review and editing; visualization; formal analysis. **Aleksandra Bigos**: Investigation; methodology; writing – review and editing. **Bartłomiej Surpeta**: Investigation; methodology; writing – review and editing. **Miguel Silva**: Investigation; writing – review and editing. **Jan Brezovský**: Conceptualization; investigation; funding acquisition; supervision; writing – original draft; writing – review and editing. **Pavel Dvořák**: Conceptualization; investigation; funding acquisition; supervision; writing – original draft; writing – review and editing. All co-authors read and approved the final version of the manuscript.

### Funding Sources

This project was funded by the Czech Science Foundation (project registration number 25-16845S). B.J. gratefully acknowledges financial support by Brno Ph.D. Talent.

## Acknowledgment

Computations by A.B. and J.B. were performed at the Poznań Supercomputing and Networking Center (Poland).

## Abbreviations

3MB: 3-methylbenzoate
Ag43: antigen 43 autotransporter
CBM: carbohydrate-binding module
CFE: cell-free extract
CFU: colony-forming unit
Coh: cohesin
Ct: Acetivibrio thermocellus
Doc: dockerin
Hc: Acetivibrio clariflavus
MD: molecular dynamics
MMGBSA: Molecular Mechanics / Generalized Born Surface Area
MMPBSA: Molecular Mechanics / Poisson–Boltzmann Surface Area
OD_600_: optical density at 600 nm
RMSD: root-mean-square deviation
RMSF: root-mean-square fluctuation
SLH: surface layer homology
SOEing: splicing by overlap extension
WT: wild type.

## Notes

https://zenodo.org/records/21851234

### Competing Interest Statement

The authors have declared no competing interest.

