## Supplementary Information for "Engineering Binding Efficiency and Interaction Stability of a Thermostable Cohesin–Dockerin Pair on the Bacterial Cell Surface"

**Contents**

**Supplementary methods**

Purification of fluorescent reporter proteins by affinity chromatography

SDS-PAGE and western blot analyses

Fluorescence microscopy

Testing the viability of *P. putida* EM371 at 50 °C

Testing the structural integrity of *P. putida* EM371 after 40 min at 50 °C using light microscopy

**Supplementary results**

Binding efficiency of mutant variants of the *A. clariflavus* (Hc) Coh-Doc pair under different experimental conditions

The expression, display, folding, or functional display of the $HcCoh^{m1}$–CtCoh scaffoldin might be increased

**Supplementary tables**

**Table S1.** Bacterial strains used in this study.

**Table S2.** List of plasmids used in this study.

**Table S3.** List of oligonucleotide primers used in this study.

**Table S4.** List of residues at the HcCoh–HcDoc interface identified by FoldX *AnalyzeComplex.*

**Table S5.** Results of FoldX PositionScan saturation mutagenesis of HcCoh-HcDoc interface residues.

**Table S6.** Per-residue MMGBSA energy decomposition for residues with |ΔG| > 1 kcal/mol.

**Table S7.** Results of FoldX *BuildModel* calculations on the 2500-frame conformational ensemble from the wild-type MD simulation.

**Table S8.** Results of FoldX AnalyzeComplex calculations on the 2500-frame ensemble of mutant structures generated by FoldX *BuildModel*.

**Table S9.** Overall MMPBSA binding free energy (ΔGbinding) calculated using internal dielectric constants of 5 and 10.

**Table S10.** Pairwise MMPBSA energy decomposition calculated using an internal dielectric constant of 5, for residue pairs with |ΔΔG_total| > 2 kcal/mol in the D46N mutant.

**Table S11.** Pairwise MMPBSA energy decomposition calculated using an internal dielectric constant of 5, for residue pairs with |ΔΔG_total| > 2 kcal/mol in the D48N mutant.

**Table S12.** Pairwise MMPBSA energy decomposition calculated using an internal dielectric constant of 5, for residue pairs with |ΔΔG_total| > 0.25 kcal/mol in the S37I/N39L mutant.

**Table S13.** Pairwise MMPBSA energy decomposition calculated using an internal dielectric constant of 5, for residue pairs with |ΔΔG_total| > 0.25 kcal/mol in the S37V/N39L mutant.

**Supplementary figures**

**Figure S1.** Verification of expression and purification of dockerin-tagged reporter proteins by sodium dodecyl sulfate polyacrylamide gel electrophoresis.

**Figure S2.** Verification of expression of the HcCoh-CtCoh scaffoldin by sodium dodecyl sulfate polyacrylamide gel electrophoresis and western blotting.

**Figure S3.** Photography of cell clumps formed after incubating *P. putida* EM371 at 50°C.

**Figure S4.** Visual verification of the structural integrity of *P. putida* EM371 after 40 min at 50 °C using light microscopy.

**Figure S5.** MD simulations of the wild-type HcCoh-HcDoc complex demonstrate stable and convergent trajectories.

**Figure S6.** Verification of expression and purification of mutant variants of mScarlet-I-HcDoc by sodium dodecyl sulfate polyacrylamide gel electrophoresis.

**Figure S7.** Verification and quantification of expression of the HcCoh$^{m1}$-CtCoh and HcCoh$^{m2}$-CtCoh scaffoldins by sodium dodecyl sulfate polyacrylamide gel electrophoresis and western blotting.

**Figure S8.** Binding efficiency of mutant variants of the *A. clariflavus* (Hc) Coh-Doc pair under different experimental conditions.

**Figure S9.** MD simulations of the D46N HcCoh-HcDoc complex demonstrate stable and convergent trajectories.

**Figure S10.** MD simulations of the D48N HcCoh-HcDoc complex demonstrate stable and convergent trajectories.

**Figure S11.** MD simulations of the S37I/N39L HcCoh-HcDoc complex demonstrate stable and convergent trajectories.

**Figure S12.** MD simulations of the S37V/N39L HcCoh-HcDoc complex demonstrate stable and convergent trajectories.

**Figure S13.** Changes in pairwise interaction energy between mutated HcCoh residues for (a) S37I/N39L and (b) S37V/N39L mutants.

**Figure S14.** Overview on mutation position and strongest changes in pair interactions with mutated residues.

**Supplementary sequences**

**Sequence 1.** DNA sequence encoding the Ag43-HcCoh-CtCoh chimeric protein.

**Sequence 2.** DNA sequence encoding the mScarlet-HcDoc chimeric protein.

**Sequence 3.** DNA sequence encoding the mScarlet-CtDoc chimeric protein.

**Supplementary methods**

<u>Purification of fluorescent reporter proteins by affinity chromatography</u>

Precultures of *E. coli* BL21-Gold (DE3) carrying the pET-21b+_*mScarlet-ctDoc-his*, pET-21b+_*mScarlet-hcDoc-his*, pET-21b+_*mScarlet-hcDoc*$^{m1}$*-his,* or pET-21b+_*mScarlet-hcDoc*$^{m2}$*-his* were grown in 10 mL LB with the appropriate antibiotics and 10 mM $CaCl_2$ at 37°C and 215 rpm (NB-205 incubator; N-BIOTEK) for 16 hours. Main cultures (100 mL LB) containing 10 mM $CaCl_2$ and the appropriate antibiotics were inoculated to an initial $OD_{600}$ of 0.05 and grown at 37°C and 215 rpm (NB-205) for 2.5 hours. I0 samples were collected for SDS-PAGE (2 mL of culture was centrifuged at 4,000 rpm; 10 min; 4°C). Reporter protein production was induced with IPTG (50 µM final concentration). Cultivation continued at 20°C and 215 rpm for 20 h. Samples I for SDS-PAGE were collected (2 mL of culture was centrifuged at 4,000 rpm; 10 min; 4°C). The cells from the remaining culture were collected by centrifugation (4°C; 2,000 g; 15 min) and washed with 50 mL of ice-cold purification buffer A (TBS-Ca-T buffer: 25 mM Tris-Cl, 137 mM NaCl, 2,7 mM KCl, 10 mM $CaCl_2$, 0.05% Tween 20; 5 mM imidazole). After washing, the cells were collected again (4°C; 2,000 g; 15 min). Cell-free extracts (CFE) were prepared from the pellets using the B-PER with Enzymes Bacterial Protein Extraction Kit (Thermo Fisher Scientific), along with protease inhibitor cocktail tablets (cOmplete Mini, EDTA-free; Roche). The reaction was incubated at room temperature for 15 min with slow mixing, then centrifuged (21,000 g; 30 min; 4°C).

Ni-NTA agarose chromatography columns (Poly-Prep Chromatography Columns) were equilibrated with 4× matrix volume of purification buffer A. CFEs were loaded onto the columns and incubated with the matrix at 4°C for 30 min with slow rotation to promote binding. The waste fraction was collected, and the column was washed with 2× the matrix volume of purification buffer A and 5× the matrix volume of purification buffer B (TBS-Ca-T buffer, 25 mM imidazole). The proteins bound to the matrix were eluted with 5× the matrix volume of purification buffer C (TBS-Ca-T buffer, 500 mM imidazole). The first 7.5 mL of flow after the addition of purification buffer C was collected (5 elution fractions of 1.5 mL each).

The elution fractions' total protein concentrations were measured at 595 nm in a transparent 96-well microtiter plate (Thermo Fisher Scientific) using Bradford reagent (Sigma-Aldrich) and the Infinite M Plex reader (Tecan). The two fractions with the highest protein concentrations were pooled and exchanged into imidazole-free TBS-Ca-T buffer by repeated concentration (centrifugation at 7,500 g; times varied between 3 and 20 min) and dilution using Amicon Ultra-4 centrifugal filter units (10 kDa cutoff). The total protein concentrations of the exchanged buffer samples were measured as described above. Purified protein samples, as well as samples I0 and I, were analyzed using SDS-PAGE (**Supplementary methods**). The purified proteins were mixed with TBS-Ca-T buffer and glycerol (final concentration 20% v/v) to a final concentration of 1 g $L^{-1}$ based on the purity of the protein of interest, and stored at -20°C.

SDS-PAGE and western blot analyses

Samples I0, I, and purified proteins were prepared as described in the Experimental Section of the main text (**Purification of reporter proteins by affinity chromatography; Detection of scaffoldin expression in *P. putida***). Samples I0 and I were lysed using B-PERwith Enzymes Bacterial Protein Extraction Kit. The lysates and purified proteins were mixed with SDS sample loading buffer (5x concentrated: 0.2 mL of β-mercaptoethanol, 0.8 mL of 87% glycerol, 0.8 mL of 10% SDS (w/v), 1.2 mL of Milli-Q $H_2O$, 0.5 mL of Tris-HCl (0.5 M, pH 6.8), a pinch of Bromphenol blue) and Milli-Q water. Samples of 2 or 4 µg of protein were loaded onto the gel for purified protein or lysate, respectively. For western blotting, the protein concentration was doubled to enhance the sample signal on the membrane. The samples were boiled at 95°C for 10 min and loaded in SDS-PAGE gels (4% stacking and 7.5% or 12% resolving for scaffoldin or reporter protein samples, respectively). Electrophoresis was run at 125 V for 80 min in SDS-PAGE running buffer (14.4 g $L^{-1}$ glycine, 1 g $L^{-1}$ SDS, 3 g $L^{-1}$ Tris base). The gels were stained with Quick Coomassie Blue (Serva), photographed, and analyzed using a Densitometer GS-800 (Bio-Rad).

For western blotting, the gels with samples of higher protein content, along with all components of the transfer apparatus (Mini-PROTEAN Tetra Cell) and the membrane (Nitrocellulose membrane, 0.45 μ, 7.9×10.5 cm), were washed in an ice-cold western blot transfer buffer (3 g $L^{-1}$ Tris base, 14.5 g $L^{-1}$ glycine, 200 ml $L^{-1}$ methanol, and 5 ml $L^{-1}$ 10% SDS (w/v)). The transfer was conducted at 375 mA for 1.5 h. Subsequently, the membrane was washed overnight in Incubation buffer A (30 g $L^{-1}$ dry milk, 1 ml $L^{-1}$ Tween 20, 2.68 g $L^{-1}$ $Na_2HPO_4{\cdot}7H_2O$, 0.2 g $L^{-1}$ KCl, 0.24 g $L^{-1}$ $KH_2PO_4$, 8 g $L^{-1}$ NaCl, pH 7.4) without agitation. Following this, 5 ml of Incubation buffer A was mixed with 5 μl of 6× His mAb/HRP Conjugate antibody (TaKaRa), added to the membrane, and incubated for 2 h at room temperature with minimal agitation. The membrane was washed four times for 4 min each in the western blot wash buffer (8.8 g $L^{-1}$ NaCl, 2.4 g $L^{-1}$ Tris, 500 μl $L^{-1}$ Tween 20). Clarity Max ECL Western Blotting Substrate (Bio-Rad) was used to detect the antibody. The membrane was photographed using the Fusion Solo S machine (Vilber).

Fluorescence microscopy

Cells with attached reporter proteins were prepared as described in **Testing the binding efficiency of the Coh-Doc pairs**. Sample preparation and fluorescence microscopy were carried out as previously described by Burýšková and colleagues.[1] Briefly, the cells were fixed with 8% formaldehyde and 0.25% glutaraldehyde solution (1:1 v/v) for 30 min in Cellview cell culture slides (polystyrene, 75/25 mm, glass bottom; Greiner). The fixed samples were washed with Milli-Q $H_2O$ and covered with Vectashield Antifade Mounting Medium (Vector Laboratories). The cells were visualized using ZEISS Elyra 7 with lattice SIM mounted with the ZEISS objective Plan-Apochromat 63x/1.4 Oil DIC M27 and captured in pixel size 60 nm for channels 488 nm and 561 nm in 3D. Data was processed with Zeiss software SIM2 and visualized as an orthogonal projection.

Testing the viability of *P. putida* EM371 at 50 °C

The *P. putida* EM371 cells carrying pSEVA238_*ag43-hcCoh-ctCoh-his* were cultured and prepared as described in **Testing the binding efficiency of the Coh-Doc pairs** in the main text. The resulting samples (0.5 mL, $OD_{600}$ = 10.0) were incubated at 50°C for 0, 10, 20, 40, or 60 min. Samples (100 µl) of several dilutions from each time point were plated on LB agar plates containing the appropriate antibiotics and incubated overnight at 30 °C. Colony-forming units (CFU) were quantified, and CFU/ml were determined for a sample of $OD_{600}$ = 10.0.

Testing the structural integrity of *P. putida* EM371 after 40 min at 50 °C using light microscopy

The *P. putida* EM371 cells carrying pSEVA238_*ag43-hcCoh-ctCoh-his* were cultured and prepared as described in **Testing the binding efficiency of the Coh-Doc pairs** in the main text. The resulting samples (0.5 mL, $OD_{600}$ = 10.0) were incubated at 50°C for 40 min. Samples of 2 µl were diluted in 18 µl of TBS-Ca-T and mixed with 2 µl of methylene blue directly on a microscope slide. They were covered with a coverslip and observed with the Olympus BX50 upright microscope (Olympus Corporation, Japan) using a UPlanFl 100x/1.30 Oil Ph3 objective under phase contrast.

**Supplementary results**

<u>Binding efficiency of mutant variants of the *A. clariflavus* (Hc) Coh-Doc pair under different experimental conditions</u>

We tested whether the mutant Hc Coh–Doc variants retained their ability to form functional complexes under the interaction conditions used for the wild-type pairs (**Fig. 5A, Fig. S8A–D**). At 4 °C for 13 h, none of the mutant combinations differed significantly from the wild-type Hc pair in normalized fluorescence values ($p > 0.05$; **Fig. S8A**). At 30 °C for 1 h, however, normalized fluorescence depended primarily on the dockerin variant used: wild-type HcDoc produced the strongest signals, whereas $HcDoc^{m2}$ yielded significantly lower fluorescence than $HcDoc^{m1}$ in combination with $HcCoh^{m1}$ and $HcCoh^{m2}$ ($p = 6.23 \times 10^{-4}$ and $2.31 \times 10^{-2}$, respectively; **Fig. S8B**). After interaction initiation at 50 °C for 40 min, *P. putida* cells with most mutant combinations reached normalized fluorescence levels comparable to the cells with the wild-type pair. The only exception was cells with HcCoh–CtCoh + $HcDoc^{m2}$, which showed significantly lower fluorescence than the cells with the wild-type pair ($p = 0.045$; **Fig. S8C**). After 1 h at 50 °C, the reduced performance of $HcDoc^{m2}$ became more pronounced, as this variant produced significantly lower normalized fluorescence than wild-type HcDoc with all tested cohesins ($p = 1.11 \times 10^{-4}$, $3.49 \times 10^{-4}$, and $1.31 \times 10^{-5}$; **Fig. S8D**). In contrast, $HcDoc^{m1}$ generally yielded normalized fluorescence values comparable to wild-type HcDoc, with the exception of its combination with $HcCoh^{m2}$–CtCoh ($p = 0.015$).

<u>The expression, display, folding, or functional display of the $HcCoh^{m1}$–CtCoh scaffoldin might be increased</u>

Unexpectedly, the $HcCoh^{m1}$–CtCoh scaffoldin increased binding of the unmodified CtDoc reporter under three of the four tested conditions when compared with the wild-type scaffoldin ($p = 0.015$, $4.82 \times 10^{-3}$, 0.023, and 0.055 for 4 °C/13 h, 30 °C/1 h, 50 °C/40 min, and 50 °C/1 h, respectively; **Fig. S8E**). Because the Ct Coh–Doc pair itself was not mutated, this improvement cannot be attributed to altered CtCoh–

CtDoc affinity. Instead, it likely reflects changes in the overall performance of the surface-displayed scaffoldin, including expression, transport across the cell envelope, surface display, folding, or their combined effects. Although HcCoh$^{m1}$–CtCoh showed a modest increase in scaffoldin abundance compared with the wild type, this difference was not statistically significant ($p = 0.075$; **Fig. S7C**), indicating that increased production alone is unlikely to explain the higher CtDoc attachment. Thus, even two substitutions in the HcCoh module (S37V/N39L) can indirectly improve the functional performance of the adjacent CtCoh module, most likely through cumulative effects on scaffoldin production, display, and/or folding.

### Supplementary tables

**Table S1: Bacterial strains used in this study.**

| Strain | Characteristics | Source or reference |
|---|---|---|
| *Escherichia coli* CC118 | Cloning host*: Δ(ara-leu) araD Δlac X174 galE galK phoA thiE1 rpoB(RifR) argE(Am) recA1* | Manoil *et* Beckwith, 1985[2] |
| *Escherichia coli* BL21-Gold (DE3) | Expression host, *E. coli* B derivative: F– *ompT hsdS(*rB- mB- *) dcm+ Tetr gal λ(*DE3 *[lacIQ lacUV5-T7 gene 1 ind1 sam7 nin5]) endA* Hte | Agilent Technologies, USA |
| *Pseudomonas putida* EM371 | Derivative of strain *P. putida* KT2440: Δprophages1,2,3,4 ΔTn7 ΔendA1 ΔendA2 ΔhsdRMS Δflagellum ΔTn4652 Martínez-García et al. 4 EM371 Derivative of strain P. putida KT2440: Δprophage4 ΔTn7 Δflagellum Δpili Δcurli Δmotility proteins Δalginate biosynthesis Δtwitching motility protein Δsurface adhesion protein Δcellulose synthesis Δouter membrane lipoprotein Δglycosyl transferase | Martínez-García *et* al., 2020[3] |

**Table S2: List of plasmids used in this study.**

| Plasmid | Description | Source or reference |
|---|---|---|
| pET-*21b*+ | Expression vector: *ori*(pBR322, f1), T7 promoter and terminator, *lacI* $Amp^R$ | Merck, USA |
| pET-21b+_*his-mScarlet-hcDoc* | pET21b+ with *his-mScarlet-I-hcDoc* gene encoding red fluorescent protein (mScarlet-I[4]) fused with a dockerin originating from *Acetivibrio clariflavus* (formerly *Hungateiclostridium clariflavum* = Hc), with an N-terminal 6xHis tag (*Nde*I, *Hind*III) | This study |
| pET-21b+_*mScarlet-ctDoc-his* | pET21b+ with *mScarlet-I-ctDoc-his* gene encoding mScarlet-I fused with a dockerin originating from *Acativibrio thermocellus* (formerly *Clostridium thermocellum* = Ct), with a C-terminal 6xHis tag (*Nde*I, *Hind*III) | This study |
| pET-21b+_*mScarlet-hcDoc-his* | pET21b+ with *mScarlet-I-hcDoc-his* gene encoding mScarlet-I fused with a dockerin originating from Hc, with a C-terminal 6xHis tag (*Nde*I, *Xho*I) | This study |
| pET-21b+_*mScarlet-hcDoc$^{m1}$-his* | pET21b+ with *mScarlet-I-hcDoc-his* gene encoding mScarlet-I fused with a dockerin originating from Hc with a D46N substitution, with a C-terminal 6xHis tag (*Nde*I, *Xho*I) | This study |
| pET-21b+_*mScarlet-hcDoc$^{m2}$-his* | pET21b+ with *his-mScarlet-I-hcDoc* gene encoding mScarlet-I fused with a dockerin originating from Hc with a D48N substitution, with a C-terminal 6xHis tag (*Nde*I, *Xho*I) | This study |
| pMRE-Tn7-145_*mScarlet-I* | Delivery vector: *ori*(f1), *araC rop101* promoter araBAD $Gm^R$ $Cm^R$ $Amp^R$ containing the coding sequence of mScarlet-I | Schlechter *et al.,* 2018[4] |
| pEX-A128_*l-hcDoc-hcCoh* | Delivery vector: *ori* (pUC), $Amp^R$ sequences encoding a synthetic linker, HcDoc, and HcCoh codon-optimised for production in *P. putida* | Eurofins Scientific |
| pSEVA238 | Expression vector: *oriV*(pBBR1) *xylS-Pm neo*, $Km^R$ | Silva-Rocha *et al.,* 2013[5] |

| Plasmid | Description | Source or reference |
|---|---|---|
| pSEVA238b | Derivative of pSEVA238 with synthetic RBS and adjacent *Nde*I site for subcloning of genes to be expressed | Dvořák *et* De Lorenzo, 2018[6] |
| pSEVA238b_*ag43AT* | pSEVA238b with gene encoding C-terminal part (487 AA) of adhesin Ag43 autotransporter from *E. coli* with original 52 AA leader sequence and inserted polylinker (*Nde*I, *Hind*III) | Dvořák *et al.,* 2020[7] |
| pSEVA238b_*ag43-acCoh-ctCoh-his* | pSEVA238b_*ag43AT* with gene encoding *acCoh-ctCoh* (cohesins originating from Ac and Ct, respectively) scaffoldin codon-optimized for expression in *P. putida* KT2440, with a C-terminal 6xHis tag (*Xho*I, *Bam*HI) | Dvořák *et al.,* 2020[7] |
| pSEVA238b_*ag43_hcCoh-ctCoh-his* | pSEVA238b_*ag43AT* with gene encoding *hcCoh-ctCoh* (cohesins originating from Hc and Ct, respectively) scaffoldin codon-optimized for expression in *P. putida* KT2440, with a C-terminal 6xHis tag (*Xho*I, *Bam*HI) | This study |
| pSEVA238b_*ag43_hcCoh$^{m1}$-ctCoh-his* | pSEVA238b_*ag43AT* with gene encoding *hcCoh$^{m1}$-ctCoh* (cohesins originating from Hc (with substitutions S37V and N39L) and Ct, respectively) scaffoldin codon-optimized for expression in *P. putida* KT2440, with a C-terminal 6xHis tag (*Xho*I, *Bam*HI) | This study |
| pSEVA238b_*ag43_hcCoh$^{m2}$-ctCoh-his* | pSEVA238b_*ag43AT* with gene encoding *hcCoh$^{m2}$-ctCoh* (cohesins originating from Hc (with substitutions S37I and N39L) and Ct, respectively) scaffoldin codon-optimized for expression in *P. putida* KT2440, with a C-terminal 6xHis tag (*Xho*I, *Bam*HI) | This study |

**Table S3: List of oligonucleotide primers used in this study.**

| Number | Name | Sequence (5'->3') | Purpose description |
|---|---|---|---|
| BH10 | MCS seq fw | GTCCGGCGTAGAGG | Colony PCR and sequencing of cargo in pET-21b(+) |
| BH11 | MCS seq rv | GGGGTTATGCTAGTTATTGC | Colony PCR and sequencing of cargo in pET-21b(+) |
| BH12 | *mScarlet-I* fw1 | ATTCATATG**CACCATCACCATCACCAT**GTGAGCAAGGGCGA | PCR amplification of *mScarlet-I* with an addition of an N-terminal 6x Histag |
| BH13 | *mScarlet-I* rv | CTTGTACAGCTCGTCCATG | PCR amplification of *mScarlet-I* |
| BH14 | *mScarlet-I* fw2 | ATTCATATGCACCATCACC | Primer for the second step of the soePCR |
| BH15 | linker *hcDoc* fw | CATGGACGAGCTGTACAAGAACCCCAACCCGAAC | Amplification of *hcDoc* with linker and following connection to *mScarlet-I* |
| BH16 | *hcDoc* rv1 | TTAAAGCTTTCATTGTTCTTCCACCGGG | Amplification of *hcDoc* and addition of a stop codon and the *Hind*III restriction site |
| BH17 | *hcDoc* rv2 | TTAAAGCTTTCATTGTTCTTCCAC | Amplification of *hcDoc* and addition of a stop codon and the *Hind*III restriction site |
| BH18 | *hcCoh* fw | ATTCTCGAGGCCGGTCAGCTGC | Amplification of *hcCoh* with addition of *Xho*I (-> cloning to *ag43*) |
| BH19 | *hcCoh* rv | GTTGCTGCCCACCA | Amplification of *hcCoh* |
| BH20 | *scaf* fw | TGGTGGGCAGCAACCGACCCCGACTC | Amplification of linker and *ctCoh* and following connection to *hcCoh* |
| BH21 | *scaf* rv | TTAGGATCCGTGATGGTGG | Amplification of *acCoh-ctCoh-His* |

Underlined: restriction sites, **bold**: sequence encoding 6x His tag

| Number | Name | Sequence (5'->3') | Purpose description |
|---|---|---|---|
| BH24 | *mScarlet-I* fw NO His | ATTCATATGGTGAGCAAGGGC | Amplification of *mScarlet-I* without the addition of the N-terminal 6xHis tag |
| BH25 | *ag43AT*-display_seq | CGTTCACCAGGTTCAGGATGG | Sequencing of cargo in pSEVA238b_*Ag43AT* |
| BH26 | *mScarlet-I*_NOHis_fw_new | AAACATATGGTGAGCAAGGGCG | Amplification of *mScarlet-I-hcDoc* from the *his-mScarlet-I-hcDoc* template |
| BH27 | *mScarlet-I*_C-ter_His_XhoI | TTTCTCGAGTTGTTCTTCCACCGGG | Addition of the *Xho*I restriction site to the *His-mScarlet-I-hcDoc* sequence for insertion into pET21b+ and connection to the 6xHis tag present in the vector |
| pBH28 | *mScarlet_C-ter_His_HindIII* | TATAAGCTTTCAGTGGTGGTGGTGGTGGTGTTGTTCTTCCACCGGG | Addition of the C-terminal 6xHis to the *mScarlet-hcDoc* sequence |
| pBH38 | *mSc 568 rv EBl* | TGGGGTCGGGGTTGGGTTCGGGTTGGGGTTCTTGTACAGCTCGTCCATG | Amplification of *mScarlet* from pMRE-Tn7-145_*mScarlet-I* and addition of EB linker to its 3' end |
| pBH39 | *ctDoc 775 fw EBl* | AACCCCGACCCCAACCCCAGGCAGCGGCGGGACGCCC | Amplification of *ctDoc* from pET-21b+_*his-bglC-ctDoc-his* and addition of EB linker to its 5' end |
| pBH43 | *hcDoc D48N rv* | GATGTTGCCGTTGCCATCCACG | Introduction of the D48N substitution to *hcDoc* to prepare *hcDoc*$^{m2}$ |
| pBH44 | *hcDoc D48N fw* | GATGGCAACGGCAACATCCGTATC | Introduction of the D48N substitution to *hcDoc* to prepare *hcDoc*$^{m2}$ |
| pBH45 | *hcDoc D46N rv* | GTCGCCGTTCACGTCGGC | Introduction of the D46N substitution to *hcDoc* to prepare *hcDoc*$^{m1}$ |

Underlined: restriction sites, **bold**: sequence encoding 6x His tag

| Number | Name | Sequence (5'->3') | Purpose description |
|---|---|---|---|
| pBH46 | *hcDoc D46N fw* | CGACGTGAACGGCGACGG | Introduction of the D46N substitution to *hcDoc* to prepare $hcDoc^{m1}$ |
| pBH47 | *hcCoh S37V N39L fw* | CCCTGGTGCTGCTGCTGAACTTC | Introduction of the S37V N39L substitution to *hcCoh* to prepare $hcCoh^{m1}$ |
| pBH48 | *hcCoh S37V N39L rv* | CAGCAGCAGCACCAGGGCGTAG | Introduction of the S37V N39L substitution to *hcCoh* to prepare $hcCoh^{m1}$ |
| pBH49 | *hcCoh S37I N39L fw* | CCCTGATCCTGCTGCTGAACTTCG | Introduction of the S37I N39L substitution to *hcCoh* to prepare $hcCoh^{m2}$ |
| pBH50 | *hcCoh S37I N39L rv* | CAGCAGCAGGATCAGGGCG | Introduction of the S37I N39L substitution to *hcCoh* to prepare $hcCoh^{m2}$ |
| p9 | *PS2 new* | CGGCAACCGAGCGTTC | Universal sequencing and colony PCR primer that anneals to the T0 terminator in pSEVA238b |
| p32 | *Pm promoter* | ATGGCTATCTCTAGTAAGG CCTAC | Universal sequencing and colony PCR primer that anneals to the *Pm* promoter in pSEVA238b |

Underlined: restriction sites, **bold**: sequence encoding 6x His tag

**Table S4: List of residues at the HcCoh–HcDoc interface identified by FoldX *AnalyzeComplex***. The analysis was performed on 2,500 simulation frames from wild-type conformational ensemble. Residues present in more than 50% of frames (i.e., count > 1,250 out of 2,500 frames) were selected as candidate mutation sites for further mutagenesis screening.

| Protein | Residue | Count | Occurrence_% |
|---|---|---|---|
| HcDoc | LEU 58 | 2500 | **100.0** |
| HcDoc | LEU 28 | 2500 | **100.0** |
| HcDoc | ASN 54 | 2500 | **100.0** |
| HcCoh | PHE 67 | 2500 | **100.0** |
| HcCoh | SER 122 | 2500 | **100.0** |
| HcDoc | LYS 30 | 2500 | **100.0** |
| HcCoh | MET 80 | 2500 | **100.0** |
| HcDoc | ILE 53 | 2500 | **100.0** |
| HcCoh | ALA 65 | 2500 | **100.0** |
| HcCoh | ALA 83 | 2500 | **100.0** |
| HcCoh | TYR 34 | 2500 | **100.0** |
| HcCoh | TYR 124 | 2500 | **100.0** |
| HcDoc | VAL 57 | 2500 | **100.0** |
| HcCoh | ALA 35 | 2500 | **100.0** |
| HcDoc | VAL 27 | 2500 | **100.0** |
| HcDoc | ARG 24 | 2500 | **100.0** |
| HcCoh | SER 78 | 2500 | **100.0** |
| HcCoh | THR 128 | 2500 | **100.0** |
| HcDoc | ARG 52 | 2500 | **100.0** |
| HcDoc | ARG 60 | 2500 | **100.0** |
| HcCoh | SER 76 | 2500 | **100.0** |
| HcCoh | SER 37 | 2500 | **100.0** |
| HcDoc | ASP 61 | 2499 | **100.0** |
| HcDoc | ASP 46 | 2498 | **99.9** |
| HcDoc | SER 56 | 2497 | **99.9** |
| HcCoh | ARG 120 | 2490 | **99.6** |
| HcDoc | PRO 71 | 2481 | **99.2** |
| HcCoh | PRO 82 | 2467 | **98.7** |
| HcCoh | LEU 36 | 2461 | **98.4** |
| HcDoc | LYS 66 | 2446 | **97.8** |
| HcDoc | LEU 64 | 2434 | **97.4** |
| HcCoh | PHE 74 | 2434 | **97.4** |
| HcCoh | ASN 39 | 2416 | **96.6** |
| HcCoh | GLU 130 | 2379 | **95.2** |
| HcCoh | MET 77 | 2277 | **91.1** |
| HcDoc | ASP 48 | 2226 | **89.0** |
| HcCoh | LEU 66 | 2181 | **87.2** |
| HcCoh | ALA 81 | 2149 | **86.0** |
| HcDoc | TYR 26 | 1973 | **78.9** |
| HcDoc | ASP 32 | 1439 | **57.6** |
| HcDoc | ASN 50 | 1424 | **57.0** |
| HcDoc | ALA 20 | 1367 | **54.7** |

HcCoh – cohesin; HcDoc – dockerin

| Protein | Residue | Count | Occurrence_% |
| --- | --- | --- | --- |
| HcCoh | ASN 69 | 1147 | **45.9** |
| HcCoh | ASP 63 | 915 | **36.6** |
| HcCoh | LYS 132 | 894 | **35.8** |
| HcCoh | ARG 85 | 803 | **32.1** |
| HcCoh | PHE 79 | 798 | **31.9** |
| HcCoh | ASP 84 | 731 | **29.2** |
| HcCoh | PHE 64 | 656 | **26.2** |
| HcDoc | ASP 25 | 650 | **26.0** |
| HcDoc | ASP 55 | 597 | **23.9** |
| HcDoc | ILE 67 | 429 | **17.2** |
| HcCoh | SER 119 | 357 | **14.3** |
| HcDoc | ILE 51 | 323 | **12.9** |
| HcDoc | VAL 21 | 197 | **7.9** |
| HcDoc | ILE 31 | 168 | **6.7** |
| HcCoh | HIS 72 | 104 | **4.2** |
| HcCoh | THR 121 | 103 | **4.1** |
| HcDoc | GLN 75 | 100 | **4.0** |
| HcCoh | THR 126 | 98 | **3.9** |
| HcDoc | ILE 17 | 70 | **2.8** |
| HcDoc | MET 40 | 42 | **1.7** |
| HcCoh | TYR 118 | 34 | **1.4** |
| HcCoh | ASP 62 | 11 | **0.4** |
| HcCoh | GLU 129 | 9 | **0.4** |
| HcDoc | GLU 74 | 7 | **0.3** |
| HcDoc | GLU 33 | 5 | **0.2** |
| HcCoh | TYR 136 | 3 | **0.1** |
| HcCoh | ASN 117 | 2 | **0.1** |
| HcCoh | GLU 71 | 1 | **0.0** |

HcCoh – cohesin; HcDoc – dockerin

**Table S5: Results of FoldX *PositionScan* saturation mutagenesis of HcCoh-HcDoc interface residues.** Each candidate mutation site identified in the interface analysis was systematically substituted with all 20 amino acids. The table lists substitutions with $\Delta\Delta G < -1$ kcal/mol ($\Delta\Delta G = \Delta G_{Mut} - \Delta G_{WT}$) indicating a predicted stabilizing effect relative to the wild-type. Values represent the mean and standard deviation calculated over the 2,500-frame conformational ensemble. Substitutions excluded based on additional risk criteria (described in the **Computation-guided mutagenesis identifies substitutions with the potential to improve the interaction stability of the Hc Coh-Doc pair** section of the main text) are indicated in the last column. The 5 candidate substitutions (across 4 mutation sites) selected for combinatorial library construction are highlighted in bold.

| No. | Protein | Mutation | ΔΔG_mean (kcal/mol) | ΔΔG_std (kcal/mol) | Exlusion criteria |
|---|---|---|---|---|---|
| 1 | **HcDoc** | **D46N** | **-2.86** | **1.17** | **-** |
| 2 | HcDoc | D46R | -2.26 | 1.17 | Positive charge (ii) |
| 3 | HcDoc | D48R | -2.25 | 0.73 | Positive charge (ii) |
| 4 | HcCoh | T128R | -2.20 | 0.70 | Stability hotspot (i) |
| 5 | HcDoc | D46K | -2.08 | 1.08 | Positive charge (ii) |
| 6 | HcDoc | D46L | -2.04 | 1.19 | $Ca^{2+}$-coordinating loop (iii) |
| 7 | HcCoh | A83P | -1.72 | 0.53 | Stability hotspot (i) |
| 8 | **HcDoc** | **D48N** | **-1.69** | **0.81** | **-** |
| 9 | HcDoc | D46M | -1.67 | 0.98 | $Ca^{2+}$-coordinating loop (iii) |
| 10 | HcDoc | D46Q | -1.51 | 1.02 | $Ca^{2+}$-coordinating loop (iii) |
| 11 | **HcCoh** | **S37V** | **-1.50** | **1.21** | **-** |
| 12 | **HcCoh** | **S37I** | **-1.37** | **1.51** | **-** |
| 13 | HcDoc | D48K | -1.37 | 0.57 | Positive charge (ii) |
| 14 | HcDoc | D46A | -1.33 | 0.87 | $Ca^{2+}$-coordinating loop (iii) |
| 15 | HcCoh | T128K | -1.28 | 0.54 | Stability hotspot (i) |
| 16 | HcCoh | S37M | -1.23 | 1.35 | Methionine destabilization (iv) |
| 17 | HcDoc | D46C | -1.21 | 0.88 | $Ca^{2+}$-coordinating loop (iii) |
| 18 | HcDoc | D48F | -1.15 | 0.58 | $Ca^{2+}$-coordinating loop (iii) |
| 19 | HcDoc | D46F | -1.12 | 1.32 | $Ca^{2+}$-coordinating loop (iii) |
| 20 | HcDoc | D46S | -1.09 | 0.99 | $Ca^{2+}$-coordinating loop (iii) |
| 21 | **HcCoh** | **N39L** | **-1.07** | **1.49** | **-** |
| 22 | HcDoc | D46Y | -1.07 | 1.35 | $Ca^{2+}$-coordinating loop (iii) |
| 23 | HcDoc | D48Y | -1.04 | 0.57 | $Ca^{2+}$-coordinating loop (iii) |
| 24 | HcCoh | T128L | -1.00 | 0.64 | Stability hotspot (i) |
| 25 | HcDoc | D46I | -0.97 | 1.08 | ΔΔG **>** -1 kcal/mol |

HcCoh – cohesin; HcDoc – dockerin

**Table S6: Per-residue MMGBSA energy decomposition for residues with |ΔG| > 1 kcal/mol.** Residues are classified as stabilizing (ΔG_Total < −1 kcal/mol) or destabilizing (ΔG_Total > 1 kcal/mol). Energy components include van der Waals (vdW), electrostatic (Elec), polar solvation (PolSolv), and non-polar solvation (NPSolv) contributions. Values represent the mean ± standard deviation (std) calculated over the 2,500 frame conformational ensemble. All energies are given in kcal/mol.

| Contribution | Protein | Residue | ΔG_vdW mean±std | ΔG_Elec mean±std | ΔG_PolSolv mean±std | ΔG_NPSolv mean±std | ΔG_Total mean±std |
|---|---|---|---|---|---|---|---|
| Stabilizing | HcDoc | $Ca^{2+}$ 222 | -0.03±0.01 | -114.22±6.52 | 107.20±6.13 | -0.00±0.00 | **-7.05±1.41** |
| Stabilizing | HcDoc | LEU 28 | -6.41±0.75 | -3.81±1.81 | 4.60±0.98 | -0.93±0.05 | **-6.55±1.23** |
| Stabilizing | HcDoc | ILE 53 | -5.78±0.55 | -0.92±0.59 | 2.23±0.50 | -0.76±0.05 | **-5.23±0.66** |
| Stabilizing | HcCoh | TYR 124 | -3.74±0.54 | -5.53±2.08 | 5.42±1.63 | -0.50±0.04 | **-4.36±1.03** |
| Stabilizing | HcCoh | MET 80 | -4.60±0.65 | -1.47±1.79 | 2.86±1.47 | -0.86±0.08 | **-4.06±0.68** |
| Stabilizing | HcCoh | PHE 67 | -4.07±0.46 | -0.39±0.75 | 0.98±0.66 | -0.55±0.05 | **-4.03±0.49** |
| Stabilizing | HcDoc | VAL 57 | -3.27±0.64 | -1.14±0.41 | 1.26±0.40 | -0.46±0.03 | **-3.61±0.69** |
| Stabilizing | HcDoc | ARG 60 | -2.91±0.81 | -91.11±4.84 | 91.02±4.73 | -0.60±0.05 | **-3.60±1.25** |
| Stabilizing | HcCoh | TYR 34 | -1.50±0.77 | -10.51±3.11 | 9.10±2.04 | -0.32±0.06 | **-3.23±1.35** |
| Stabilizing | HcDoc | ASN 54 | -3.20±0.65 | -3.85±1.06 | 4.31±0.80 | -0.42±0.03 | **-3.15±0.74** |
| Stabilizing | HcCoh | ALA 83 | -2.23±0.66 | -1.92±2.97 | 2.19±2.35 | -0.57±0.08 | **-2.52±0.84** |
| Stabilizing | HcCoh | GLY 127 | -1.65±0.62 | -2.54±0.97 | 2.19±0.71 | -0.41±0.06 | **-2.41±0.61** |
| Stabilizing | HcCoh | ARG 120 | -3.10±1.55 | -62.93±13.79 | 64.29±13.31 | -0.63±0.32 | **-2.36±2.31** |
| Stabilizing | HcCoh | ALA 81 | 0.21±0.69 | -7.62±1.22 | 5.18±0.76 | -0.06±0.02 | **-2.29±0.64** |
| Stabilizing | HcDoc | ARG 24 | -2.68±0.65 | -85.55±7.25 | 86.60±7.76 | -0.43±0.13 | **-2.06±0.62** |
| Stabilizing | HcDoc | LEU 58 | -1.57±0.25 | -0.58±0.28 | 0.52±0.27 | -0.18±0.02 | **-1.81±0.28** |
| Stabilizing | HcDoc | LYS 30 | -2.48±0.38 | -61.85±4.47 | 62.90±4.68 | -0.36±0.04 | **-1.78±0.66** |
| Stabilizing | HcDoc | ARG 52 | -2.84±0.92 | -82.23±10.30 | 84.09±9.73 | -0.63±0.11 | **-1.61±1.00** |
| Stabilizing | HcCoh | THR 128 | -2.06±0.35 | 0.39±0.91 | 0.51±0.89 | -0.39±0.06 | **-1.56±0.32** |
| Stabilizing | HcCoh | ALA 35 | -1.54±0.34 | 0.75±0.22 | -0.57±0.29 | -0.19±0.03 | **-1.55±0.37** |
| Stabilizing | HcDoc | GLY 29 | -1.91±0.39 | -0.05±0.98 | 0.95±1.01 | -0.30±0.06 | **-1.31±0.42** |
| Stabilizing | HcDoc | $CA^{2+}$ 223 | -0.00±0.00 | -91.32±2.74 | 90.09±2.71 | 0.00±0.00 | **-1.23±0.16** |

HcCoh – cohesin; HcDoc – dockerin

| Contribution | Protein | Residue | ΔG_vdW mean±std | ΔG_Elec mean±std | ΔG_PolSolv mean±std | ΔG_NPSolv mean±std | ΔG_Total mean±std |
|---|---|---|---|---|---|---|---|
| Stabilizing | HcDoc | VAL 27 | -2.36±0.70 | -3.59±1.88 | 5.11±1.45 | -0.36±0.07 | **-1.20±0.77** |
| Stabilizing | HcCoh | SER 76 | -1.24±0.56 | -3.60±2.23 | 3.84±1.52 | -0.14±0.03 | **-1.15±1.01** |
| Stabilizing | HcCoh | ALA 65 | -1.49±0.27 | 0.87±0.61 | -0.17±0.57 | -0.31±0.04 | **-1.09±0.35** |
| Stabilizing | HcCoh | LEU 36 | -1.19±0.24 | -0.64±0.27 | 0.85±0.38 | -0.04±0.01 | **-1.03±0.38** |
| Stabilizing | HcCoh | PHE 74 | -0.85±0.26 | -0.88±0.42 | 0.84±0.38 | -0.10±0.04 | **-1.00±0.33** |
| Destabilizing | HcDoc | ASP 61 | -0.59±0.70 | 57.86±5.36 | -54.13±5.47 | -0.13±0.05 | **3.01±1.24** |
| Destabilizing | HcDoc | ASP 55 | -0.42±0.04 | 58.41±2.88 | -56.33±2.82 | 0.00±0.00 | **1.66±0.35** |
| Destabilizing | HcDoc | ASP 46 | -0.73±0.18 | 56.38±3.38 | -54.51±3.31 | -0.08±0.03 | **1.05±0.35** |

HcCoh – cohesin; HcDoc – dockerin

**Table S7: Results of FoldX *BuildModel* calculations on the 2,500-frame conformational ensemble from the wild-type MD simulation, representing differences (Mut − WT) in the Gibbs energy of protein folding decomposed into individual energy contributions.** Included energy terms: Sidechain Hydrogen Bond (SHbond), Electrostatic (Elec), Solvation Hydrophobic (NPSolv), Solvation Polar (PolSolv), Van der Waals clashes (vdW clashes), Van der Waals (vdW), Partial Covalent Bonds (PCbond), and Total Energy. Values represent the mean ± standard deviation calculated over the 2,500-frame conformational ensemble (kcal/mol).

| Variant | ΔG SHbond mean±std | ΔG Elec mean±std | ΔG NPSolv mean±std | ΔG PolSolv mean±std | ΔG vdW clashes mean±std | ΔG vdW mean±std | ΔG PCbond mean±std | ΔG Total mean±std |
|---|---|---|---|---|---|---|---|---|
| D46N (HcDoc) | 0.58±1.12 | -2.00±0.85 | -0.23±0.27 | 0.11±1.54 | 0.98±1.00 | -0.36±0.40 | -2.15±1.26 | **-2.56±1.19** |
| D48N (HcDoc) | -0.42±0.87 | -0.79±0.51 | -0.48±0.27 | 1.84±1.28 | 0.62±0.66 | -0.70±0.39 | -2.51±1.50 | **-1.93±0.80** |
| N39L (HcCoh) | 1.60±0.97 | -0.00±0.18 | -2.18±0.29 | -0.71±0.44 | 0.60±0.94 | -0.67±0.25 | -0.00±0.00 | **-1.16±1.42** |
| S37I (HcCoh) | 0.55±0.39 | -0.00±0.10 | -3.70±0.21 | 0.69±0.32 | 1.57±1.30 | -1.71±0.15 | -0.00±0.00 | **-1.31±1.55** |
| S37V (HcCoh) | 0.56±0.35 | -0.00±0.05 | -2.54±0.21 | 0.26±0.24 | 0.99±0.96 | -1.08±0.11 | -0.00±0.00 | **-1.44±1.19** |
| N39L (HcCoh) / D46N (HcDoc) | 2.17±1.47 | -1.99±0.90 | -2.39±0.42 | -0.66±1.72 | 1.51±1.33 | -0.99±0.52 | -2.21±1.26 | **-3.84±1.85** |
| N39L (HcCoh) / D48N (HcDoc) | 1.20±1.35 | -0.74±0.65 | -2.63±0.42 | 1.06±1.49 | 1.17±1.11 | -1.33±0.51 | -2.52±1.55 | **-3.11±1.60** |
| S37I (HcCoh) / D46N (HcDoc) | 1.08±1.19 | -1.93±0.88 | -3.93±0.37 | 0.73±1.67 | 2.27±1.59 | -2.04±0.47 | -2.15±1.32 | **-4.10±1.90** |
| S37I (HcCoh) / D48N (HcDoc) | 0.16±0.95 | -0.74±0.59 | -4.16±0.36 | 2.43±1.41 | 2.03±1.41 | -2.37±0.45 | -2.46±1.57 | **-3.30±1.68** |
| S37I (HcCoh) / N39L (HcCoh) | 2.14±1.05 | -0.01±0.27 | -5.85±0.36 | -0.12±0.58 | 2.08±1.62 | -2.32±0.30 | 0.00±0.00 | **-2.61±2.13** |
| S37V (HcCoh) / D46N (HcDoc) | 1.10±1.20 | -1.96±0.86 | -2.78±0.37 | 0.34±1.63 | 1.79±1.37 | -1.42±0.44 | -2.13±1.35 | **-4.13±1.68** |
| S37V (HcCoh) / D48N (HcDoc) | 0.15±0.93 | -0.75±0.58 | -2.99±0.35 | 2.01±1.40 | 1.53±1.16 | -1.74±0.44 | -2.45±1.55 | **-3.35±1.42** |
| S37V (HcCoh) / N39L (HcCoh) | 2.15±1.02 | -0.00±0.22 | -4.71±0.38 | -0.51±0.53 | 1.56±1.38 | -1.72±0.28 | -0.00±0.00 | **-2.68±1.89** |

HcCoh – cohesin; HcDoc – dockerin

| Variant | ΔG SHbond mean±std | ΔG Elec mean±std | ΔG NPSolv mean±std | ΔG PolSolv mean±std | ΔG vdW clashes mean±std | ΔG vdW mean±std | ΔG PCbond mean±std | ΔG Total mean±std |
|---|---|---|---|---|---|---|---|---|
| S37I (HcCoh) / N39L (HcCoh) / D46N (HcDoc) | 2.72±1.57 | -1.91±0.95 | -6.10±0.49 | -0.05±1.86 | 2.87±1.91 | -2.66±0.57 | -2.06±1.47 | **-5.18±2.46** |
| S37I (HcCoh) / N39L (HcCoh) / D48N (HcDoc) | 1.77±1.39 | -0.71±0.70 | -6.29±0.47 | 1.55±1.56 | 2.60±1.74 | -2.95±0.55 | -2.43±1.60 | **-4.49±2.25** |
| S37V (HcCoh) / N39L (HcCoh) / D46N (HcDoc) | 2.69±1.55 | -1.96±0.91 | -4.97±0.49 | -0.40±1.77 | 2.39±1.70 | -2.07±0.53 | -2.08±1.43 | **-5.28±2.23** |
| S37V (HcCoh) / N39L (HcCoh) / D48N (HcDoc) | 1.80±1.39 | -0.72±0.67 | -5.15±0.49 | 1.19±1.55 | 2.15±1.54 | -2.35±0.53 | -2.52±1.59 | **-4.56±2.05** |

HcCoh – cohesin; HcDoc – dockerin

**Table S8: Results of FoldX *AnalyzeComplex* calculations on the 2,500-frame ensemble of mutant structures generated by FoldX *BuildModel***, representing the HcCoh–HcDoc interaction energy, intrachain clashes within each domain, and the folding stability of each domain. Values represent the mean ± standard deviation calculated over the 2,500-frame ensemble (kcal/mol).

| Variant | Interaction energy mean±std | Interclashes HcCoh mean±std | Interclashes HcDoc mean±std | Stability HcCoh mean±std | Stability HcDoc mean±std |
|---|---|---|---|---|---|
| D46N (HcDoc) | **-0.16±0.05** | 0.00±0.00 | 0.33±0.12 | -0.05±0.00 | -4.10±0.12 |
| D48N (HcDoc) | **-0.44±0.01** | 0.01±0.00 | 0.23±0.04 | -0.08±0.01 | -3.23±0.13 |
| N39L (HcCoh) | **-0.62±0.04** | 0.42±0.09 | 0.00±0.00 | -1.45±0.05 | 0.01±0.00 |
| S37I (HcCoh) | **-2.08±0.02** | 0.75±0.15 | 0.10±0.02 | -0.10±0.10 | -0.01±0.00 |
| S37V (HcCoh) | **-1.94±0.04** | 0.51±0.10 | 0.04±0.01 | -0.03±0.07 | -0.02±0.00 |
| N39L (HcCoh) / D46N (HcDoc) | **-0.83±0.01** | 0.41±0.10 | 0.31±0.11 | -1.53±0.05 | -4.21±-0.12 |
| N39L (HcCoh) / D48N (HcDoc) | **-1.21±0.00** | 0.39±0.08 | 0.24±0.05 | -1.59±0.05 | -3.27±0.14 |
| S37I (HcCoh) / D46N (HcDoc) | **-2.34±0.06** | 0.73±0.13 | 0.31±0.11 | -0.16±0.09 | -4.36±0.11 |
| S37I (HcCoh) / D48N (HcDoc) | **-2.69±0.08** | 0.75±0.13 | 0.30±0.08 | -0.19±0.09 | -3.30±0.11 |
| S37I (HcCoh) / N39L (HcCoh) | **-2.64±0.03** | 1.21±0.27 | 0.12±0.02 | -1.61±0.13 | -0.02±0.00 |
| S37V (HcCoh) / D46N (HcDoc) | **-2.18±0.08** | 0.51±0.09 | 0.26±0.10 | -0.09±0.08 | -4.35±0.13 |
| S37V (HcCoh) / D48N (HcDoc) | **-2.49±0.06** | 0.52±0.08 | 0.24±0.04 | -0.15±0.07 | -3.31±0.12 |
| S37V (HcCoh) / N39L (HcCoh) | **-2.60±0.05** | 0.95±0.19 | 0.05±0.01 | -1.54±0.10 | -0.02±0.00 |
| S37I (HcCoh) / N39L (HcCoh) / D46N (HcDoc) | **-2.85±0.05** | 1.21±0.23 | 0.30±0.12 | -1.67±0.15 | -4.26±0.11 |
| S37I (HcCoh) / N39L (HcCoh) / D48N (HcDoc) | **-3.18±0.02** | 1.20±0.24 | 0.30±0.08 | -1.74±0.14 | -3.30±0.10 |
| S37V (HcCoh) / N39L (HcCoh) / D46N (HcDoc) | **-2.79±0.01** | 0.93±0.19 | 0.26±0.10 | -1.63±0.11 | -4.30±0.14 |
| S37V (HcCoh) / N39L (HcCoh) / D48N (HcDoc) | **-3.14±0.03** | 0.94±0.19 | 0.26±0.05 | -1.68±0.11 | -3.33±0.12 |

HcCoh – cohesin; HcDoc – dockerin.

**Table S9: Overall MMPBSA binding free energy ($\Delta G_{binding}$) calculated using internal dielectric constants of 5 and 10.** Values represent the mean ± standard deviation over the 2,500-frame conformational ensemble (kcal/mol). HcCoh mutants show small or no improvement in binding affinity relative to the WT, while HcDoc mutants exhibit a 6–17 kcal/mol decrease in $\Delta G_{binding}$, indicating substantially enhanced complex stability.

| Variant | $\Delta G_{binding}$, indi = 5 mean±std [kcal/mol] | $\Delta G_{binding}$, indi = 10 mean±std [kcal/mol] |
|---|---|---|
| HcCoh-WT – HcDoc-WT | -40.6±0.2 | -36.3±0.2 |
| HcCoh-WT – HcDoc-D46N | -54.6±0.2 | -41.9±0.1 |
| HcCoh-WT – HcDoc-D48N | -57.2±0.2 | -44.0±0.2 |
| HcCoh-S37I+N39L – HcDoc-WT | -39.0±0.2 | -35.5±0.2 |
| HcCoh-S37V+N39L – HcDoc-WT | -41.8±0.3 | -36.8±0.2 |

HcCoh – cohesin; HcDoc – dockerin.

**Table S10: Pairwise MMPBSA energy decomposition calculated using an internal dielectric constant of 5, for residue pairs with |ΔΔG_total| > 2 kcal/mol in the D46N mutant.** Interactions are classified as favorable (ΔΔG_total < 0 kcal/mol) or unfavorable (ΔΔG_total > 0 kcal/mol). Energy components include van der Waals (vdW), electrostatic (Elec), and polar solvation (PolSolv) contributions. ΔG_total WT and ΔG_total Mut columns report the original pairwise interaction energies for the wild-type and mutant systems, respectively. ΔΔG_total = ΔG_Total(Mut) − ΔG_Total(WT), reflecting the change in pairwise interaction energy upon mutation. Values represent mean ± standard deviation calculated over the 2,500-frame conformational ensemble. All energies are given in kcal/mol.

| Mutated residue(s) | Interacting residue | ΔΔG_vdW mean±std | ΔΔG_Elec mean±std | ΔΔG_PolSolv mean±std | ΔΔG_total mean±std | ΔG_Total WT mean±std | ΔG_Total Mut mean±std |
|---|---|---|---|---|---|---|---|
| D46N (HcDoc) | GLU130 (HcCoh) | -0.00±0.01 | -17.46±1.33 | 3.14±0.22 | **-14.32±1.11** | 13.71±1.06 | -0.61±0.32 |
| D46N (HcDoc) | GLU129 (HcCoh) | -0.00±0.02 | -15.48±1.67 | 2.80±0.29 | **-12.68±1.38** | 11.85±1.33 | -0.83±0.36 |
| D46N (HcDoc) | ASP63 (HcCoh) | 0.00±0.00 | -7.89±0.18 | 1.48±0.03 | **-6.41±0.15** | 6.32±0.13 | -0.09±0.07 |
| D46N (HcDoc) | ASP84 (HcCoh) | 0.00±0.00 | -7.32±0.19 | 1.38±0.04 | **-5.94±0.16** | 5.81±0.14 | -0.12±0.06 |
| D46N (HcDoc) | ASP90 (HcCoh) | 0.00±0.00 | -6.83±0.16 | 1.29±0.03 | **-5.54±0.13** | 5.41±0.12 | -0.12±0.05 |
| D46N (HcDoc) | ASP62 (HcCoh) | 0.00±0.00 | -6.54±0.32 | 1.24±0.06 | **-5.30±0.26** | 5.26±0.26 | -0.04±0.05 |
| D46N (HcDoc) | ASP60 (HcCoh) | 0.00±0.00 | -6.30±0.16 | 1.19±0.03 | **-5.11±0.13** | 5.05±0.12 | -0.06±0.05 |
| D46N (HcDoc) | ASP92 (HcCoh) | 0.00±0.00 | -6.12±0.15 | 1.16±0.03 | **-4.96±0.12** | 4.87±0.11 | -0.09±0.04 |
| D46N (HcDoc) | ASP138 (HcCoh) | 0.00±0.00 | -6.08±0.31 | 1.14±0.06 | **-4.93±0.25** | 4.91±0.25 | -0.03±0.03 |
| D46N (HcDoc) | ASP7 (HcCoh) | 0.00±0.00 | -5.81±0.18 | 1.10±0.03 | **-4.71±0.15** | 4.66±0.14 | -0.05±0.03 |
| D46N (HcDoc) | GLU59 (HcCoh) | 0.00±0.00 | -5.61±0.19 | 1.06±0.04 | **-4.54±0.15** | 4.48±0.15 | -0.07±0.04 |
| D46N (HcDoc) | GLU53 (HcCoh) | 0.00±0.00 | -5.22±0.16 | 0.99±0.03 | **-4.23±0.13** | 4.21±0.13 | -0.02±0.03 |
| D46N (HcDoc) | ASP113 (HcCoh) | 0.00±0.00 | -5.11±0.18 | 0.96±0.03 | **-4.15±0.15** | 4.15±0.15 | -0.00±0.02 |
| D46N (HcDoc) | GLU71 (HcCoh) | -0.00±0.00 | -4.91±0.19 | 0.93±0.04 | **-3.99±0.15** | 4.01±0.15 | 0.02±0.02 |
| D46N (HcDoc) | ASP43 (HcCoh) | 0.00±0.00 | -4.80±0.14 | 0.90±0.03 | **-3.89±0.12** | 3.90±0.11 | 0.01±0.02 |
| D46N (HcDoc) | GLU110 (HcCoh) | 0.00±0.00 | -3.90±0.11 | 0.73±0.02 | **-3.17±0.09** | 3.17±0.09 | -0.00±0.01 |

HcCoh – cohesin; HcDoc – dockerin.

| Mutated residue(s) | Interacting residue | ΔΔG_vdW mean±std | ΔΔG_Elec mean±std | ΔΔG_PolSolv mean±std | ΔΔG_total mean±std | ΔG_Total WT mean±std | ΔG_Total Mut mean±std |
|---|---|---|---|---|---|---|---|
| D46N (HcDoc) | GLU104 (HcCoh) | 0.00±0.00 | -3.61±0.15 | 0.68±0.03 | **-2.93±0.12** | 2.93±0.12 | 0.00±0.01 |
| D46N (HcDoc) | GLU14 (HcCoh) | 0.00±0.00 | -3.58±0.10 | 0.68±0.02 | **-2.91±0.08** | 2.90±0.08 | -0.01±0.01 |
| D46N (HcDoc) | ASN146 (HcCoh) | 0.00±0.00 | -3.42±0.19 | 0.64±0.04 | **-2.78±0.15** | 2.77±0.15 | -0.01±0.01 |
| D46N (HcDoc) | GLY127 (HcCoh) | 0.03±0.08 | -2.38±0.37 | 0.21±0.05 | **-2.14±0.35** | 1.88±0.24 | -0.26±0.26 |
| D46N (HcDoc) | LYS101 (HcCoh) | 0.00±0.00 | 3.99±0.12 | -0.75±0.02 | **3.24±0.10** | -3.24±0.10 | 0.00±0.01 |
| D46N (HcDoc) | LYS46 (HcCoh) | 0.00±0.00 | 4.33±0.16 | -0.81±0.03 | **3.52±0.13** | -3.52±0.13 | -0.01±0.02 |
| D46N (HcDoc) | LYS99 (HcCoh) | 0.00±0.00 | 4.58±0.14 | -0.87±0.03 | **3.71±0.11** | -3.70±0.11 | 0.02±0.02 |
| D46N (HcDoc) | ARG10 (HcCoh) | 0.00±0.00 | 4.86±0.29 | -0.92±0.06 | **3.95±0.24** | -3.92±0.24 | 0.02±0.02 |
| D46N (HcDoc) | LYS91 (HcCoh) | 0.00±0.00 | 5.71±0.26 | -1.08±0.05 | **4.62±0.21** | -4.54±0.21 | 0.08±0.03 |
| D46N (HcDoc) | ALA1 (HcCoh) | 0.00±0.00 | 6.21±0.55 | -1.17±0.10 | **5.04±0.45** | -4.94±0.45 | 0.10±0.04 |
| D46N (HcDoc) | ARG87 (HcCoh) | 0.00±0.00 | 6.79±0.16 | -1.29±0.03 | **5.50±0.13** | -5.42±0.12 | 0.08±0.05 |
| D46N (HcDoc) | ARG120 (HcCoh) | 0.00±0.00 | 9.89±0.94 | -1.84±0.17 | **8.06±0.77** | -8.16±0.77 | -0.10±0.07 |
| D46N (HcDoc) | ARG85 (HcCoh) | -0.00±0.00 | 10.15±0.67 | -1.89±0.12 | **8.27±0.55** | -7.97±0.54 | 0.29±0.12 |
| D46N (HcDoc) | LYS132 (HcCoh) | 0.00±0.00 | 14.23±1.89 | -2.58±0.32 | **11.65±1.57** | -11.27±1.56 | 0.39±0.21 |

HcCoh – cohesin; HcDoc – dockerin.

**Table S11: Pairwise MMPBSA energy decomposition calculated using an internal dielectric constant of 5, for residue pairs with |ΔΔG_total| > 2 kcal/mol in the D48N mutant.** Interactions are classified as favorable (ΔΔG_total < 0 kcal/mol) or unfavorable (ΔΔG_total > 0 kcal/mol). Energy components include van der Waals (vdW), electrostatic (Elec), and polar solvation (PolSolv) contributions. ΔG_total WT and ΔG_total Mut columns report the original pairwise interaction energies for the wild-type and mutant systems, respectively. ΔΔG_total = ΔG_Total(Mut) − ΔG_Total(WT), reflecting the change in pairwise interaction energy upon mutation. Values represent mean ± standard deviation calculated over the 2,500-frame conformational ensemble. All energies are given in kcal/mol.

| Mutated residue(s) | Interacting residue | ΔΔG vdW mean±std | ΔΔG Elec mean±std | ΔΔG PolSolv mean±std | ΔΔG Total mean±std | ΔG Total WT mean±std | ΔG Total Mut mean±std |
|---|---|---|---|---|---|---|---|
| D48N (HcDoc) | GLU130 (HcCoh) | 0.00±0.05 | -23.25±2.43 | 4.15±0.41 | **-19.09±2.00** | 17.57±1.90 | -1.52±0.61 |
| D48N (HcDoc) | GLU129 (HcCoh) | 0.00±0.01 | -13.91±1.20 | 2.55±0.21 | **-11.36±0.99** | 10.71±0.97 | -0.65±0.23 |
| D48N (HcDoc) | ASP63 (HcCoh) | 0.00±0.00 | -7.30±0.17 | 1.38±0.03 | **-5.92±0.14** | 5.88±0.13 | -0.04±0.05 |
| D48N (HcDoc) | ASP138 (HcCoh) | 0.00±0.00 | -6.73±0.39 | 1.26±0.07 | **-5.47±0.32** | 5.44±0.31 | -0.03±0.04 |
| D48N (HcDoc) | ASP90 (HcCoh) | 0.00±0.00 | -6.55±0.15 | 1.24±0.03 | **-5.31±0.12** | 5.21±0.12 | -0.10±0.04 |
| D48N (HcDoc) | ASP84 (HcCoh) | 0.00±0.00 | -6.54±0.16 | 1.24±0.03 | **-5.30±0.13** | 5.23±0.12 | -0.07±0.04 |
| D48N (HcDoc) | ASP92 (HcCoh) | 0.00±0.00 | -6.20±0.16 | 1.17±0.03 | **-5.03±0.13** | 4.93±0.12 | -0.09±0.03 |
| D48N (HcDoc) | ASP7 (HcCoh) | 0.00±0.00 | -6.16±0.21 | 1.16±0.04 | **-5.00±0.17** | 4.94±0.17 | -0.05±0.03 |
| D48N (HcDoc) | ASP62 (HcCoh) | 0.00±0.00 | -6.12±0.27 | 1.16±0.05 | **-4.96±0.22** | 4.96±0.21 | -0.00±0.03 |
| D48N (HcDoc) | ASP60 (HcCoh) | 0.00±0.00 | -5.93±0.14 | 1.12±0.03 | **-4.81±0.12** | 4.78±0.11 | -0.02±0.03 |
| D48N (HcDoc) | ASP113 (HcCoh) | 0.00±0.00 | -5.58±0.23 | 1.05±0.04 | **-4.53±0.19** | 4.54±0.18 | 0.01±0.03 |
| D48N (HcDoc) | GLU59 (HcCoh) | 0.00±0.00 | -5.37±0.17 | 1.02±0.03 | **-4.35±0.14** | 4.31±0.13 | -0.04±0.03 |
| D48N (HcDoc) | GLU53 (HcCoh) | 0.00±0.00 | -5.24±0.16 | 0.99±0.03 | **-4.24±0.13** | 4.24±0.13 | -0.01±0.02 |
| D48N (HcDoc) | ASP43 (HcCoh) | 0.00±0.00 | -5.16±0.18 | 0.97±0.03 | **-4.19±0.15** | 4.22±0.15 | 0.03±0.03 |
| D48N (HcDoc) | GLU71 (HcCoh) | 0.00±0.00 | -5.16±0.21 | 0.97±0.04 | **-4.19±0.17** | 4.24±0.17 | 0.05±0.02 |
| D48N (HcDoc) | GLU110 (HcCoh) | 0.00±0.00 | -4.16±0.13 | 0.78±0.03 | **-3.38±0.11** | 3.39±0.11 | 0.01±0.02 |

HcCoh – cohesin; HcDoc – dockerin

| Mutated residue(s) | Interacting residue | ΔΔG vdW mean±std | ΔΔG Elec mean±std | ΔΔG PolSolv mean±std | ΔΔG Total mean±std | ΔG Total WT mean±std | ΔG Total Mut mean±std |
|---|---|---|---|---|---|---|---|
| D48N (HcDoc) | GLU104 (HcCoh) | 0.00±0.00 | -3.75±0.16 | 0.71±0.03 | **-3.04±0.13** | 3.05±0.13 | 0.01±0.01 |
| D48N (HcDoc) | GLU14 (HcCoh) | 0.00±0.00 | -3.71±0.10 | 0.70±0.02 | **-3.01±0.08** | 3.01±0.08 | 0.00±0.01 |
| D48N (HcDoc) | ASN146 (HcCoh) | 0.00±0.00 | -3.57±0.20 | 0.67±0.04 | **-2.89±0.17** | 2.89±0.16 | -0.01±0.01 |
| D48N (HcDoc) | LYS101 (HcCoh) | 0.00±0.00 | 4.12±0.13 | -0.78±0.02 | **3.34±0.11** | -3.35±0.11 | -0.01±0.01 |
| D48N (HcDoc) | LYS46 (HcCoh) | 0.00±0.00 | 4.63±0.20 | -0.87±0.04 | **3.76±0.16** | -3.79±0.16 | -0.02±0.02 |
| D48N (HcDoc) | LYS99 (HcCoh) | 0.00±0.00 | 4.65±0.14 | -0.88±0.03 | **3.77±0.11** | -3.77±0.11 | 0.00±0.02 |
| D48N (HcDoc) | ARG10 (HcCoh) | 0.00±0.00 | 5.22±0.34 | -0.98±0.06 | **4.24±0.28** | -4.22±0.28 | 0.02±0.02 |
| D48N (HcDoc) | LYS91 (HcCoh) | 0.00±0.00 | 5.58±0.24 | -1.06±0.05 | **4.52±0.19** | -4.45±0.19 | 0.07±0.03 |
| D48N (HcDoc) | ARG87 (HcCoh) | 0.00±0.00 | 6.27±0.14 | -1.19±0.03 | **5.08±0.11** | -5.04±0.11 | 0.04±0.04 |
| D48N (HcDoc) | ALA1 (HcCoh) | 0.00±0.00 | 6.41±0.57 | -1.20±0.10 | **5.20±0.46** | -5.08±0.46 | 0.12±0.04 |
| D48N (HcDoc) | ARG85 (HcCoh) | -0.00±0.00 | 8.95±0.55 | -1.68±0.10 | **7.27±0.45** | -7.10±0.44 | 0.17±0.09 |
| D48N (HcDoc) | ARG120 (HcCoh) | 0.00±0.00 | 11.18±1.23 | -2.07±0.22 | **9.11±1.01** | -9.31±1.00 | -0.20±0.13 |
| D48N (HcDoc) | LYS132 (HcCoh) | 0.00±0.01 | 17.32±2.86 | -3.14±0.50 | **14.18±2.37** | -13.31±2.34 | 0.87±0.37 |

HcCoh – cohesin; HcDoc – dockerin

**Table S12: Pairwise MMPBSA energy decomposition calculated using an internal dielectric constant of 5, for residue pairs with |ΔΔG_total| > 0.25 kcal/mol in the S37I/N39L mutant.** Interactions are classified as favorable (ΔΔG_total < 0 kcal/mol) or unfavorable (ΔΔG_total > 0 kcal/mol). Energy components include van der Waals (vdW), electrostatic (Elec), and polar solvation (PolSolv) contributions. For the S37I/N39L double mutant, the reported ΔΔG values represent the sum of pairwise interaction energy changes for both mutated residues (positions 37 and 39) with each interacting partner, reflecting the combined effect of both substitutions. ΔG Total WT and ΔG Total Mut columns report the original pairwise interaction energies for the wild-type and mutant systems, respectively. ΔΔG_Total = ΔG_Total(Mut) − ΔG_Total(WT), reflecting the change in pairwise interaction energy upon mutation. Values represent mean ± standard deviation calculated over the 2,500-frame conformational ensemble. All energies are given in kcal/mol.

| Mutated residue(s) | Interacting residue | ΔΔG vdW mean±std | ΔΔG Elec mean±std | ΔΔG PolSolv mean±std | ΔΔG Total mean±std | ΔG Total WT mean±std | ΔG Total Mut mean±std |
|---|---|---|---|---|---|---|---|
| S37I (HcCoh) + N39L (HcCoh) | ILE37 (HcCoh) | 0.00±0.00 | 0.00±0.00 | -0.77±0.30 | **-0.77±0.30** | 0.84±0.30 | 0.07±0.01 |
| S37I (HcCoh) + N39L (HcCoh) | LYS30 (HcDoc) | -0.01±0.01 | -0.37±0.59 | 0.06±0.09 | **-0.33±0.50** | 0.98±0.49 | 0.65±0.07 |
| S37I (HcCoh) + N39L (HcCoh) | LEU28 (HcDoc) | -0.58±0.32 | 0.29±0.54 | -0.00±0.03 | **-0.30±0.62** | -0.94±0.55 | -1.24±0.28 |
| S37I (HcCoh) + N39L (HcCoh) | ARG24 (HcDoc) | -0.06±0.05 | -0.26±0.49 | 0.05±0.08 | **-0.27±0.42** | 0.55±0.41 | 0.28±0.07 |
| S37I (HcCoh) + N39L (HcCoh) | LEU39 (HcCoh) | 0.00±0.00 | 0.00±0.00 | -0.25±0.22 | **-0.25±0.22** | 0.26±0.22 | 0.01±0.00 |
| S37I (HcCoh) + N39L (HcCoh) | CA 222 (HcDoc) | -0.00±0.00 | 0.33±0.87 | -0.05±0.13 | **0.28±0.74** | -0.50±0.72 | -0.22±0.16 |
| S37I (HcCoh) + N39L (HcCoh) | ARG52 (HcDoc) | -0.09±0.08 | 0.47±0.95 | -0.06±0.12 | **0.32±0.84** | -1.07±0.82 | -0.75±0.19 |
| S37I (HcCoh) + N39L (HcCoh) | ASP25 (HcDoc) | -0.02±0.01 | 0.42±0.70 | -0.05±0.10 | **0.35±0.61** | -1.13±0.60 | -0.78±0.08 |

HcCoh – cohesin; HcDoc – dockerin.

**Table S13: Pairwise MMPBSA energy decomposition calculated using an internal dielectric constant of 5, for residue pairs with |ΔΔG_total| > 0.25 kcal/mol in the S37V/N39L mutant.** Interactions are classified as favorable (ΔΔG_total < 0 kcal/mol) or unfavorable (ΔΔG_total > 0 kcal/mol). Energy components include van der Waals (vdW), electrostatic (Elec), and polar solvation (PolSolv) contributions. For the S37I/N39L double mutant, the reported ΔΔG values represent the sum of pairwise interaction energy changes for both mutated residues (positions 37 and 39) with each interacting partner, reflecting the combined effect of both substitutions. ΔG Total WT and ΔG Total Mut columns report the original pairwise interaction energies for the wild-type and mutant systems, respectively. ΔΔG Total = ΔG_Total(Mut) − ΔG_Total(WT), reflecting the change in pairwise interaction energy upon mutation. Values represent mean ± standard deviation calculated over the 2,500-frame conformational ensemble. All energies are given in kcal/mol.

| Mutated residue(s) | Interacting residue | ΔΔG vdW mean±std | ΔΔG Elec mean±std | ΔΔG PolSolv mean±std | ΔΔG Total mean±std | ΔG Total WT mean±std | ΔG Total Mut mean±std |
|---|---|---|---|---|---|---|---|
| S37V (HcCoh) + N39L (HcCoh) | VAL37 (HcCoh) | 0.00±0.00 | 0.00±0.00 | -0.76±0.30 | **-0.76±0.30** | 0.84±0.30 | 0.08±0.01 |
| S37V (HcCoh) + N39L (HcCoh) | LYS30 (HcDoc) | -0.01±0.01 | -0.33±0.59 | 0.05±0.09 | **-0.29±0.50** | 0.98±0.49 | 0.69±0.07 |
| S37V (HcCoh) + N39L (HcCoh) | LEU28 (HcDoc) | -0.54±0.31 | 0.27±0.55 | -0.01±0.03 | **-0.27±0.62** | -0.94±0.55 | -1.21±0.28 |
| S37V (HcCoh) + N39L (HcCoh) | LEU39 (HcCoh) | 0.00±0.00 | 0.00±0.00 | -0.25±0.22 | **-0.25±0.22** | 0.26±0.22 | 0.01±0.00 |
| S37V (HcCoh) + N39L (HcCoh) | ARG52 (HcDoc) | -0.04±0.05 | 0.38±0.95 | -0.05±0.12 | **0.29±0.84** | -1.07±0.82 | -0.78±0.18 |
| S37V (HcCoh) + N39L (HcCoh) | ASP25 (HcDoc) | -0.01±0.01 | 0.38±0.70 | -0.05±0.10 | **0.31±0.61** | -1.13±0.60 | -0.82±0.08 |

HcCoh – cohesin; HcDoc – dockerin.

### Supplementary figures

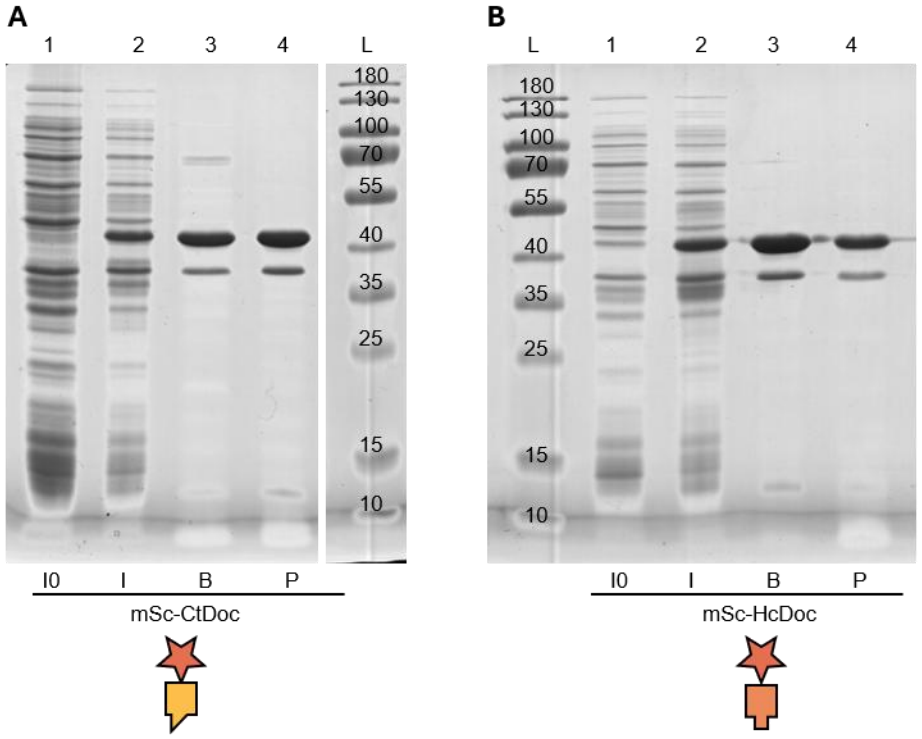


**Supplementary figure S1. Verification of expression and purification of dockerin-tagged reporter proteins by sodium dodecyl sulfate polyacrylamide gel electrophoresis.** (A) mScarlet-I-CtDoc. (B) mScarlet-I-HcDoc (lanes 1-4) and GFP-CtDoc (lanes 6-9). L = PageRuler Prestained Protein Ladder, 10 to 180 kDa. I0 is lysate of *E. coli* BL21-Gold (DE3) carrying pET21b+_*mScarlet-I-ctDoc-his*, pET21b+_*mScarlet-I-hcDoc-his,* or pET21b+_*gfp-ctDoc-his*, harvested 2.5 hours after starting the cultivation, before induction. I is lysate of *E. coli* BL21-Gold (DE3) carrying pET21b+_*mScarlet-I-ctDoc-his*, pET21b+_*mScarlet-I-hcDoc-his,* or pET21b+_*gfp-ctDoc-his*, harvested 20 hours after inducing the production of the protein of interest. B is sample taken from the purification column after washing it with purification buffer B. P is sample of purified mScarlet-I-CtDoc, mScarlet-I-HcDoc, or GFP-CtDoc collected after the imidazole dilution. Theoretical molecular weights of the target proteins were 36.5 kDa, 37.3 kDa, and 36.2 kDa for mScarlet-I-CtDoc, mScarlet-I-HcDoc, or GFP-CtDoc, respectively. The two major bands in B and P lines likely represent fully and partially denatured chimeric molecules of mScarlet-I linked to the thermophilic dockerin.

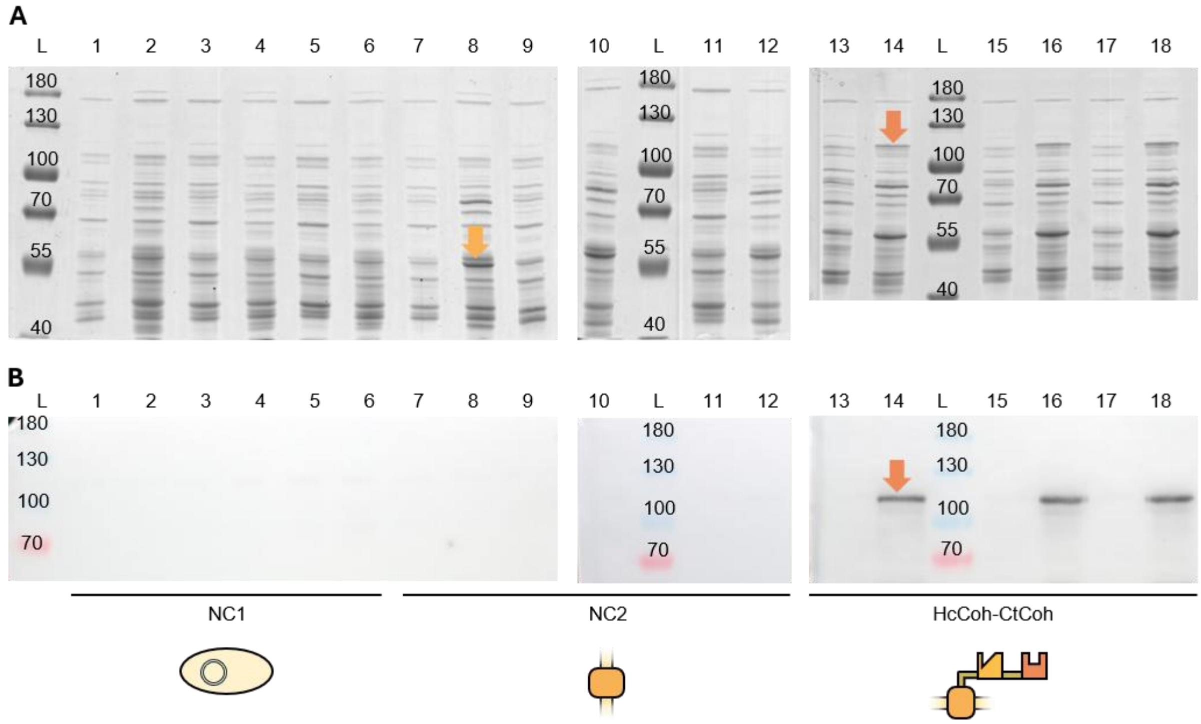


**Supplementary figure S2. Verification of expression of the HcCoh-CtCoh scaffoldin by sodium dodecyl sulfate polyacrylamide gel electrophoresis (A) and western blotting (B).** Lysates of *P. putida* EM371 (NC1; lanes 1-6), *P. putida* EM371 carrying pSEVA238_*ag43AT* (NC2; lanes 7-12), or *P. putida* EM371 carrying pSEVA238_*ag43-hcCoh-ctCoh-his* (HcCoh-CtCoh; lanes 13-18). Odd lanes contain samples collected 2.5 hours after the start of cultivation, before induction (I0 samples). Even lanes contain samples collected 4.5 hours after inducing the production of the protein of interest (I samples). L = PageRuler Prestained Protein Ladder, 10 to 180 kDa. The light orange arrow in lane 8 indicates the band representing the Ag43 autotransporter (theoretical molecular weight = 59.2 kDa). The dark orange arrow in lane 14 indicates the band representing the HcCoh-CtCoh scaffoldin (theoretical molecular weight = 93.5 kDa). The band representing the scaffoldin appears to have a higher molecular weight than the calculated value, which is likely caused by incomplete denaturation of the complex of thermophilic origin.[7] The correct identification of the band representing the HcCoh-CtCoh scaffoldin was confirmed by western blotting using an anti-His-tag antibody.

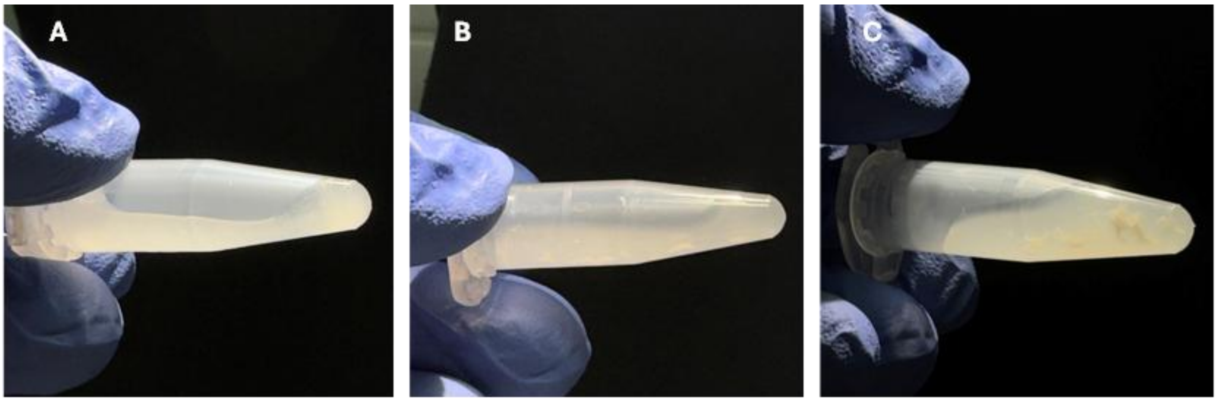


**Supplementary figure S3. Photographs of *Pseudomonas putida* EM371 cell suspensions incubated at 50°C for different time intervals.** Harvested cells were washed with TBS-Ca-T buffer, samples of $OD_{600}$ = 10.0 were prepared, and incubated at 50 °C for 0 (A), 40 (B), or 60 min (C). After the incubation, the cells were centrifuged at 5,000 g for 3 min, resuspended in TBS-Ca-T buffer, and photographed. Cell clumps were visible after 60 min of incubation.

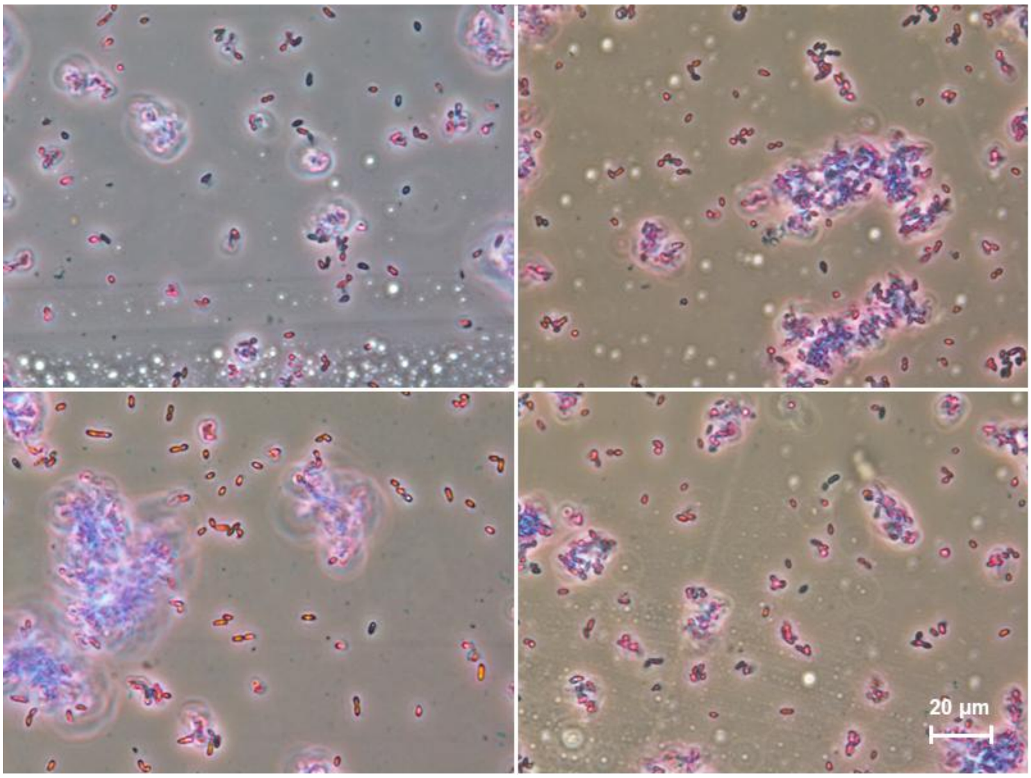


**Supplementary figure S4. Visual verification of the structural integrity of *Pseudomonas putida* EM371 after 40 min at 50 °C using light microscopy.** Four microphotographs showing structurally intact *P. putida* EM371 individual cells or small clusters after the high-temperature treatment. Scale (white line in the bottom right corner) is 20 µm. Details on sample preparation and microphotography equipment are available in the **Supplementary methods**.

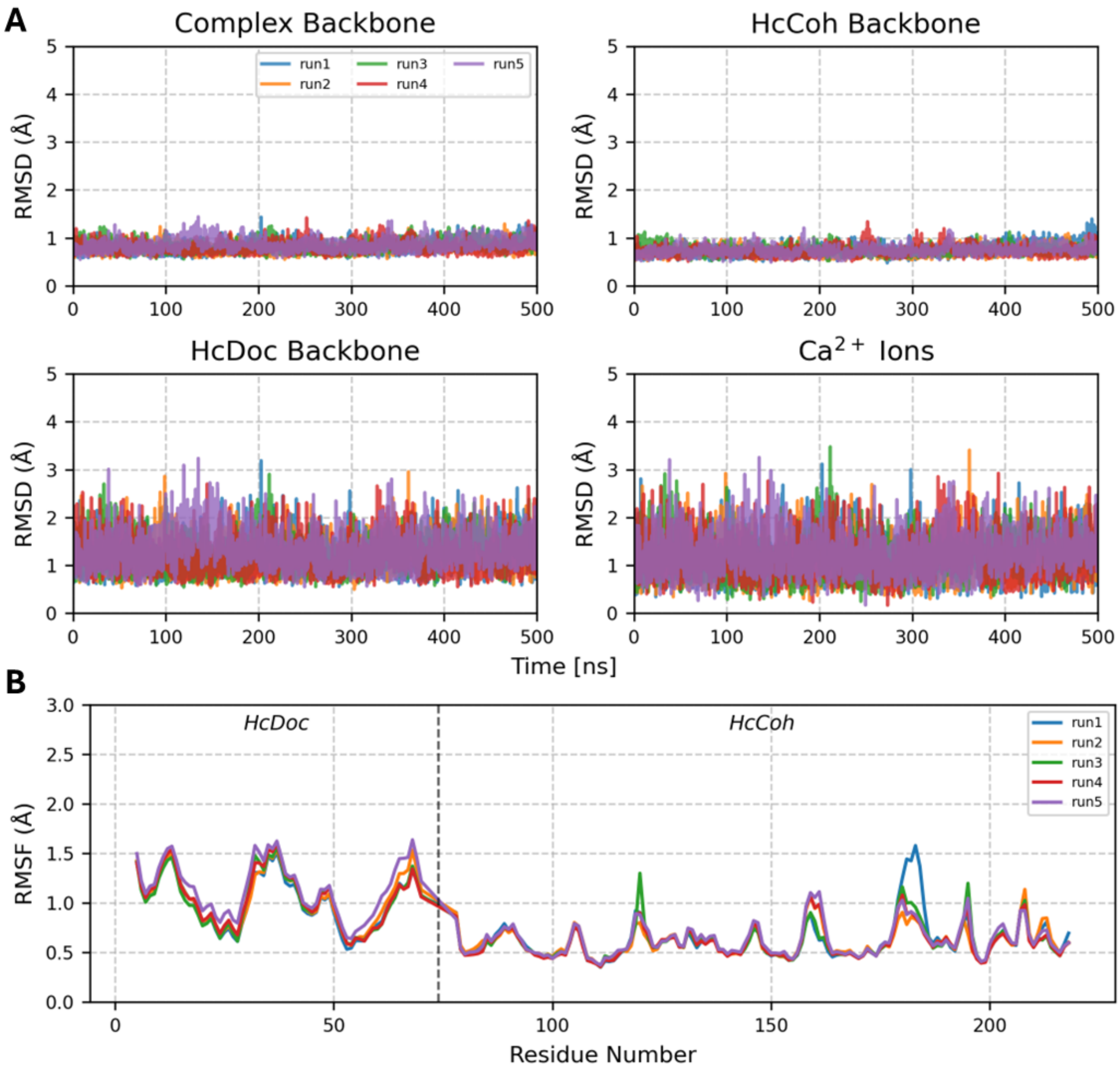


**Supplementary figure S5**. **MD simulations of the wild-type HcCoh-HcDoc complex demonstrate stable and convergent trajectories.** (A) Backbone RMSD over 500 ns calculated relative to the energy-minimized starting structure, shown for the whole complex, HcCoh, HcDoc, and $Ca^{2+}$ ions across five independent runs. The complex and HcCoh backbone were aligned to the reference structure prior to RMSD calculations, while HcDoc and $Ca^{2+}$ ions RMSD were calculated without prior alignment, reflecting relative interdomain mobility. (B) Per-residue backbone RMSF across five independent runs. The dashed vertical line separates HcDoc (residues 5-70) from HcCoh (residues 78-218); terminal residues were excluded from analysis.

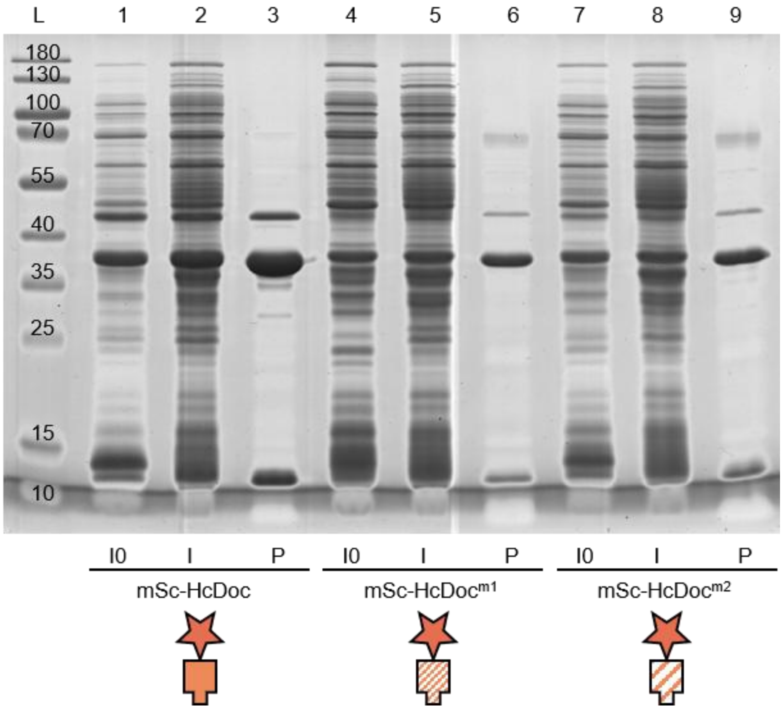


**Supplementary figure S6. Verification of expression and purification of mutant variants of mScarlet-I-HcDoc by sodium dodecyl sulfate polyacrylamide gel electrophoresis.** L is PageRuler Prestained Protein Ladder, 10 to 180 kDa; mScarlet-I-HcDoc (lanes 1-3), mScarlet-I-HcDoc$^{m1}$ (lanes 4-6), mScarlet-I-HcDoc$^{m2}$ (lanes 7-9). I0 is lysate of *E. coli* BL21-Gold (DE3) carrying pET21b+_*mScarlet-I-hcDoc-his,* pET21b+_*mScarlet-I-hcDoc$^{m1}$-his*, or pET21b+_*mScarlet-I-hcDoc$^{m2}$-his* harvested 2.5 hours after starting the cultivation, before induction. I is lysate of *E. coli* BL21-Gold (DE3) carrying pET21b+_*mScarlet-I-hcDoc-his,* pET21b+_*mScarlet-I-hcDoc$^{m1}$-his*, or pET21b+_*mScarlet-I-hcDoc$^{m2}$-his*, harvested 20 hours after inducing the production of the protein of interest. P is sample of purified mScarlet-I-HcDoc, mScarlet-I-HcDoc$^{m1}$, or mScarlet-I-HcDoc$^{m2}$ collected after the imidazole dilution. Theoretical molecular weight of the target proteins was 37.3 kDa. The two major bands in lines 3, 6, and 9 likely represent fully and partially denatured chimeric molecules of mScarlet-I linked to the thermophilic dockerin.

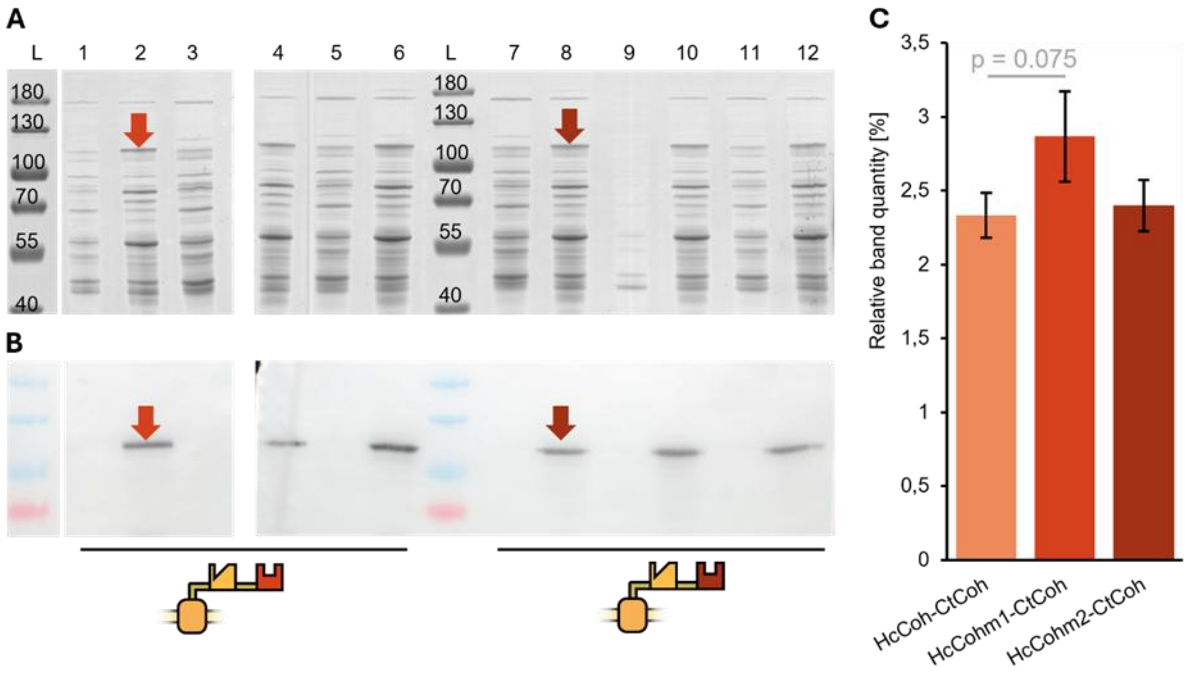


**Supplementary figure S7. Verification (A,B) and quantification (C) of expression of the HcCoh$^{m1}$-CtCoh and HcCoh$^{m2}$-CtCoh scaffoldins by sodium dodecyl sulfate polyacrylamide gel electrophoresis (A) and western blotting (B).** Lysates of *P. putida* EM371 carrying pSEVA238\_*ag43-hcCoh$^{m1}$-ctCoh-his* (HcCoh$^{m1}$-CtCoh; lanes 1-6) or pSEVA238\_*ag43-hcCoh$^{m2}$-ctCoh-his* (HcCoh$^{m2}$-CtCoh; lanes 7-12). Odd lanes contain samples collected 2.5 hours after the start of cultivation, before induction (I0 samples). Even lanes contain samples collected 4.5 hours after inducing the production of the protein of interest (I samples). L is PageRuler Prestained Protein Ladder, 10 to 180 kDa. The dark orange arrows in lanes 2 and 8 indicate the band representing the HcCoh$^{m1}$-CtCoh and HcCoh$^{m2}$-CtCoh scaffoldin, respectively (theoretical molecular weight = 93.5 kDa). The correct identification of the band representing the HcCoh-CtCoh scaffoldin variants was confirmed by western blotting using an anti-His-tag antibody. (C) The relative band quantity of bands representing the wildtype HcCoh-CtCoh scaffoldin (**Fig. S2**) and its mutant variants was averaged across three biological replicates and shown as average ± SD.

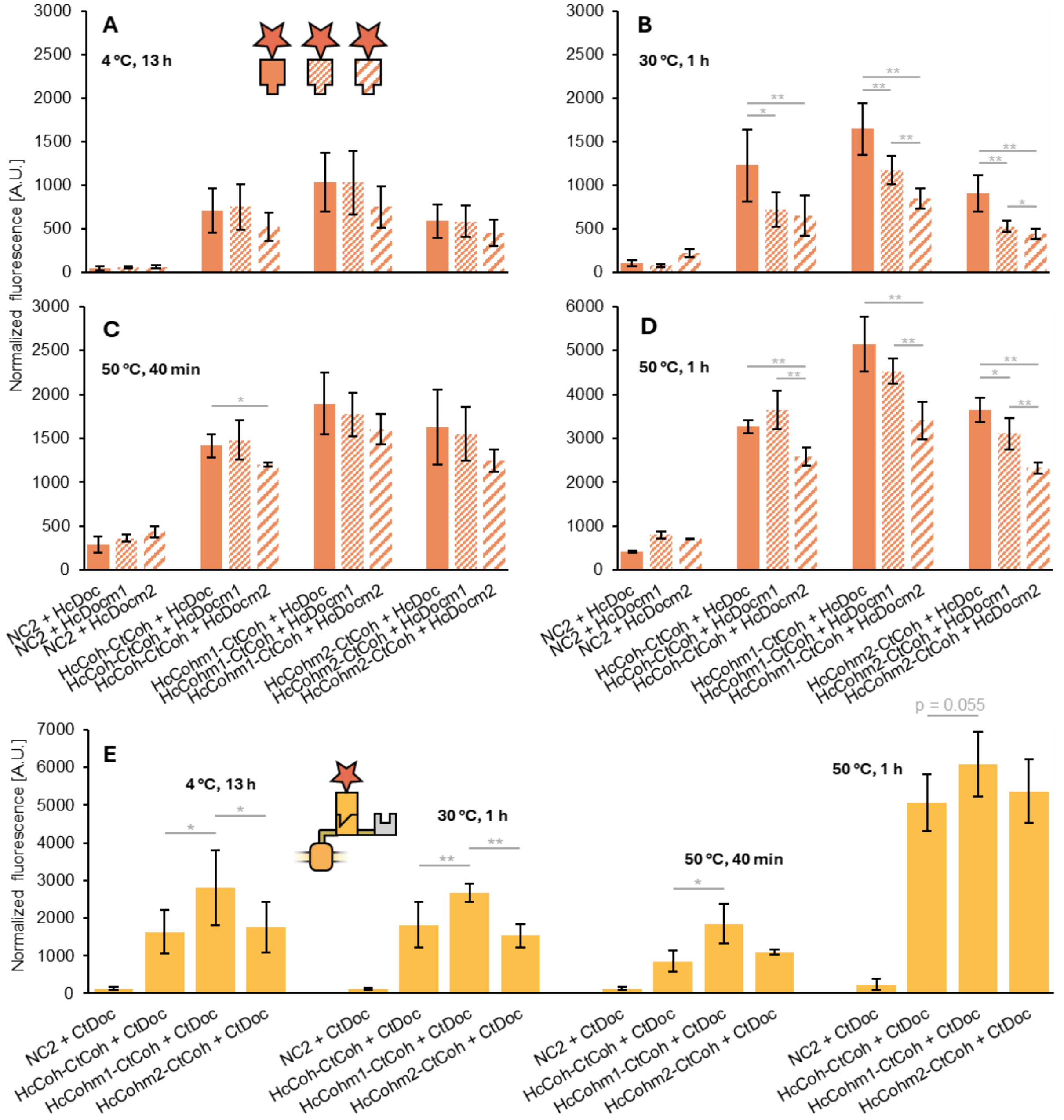


**Supplementary figure S8. Binding efficiency of mutant variants of the *A. clariflavus* (Hc) Coh-Doc pair under different experimental conditions.** Binding efficiency (represented by normalized fluorescence of mScarlet attached to the cell surface of *P. putida* EM371 via the Hc Coh-Doc or Ct Coh-Doc (E) interaction) after initiating the interaction at 4 °C for 13 h (A), at 30 °C for 1 h (B), at 50 °C for 1 h (C), and at 50 °C for 40 min (D). Data are shown as mean ± SD from three to four biological replicates. Asterisk denotes significance in the difference between two means at $p < 0.05$ (*) or at $p < 0.01$ (**).

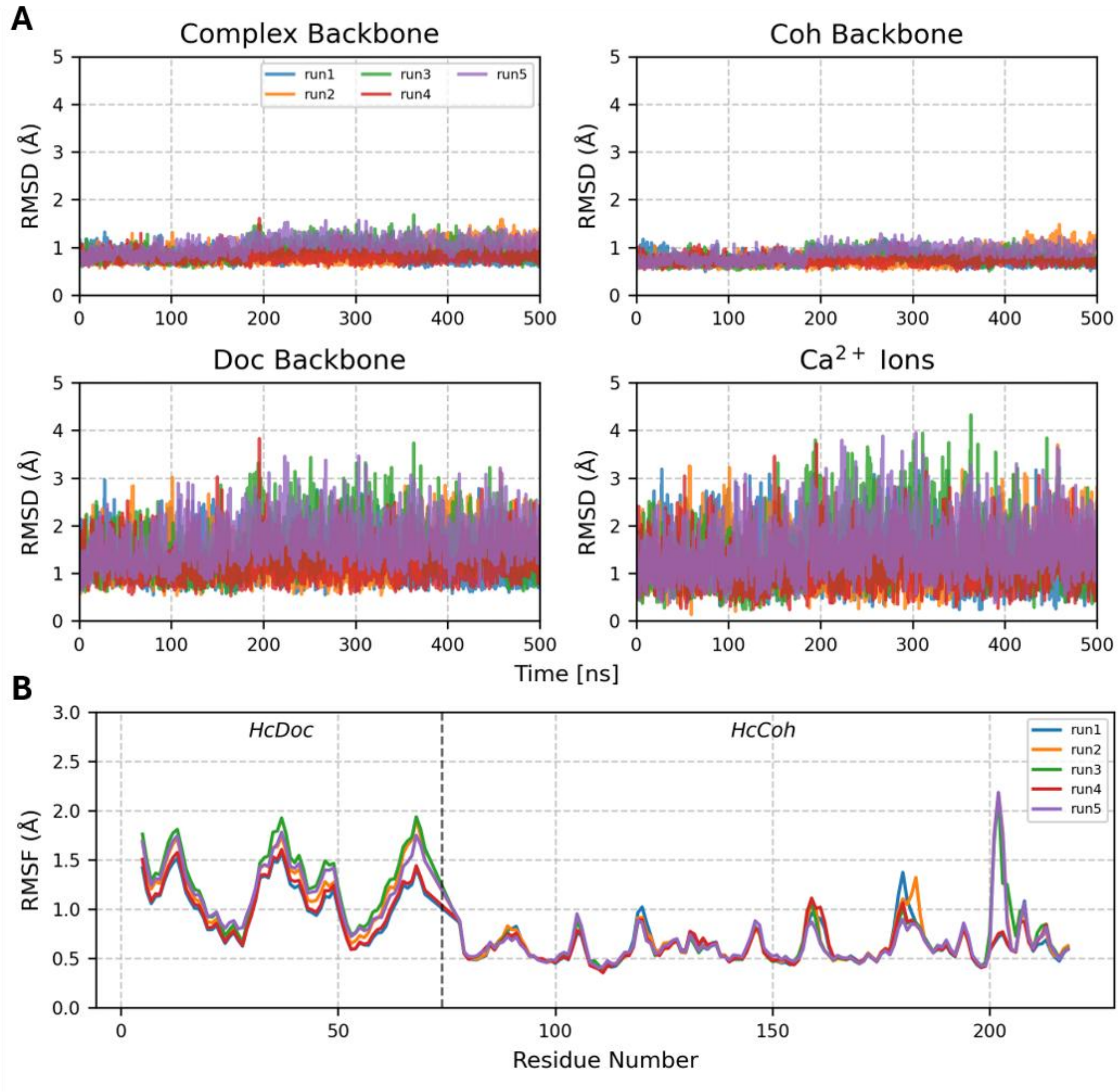


**Supplementary figure S9. MD simulations of the D46N HcCoh-HcDoc complex demonstrate stable and convergent trajectories.** (A) Backbone RMSD over 500 ns calculated relative to the energy-minimized starting structure, shown for the whole complex, HcCoh, HcDoc, and $Ca^{2+}$ ions across five independent runs. The complex and HcCoh backbone were aligned to the reference structure prior to RMSD calculations, while HcDoc and $Ca^{2+}$ ions RMSD were calculated without prior alignment, reflecting relative interdomain mobility. (B) Per-residue backbone RMSF across five independent runs. The dashed vertical line separates HcDoc (residues 5-70) from HcCoh (residues 78-218); terminal residues were excluded from analysis.

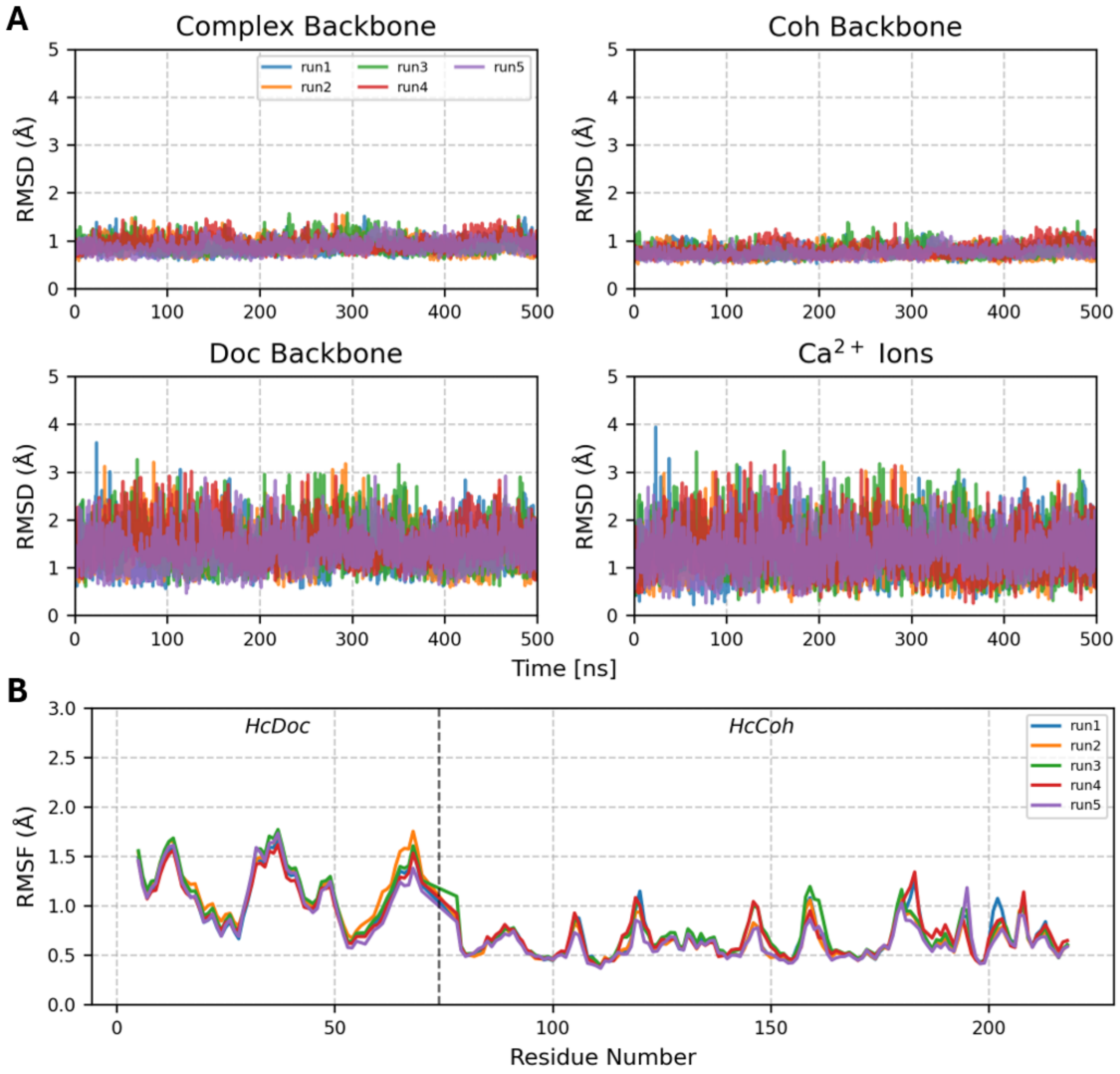


**Supplementary figure S10. MD simulations of the D48N HcCoh-HcDoc complex demonstrate stable and convergent trajectories.** (A) Backbone RMSD over 500 ns calculated relative to the energy-minimized starting structure, shown for the whole complex, HcCoh, HcDoc, and $Ca^{2+}$ ions across five independent runs. The complex and HcCoh backbone were aligned to the reference structure prior to RMSD calculations, while HcDoc and $Ca^{2+}$ ions RMSD were calculated without prior alignment, reflecting relative interdomain mobility. (B) Per-residue backbone RMSF across five independent runs. The dashed vertical line separates HcDoc (residues 5-70) from HcCoh (residues 78-218); terminal residues were excluded from analysis.

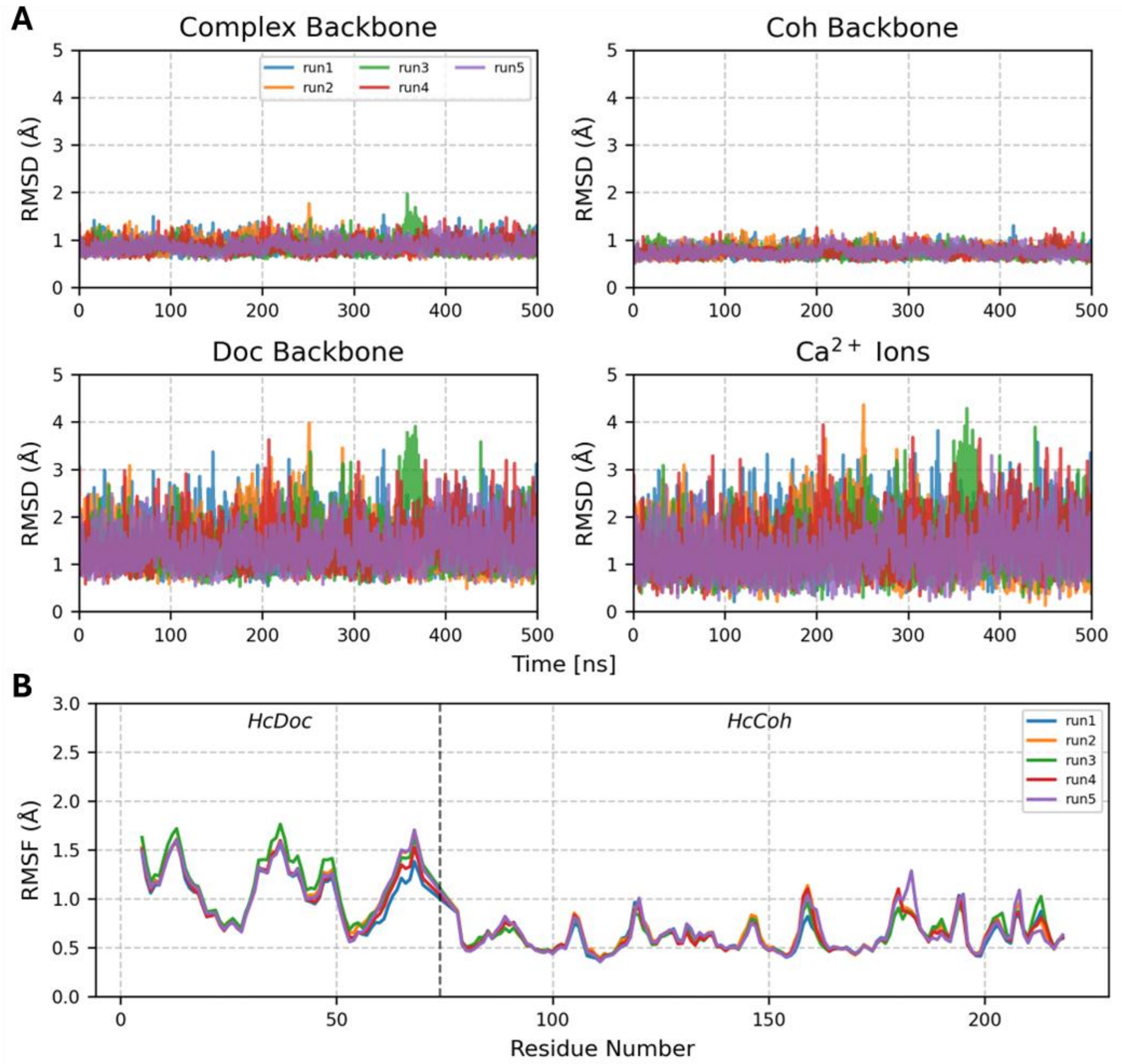


**Supplementary figure S11. MD simulations of the S37I/N39L HcCoh-HcDoc complex demonstrate stable and convergent trajectories.** (A) Backbone RMSD over 500 ns calculated relative to the energy-minimized starting structure, shown for the whole complex, HcCoh, HcDoc, and $Ca^{2+}$ ions across five independent runs. The complex and HcCoh backbone were aligned to the reference structure prior to RMSD calculations, while HcDoc and $Ca^{2+}$ ions RMSD were calculated without prior alignment, reflecting relative interdomain mobility. (B) Per-residue backbone RMSF across five independent runs. The dashed vertical line separates HcDoc (residues 5-70) from HcCoh (residues 78-218); terminal residues were excluded from analysis.

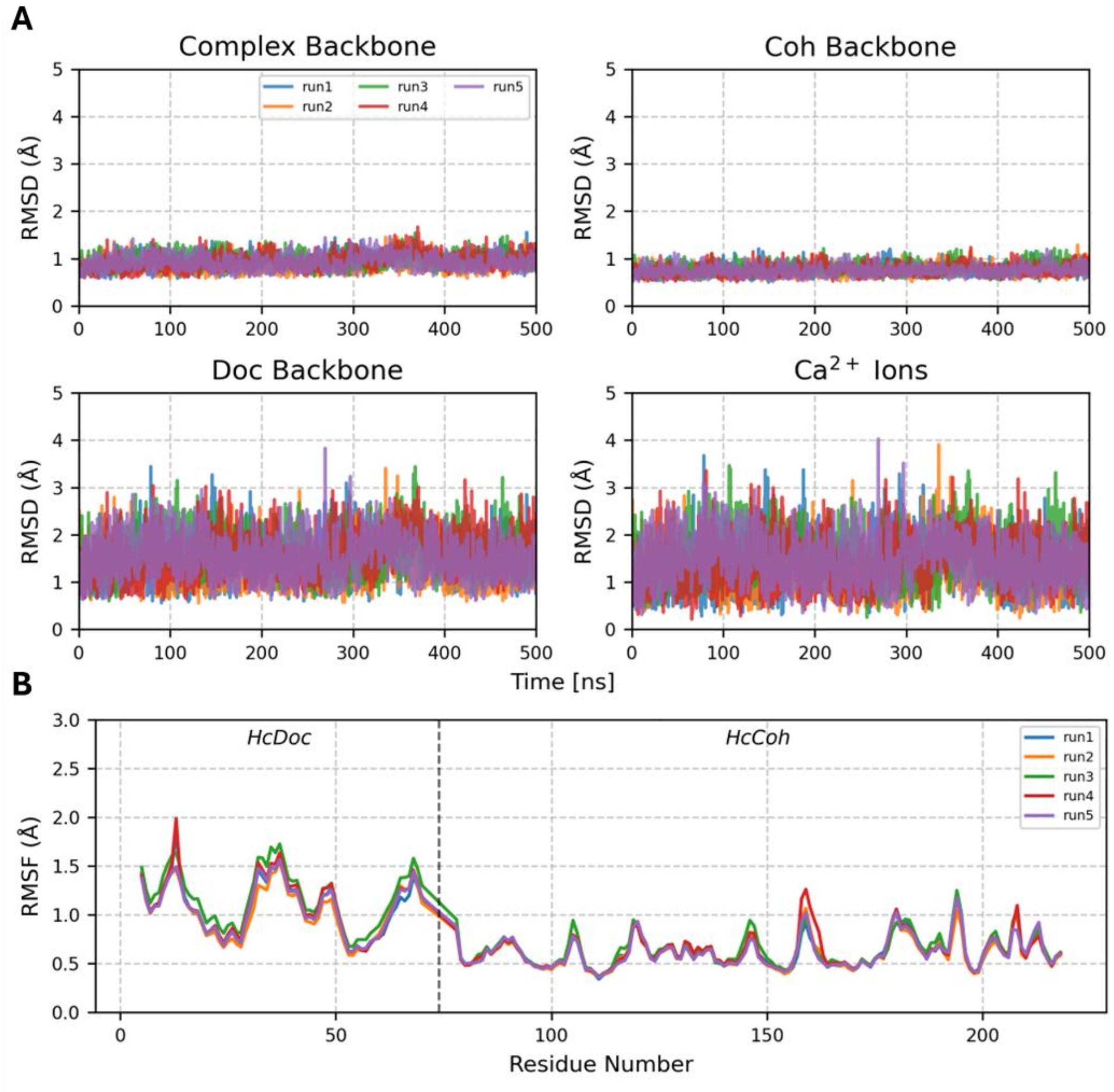


**Supplementary figure S12. MD simulations of the S37V/N39L HcCoh-HcDoc complex demonstrate stable and convergent trajectories.** (A) Backbone RMSD over 500 ns calculated relative to the energy-minimized starting structure, shown for the whole complex, HcCoh, HcDoc, and $Ca^{2+}$ ions across five independent runs. The complex and HcCoh backbone were aligned to the reference structure prior to RMSD calculations, while HcDoc and $Ca^{2+}$ ions RMSD were calculated without prior alignment, reflecting relative interdomain mobility. (B) Per-residue backbone RMSF across five independent runs. The dashed vertical line separates HcDoc (residues 5-70) from HcCoh (residues 78-218); terminal residues were excluded from analysis.

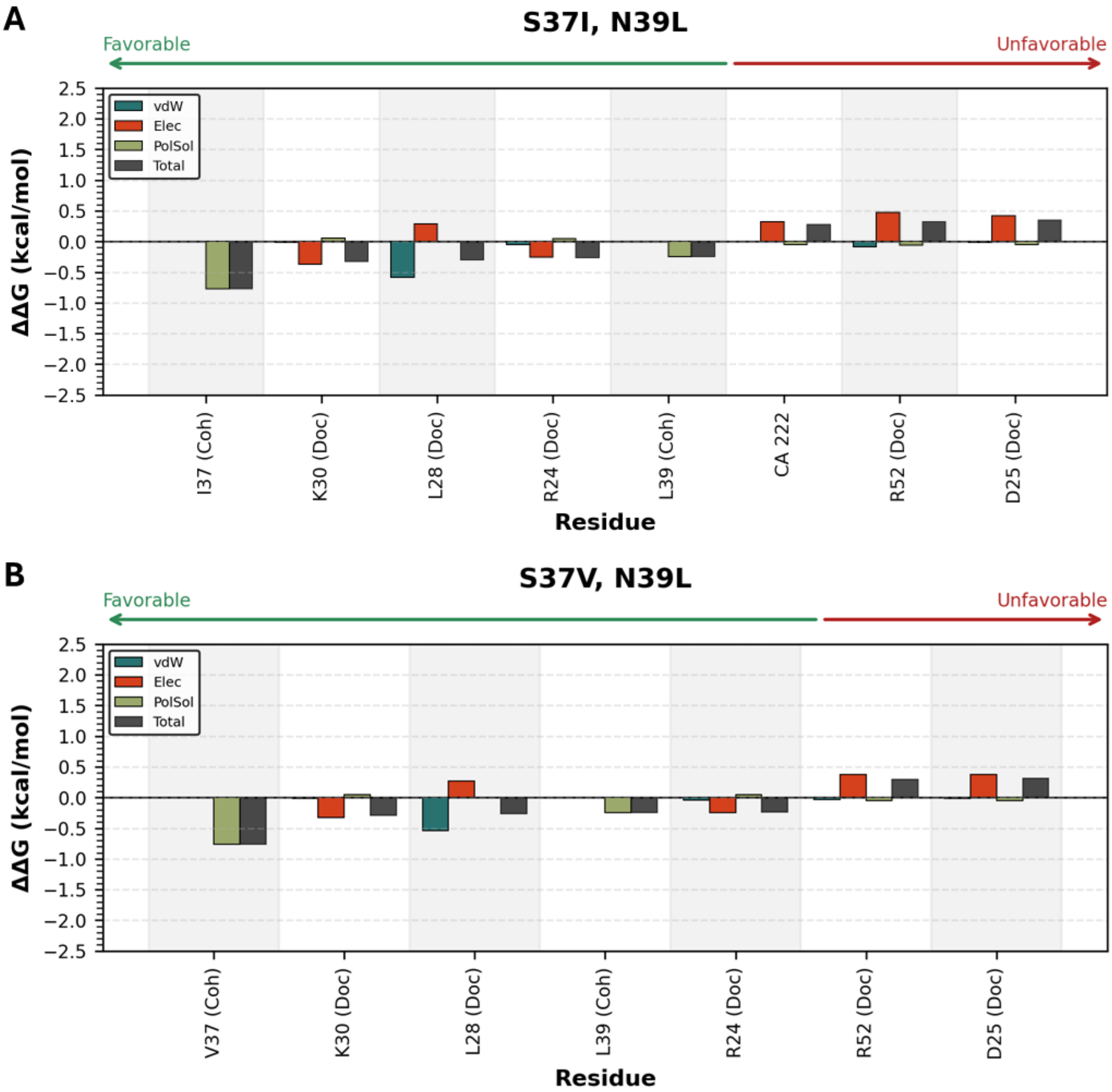


**Supplementary figure S13. Changes in pairwise interaction energy** between mutated HcCoh residues for (A) S37I/N39L and (B) S37V/N39L mutants. $\Delta\Delta G = \Delta G_{mut} - \Delta G_{WT}$, represents the change in pairwise interaction energy upon mutation, decomposed into van der Waals (vdW), electrostatic (Elec), and polar solvation (PolSol) terms and total (Total) contributions. Only residues with $|\Delta\Delta G| > 0.25$ kcal/mol are shown. Negative values indicate improved interactions relative to the wild type (more favorable); positive values indicate worsened interactions (unfavorable).

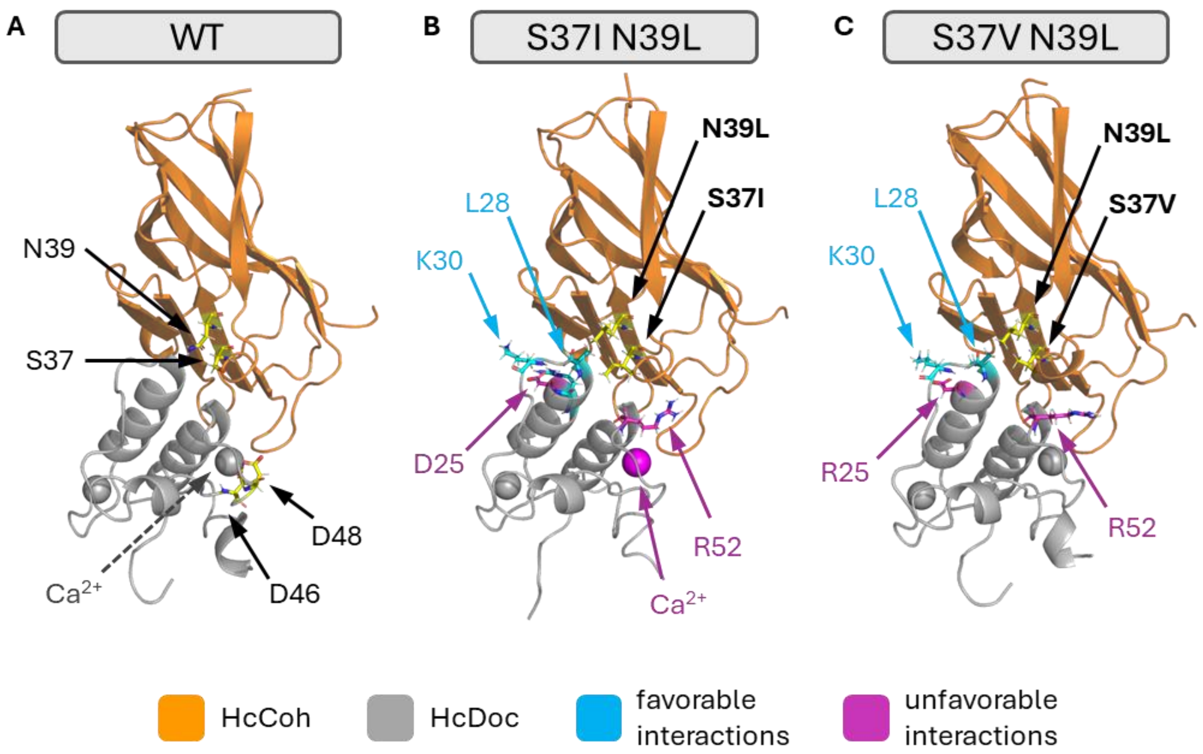


**Supplementary figure S14. Overview on mutation position and strongest changes in pair interactions with mutated residues.** A) Mutations and $Ca^{2+}$ ion positions on the HcCoh-HcDoc WT structure. Mutated residues are shown in yellow sticks with labeled black arrows, additionally with gray dashed line $Ca^{2+}$ ion, which is coordinated by D46 and D48, was also marked. Key altered favorable (in cyan) and unfavorable (in magenta) interactions in HcCoh mutants B) S37I/N39L and C) S37V/N39L were shown in sticks with labeled arrows.

### Supplementary sequences

**Supplementary sequence 1. DNA sequence encoding the Ag43-HcCoh-CtCoh chimeric protein.** Nucleotides encoding individual parts of the chimeric proteins are color-coded as follows: signal peptide – blue; multiple cloning site – green; HcCoh – orange, linker – pink; CtCoh – yellow; histidine tag – lavender; transmembrane domain of the Ag43 autotransporter domain – white.

SIGNAL PEPTIDE MULTIPLE CLONING SITE HCCOH LINKER CTCOH 6XHIS
MULTIPLE CLONING SITE TRANSMEMBRANE DOMAIN

ATGAAACGACATCTGAATACCTGCTACAGGCTGGTATGGAATCACATGACGGGCGCTTTCGT
GGTTGCCTCCGAACTGGCCCGCGCACGGGGTAAACGTGGCGGTGTGGCGGTTGCACCGTCTC
TTGCCGCAGTCACGTCACTCCCGGTGCTGGCTGCTGACAGATCTGCCATGGCGGAATTCGAG
CTCGGTACCCTCGAGGCCGGTCAGCTGCAAATCGACATCGGTCGCGTGAGCGGTGAACCGGG
TAGCATCGTGACCGTGCCGGTGACCTTCAGCAACGTGCCAGCCACCGGCATCTACGCCCTGA
GCCTGAACCTGAACTTCGACAGCACCAAGATCAGCGTGGTGAGCGTGGAACCAGGCTCGCTG
GTCGAGGACCCGGACGACTTCGCCCTGTTCACCAACAACGAGCACGGCTTCACCAGCATGAG
CTTCATGGCCCCAGCCGACCGCAGCCGCATCGTGGACAAGGATGGTGTGTTCGCCGCCATCA
AGTTCAAGATCTCGGAGGCGAACCCGGTGGGCGAAGTGTACGACATCAGCGTGAACTACAGC
CGCACCAGCTTCTACAGCACCGGCACCGAAGAGATCAAGAACGTGCTGTACAACGACGGTGC
CATCCTGGTGGGCAGCAACCCGACCCCGACTCAGTCGGCCACCCCGACGGTAACCCCGTCGG
CCACGGCCACGCCGACGCAATCCGCCACGCCGACCGTAACGCCCTCCGATGGCGTAGTAGTG
GAAATTGGCAAAGTGACGGGCAGCGTGGGCACCACCGTGGAAATTCCGGTGTACTTTCGCGG
CGTGCCCTCCAAAGGCATTGCCAATTGTGATTTTGTGTTTCGCTACGATCCGAATGTGTTGG
AAATTATTGGGATTGATCCCGGCGATATCATCGTGGACCCGAACCCGACCAAGAGCTTTGAT
ACCGCCATCTACCCGGATCGCAAGATCATCGTGTTCCTGTTCGCGGAAGACAGCGGCACCGG
CGCGTATGCCATTACCAAGGATGGCGTGTTCGCCAAAATCCGCGCCACCGTGAAATCGTCCG
CGCCGGGCTATATTACCTTTGACGAAGTGGGCGGCTTTGCCGATAATGACTTGGTGGAACAG
AAGGTCTCGTTCATCGACGGGGGGGTGAACGTGGGCAACGCCACCCACCATCACCACCATCA
CGGATCCATTGACCCCACGAATGTCACTCTCGCCTCCGGTGCCACCTGGAATATCCCCGATA
ACGCCACGGTGCAGTCGGTGGTGGATGACCTCAGCCATGCCGGACAGATTCATTTCACCTCC
ACCCGCACAGGGAAGTTCGTACCGGCAACCCTGAAAGTGAAAAACCTGAACGGACAGAATGG
CACCATCAGCCTGCGTGTACGCCCGGATATGGCACAGAACAATGCTGACAGACTGGTCATTG
ACGGCGGCAGGGCAACCGGAAAAACCATCCTGAACCTGGTGAACGCCGGCAACAGTGCGTCG
GGGCTGGCGACCAGCGGTAAGGGTATTCAGGTGGTGGAAGCCATTAACGGTGCCACCACGGA
GGAAGGGGCCTTTGTCCAGGGGAACAGGCTGCAGGCCGGTGCCTTTAACTACTCCCTCAACC
GGGACAGTGATGAGAGCTGGTATCTGCGCAGTGAAAATGCTTATCGTGCAGAAGTCCCCCTG
TATGCCTCCGTGCTGACACAGGCAATGGACTATGACCGGATTGTGGCAGGCTCCCGCAGCCA
TCAGACCGGTGTAAATGGTGAAAACAACAGCGTCCGTCTCAGCATTCAGGGCGGTCATCTCG
GTCACGATAACAATGGCGGTATTGCCCGTGGGGCCACGCCGGAAAGCAGCGGCAGCTATGGA

TTCGTCCGTCTGGAGGGTGACCTGATGAGAACAGAGGTTGCCGGTATGTCTGTGACCGCGGG
GGTATATGGTGCTGCTGGCCATTCTTCCGTTGATGTTAAGGATGATGACGGCTCCCGTGCCG
GCACGGTCCGGGATGATGCCGGCAGCCTGGGCGGATACCTGAATCTGGTACACACGTCCTCC
GGCCTGTGGGCTGACATTGTGGCACAGGGAACCCGCCACAGCATGAAAGCGTCATCGGACAA
TAACGACTTCCGCGCCCGGGGCTGGGGCTGGCTGGGCTCACTGGAAACCGGTCTGCCCTTCA
GTATCACTGACAACCTGATGCTGGAGCCACAACTGCAGTATACCTGGCAGGGACTTTCCCTG
GATGACGGTAAGGACAACGCCGGTTATGTGAAGTTCGGGCATGGCAGTGCACAACATGTGCG
TGCCGGTTTCCGTCTGGGCAGCCACAACGATATGACCTTTGGCGAAGGCACCTCATCCCGTG
CCCCCCTGCGTGACAGTGCAAAACACAGTGTGAGTGAATTACCGGTGAACTGGTGGGTACAG
CCTTCTGTTATCCGCACCTTCAGCTCCCGGGGAGATATGCGTGTGGGGACTTCCACTGCAGG
CAGCGGGATGACGTTCTCTCCCTCACAGAATGGCACATCACTGGACCTGCAGGCCGGACTGG
AAGCCCGTGTCCGGGAAAATATCACCCTGGGCGTTCAGGCCGGTTATGCCCACAGCGTCAGC
GGCAGCAGCGCTGAAGGGTATAACGGTCAGGCCACACTGAATGTGACCTTCTGA

**Supplementary sequence 2. DNA sequence encoding the mScarlet-HcDoc chimeric protein.** Nucleotides encoding individual parts of the chimeric proteins are color-coded as follows: mScarlet-I – pink; linker – gray; HcDoc – orange; histidine tag – lavender; stop codon – white.

MSCARLET-I LINKER HCDOC 6XHIS STOP

ATGGTGAGCAAGGGCGAGGCAGTGATCAAGGAGTTCATGCGGTTCAAGGTGCACATGGAGGG
CTCCATGAACGGCCACGAGTTCGAGATCGAGGGCGAGGGCGAGGGCCGCCCCTACGAGGGCA
CCCAGACCGCCAAGCTGAAGGTGACCAAGGGTGGCCCCCTGCCCTTCTCCTGGGACATCCTG
TCCCCTCAGTTCATGTACGGCTCCAGGGCCTTCATCAAGCACCCCGCCGACATCCCCGACTA
CTATAAGCAGTCCTTCCCCGAGGGCTTCAAGTGGGAGCGCGTGATGAACTTCGAGGACGGCG
GCGCCGTGACCGTGACCCAGGACACCTCCCTGGAGGACGGCACCCTGATCTACAAGGTGAAG
CTCCGCGGCACCAACTTCCCTCCTGACGGCCCCGTAATGCAGAAGAAGACAATGGGCTGGGA
AGCGTCCACCGAGCGGTTGTACCCCGAGGACGGCGTGCTGAAGGGCGACATTAAGATGGCCC
TGCGCCTGAAGGACGGCGGCCGCTACCTGGCGGACTTCAAGACCACCTACAAGGCCAAGAAG
CCCGTGCAGATGCCCGGCGCCTACAACGTCGACCGCAAGTTGGACATCACCTCCCACAACGA
GGACTACACCGTGGTGGAACAGTACGAACGCTCCGAGGGCCGCCACTCCACCGGCGGCATGG
ACGAGCTGTACAAGAACCCCAACCCGAACCCAACCCCGACCCCAACCCCAGGCAGCGGCAGC
CACAAGTTCATCTACGGCGACGTGGACGGCAACGAGAGCGTGCGCATCAACGACGCCGTGCT
GGTGCGCGACTACGTGCTGGGCAAGATCGACGAGTTCCCGTACGAGTACGGTATGCTGGCCG
CCGACGTGGATGGCGACGGCAACATCCGTATCAACGACAGCGTCCTGATCCGCGATTTCGTC
CTGGGCAAAATCTCGCTGTTCCCGGTGGAAGAACAACTCGAGCACCACCACCACCACCACTG
A

**Supplementary sequence 3. DNA sequence encoding the mScarlet-CtDoc chimeric protein.** Nucleotides encoding individual parts of the chimeric proteins are color-coded as follows: mScarlet-I – pink; linker – gray; CtDoc – yellow; histidine tag – lavender; stop codon – white.

MSCARLET-I LINKER CTDOC 6XHIS STOP

```
ATGGTGAGCAAGGGCGAGGCAGTGATCAAGGAGTTCATGCGGTTCAAGGTGCACATGGAGGG
CTCCATGAACGGCCACGAGTTCGAGATCGAGGGCGAGGGCGAGGGCCGCCCCTACGAGGGCA
CCCAGACCGCCAAGCTGAAGGTGACCAAGGGTGGCCCCCTGCCCTTCTCCTGGGACATCCTG
TCCCCTCAGTTCATGTACGGCTCCAGGGCCTTCATCAAGCACCCCGCCGACATCCCCGACTA
CTATAAGCAGTCCTTCCCCGAGGGCTTCAAGTGGGAGCGCGTGATGAACTTCGAGGACGGCG
GCGCCGTGACCGTGACCCAGGACACCTCCCTGGAGGACGGCACCCTGATCTACAAGGTGAAG
CTCCGCGGCACCAACTTCCCTCCTGACGGCCCCGTAATGCAGAAGAAGACAATGGGCTGGGA
AGCGTCCACCGAGCGGTTGTACCCCGAGGACGGCGTGCTGAAGGGCGACATTAAGATGGCCC
TGCGCCTGAAGGACGGCGGCCGCTACCTGGCGGACTTCAAGACCACCTACAAGGCCAAGAAG
CCCGTGCAGATGCCCGGCGCCTACAACGTCGACCGCAAGTTGGACATCACCTCCCACAACGA
GGACTACACCGTGGTGGAACAGTACGAACGCTCCGAGGGCCGCCACTCCACCGGCGGCATGG
ACGAGCTGTACAAGAACCCCAACCCGAACCCAACCCCGACCCCAACCCCAGGCAGCGGCGGG
ACGCCCAGCACCAAACTGTACGGCGATGTCAATGATGACGGCAAAGTGAACTCGACCGACGC
CGTCGCCCTGAAGCGGTATGTGCTGCGGTCGGGCATCAGCATCAACACCGACAATGCCGACC
TGAATGAAGACGGCCGGGTCAATTCGACCGACCTGGGCATTCTGAAGCGCTATATTCTCAAA
GAAATTGACACGCTGCCGTACAAGAACCACCACCATCACCACCACTAA
```
